# Concurrent de novo inference of cell types and gene regulatory networks with NetMap

**DOI:** 10.64898/2026.10.04.756614

**Authors:** Anne Hartebrodt, Mhaned Oubounyt, Gustavo P. de Almeida, Timo Reim, Nina Wenke, Jan Baumbach, Dietmar Zehn, David B. Blumenthal

## Abstract

Existing methods for GRN inference based on single-cell RNA-sequencing (scRNA-seq) data typically require prior cell type calling and then infer GRNs at cell type resolution. This risks propagation of errors from cell annotation to GRN inference. Moreover, it makes existing methods blind to regulatory programs that are shared between different cell types, as well as to subtle differences in regulatory activity between cells of the same type. To address these limitations, we present NetMap, a pipeline that jointly infers cell-level GRNs and clusters cells based on shared regulatory programs, combining state-of-the-art single-cell autoencoders and explainable artificial intelligence methods. NetMap outperforms existing methods on synthetic data with a known ground-truth GRN and is able to reliably detect biologically plausible regulons on real data. Moreover, requiring only a small set of user-provided marker genes, NetMap can be used for GRN structure-informed probabilistic cell type assignment. We demonstrate the utility of this approach using PBMC data, where we achieve near perfect clustering and cell type recovery performance compared with manually created cell type labels. Finally, we applied NetMap to a data set with naïve, activated and exhausted T-cells, and were able to verify that NetMap retrieves relevant T-cell state signatures from the data, with two approaches. NetMap can be installed as a python package from https://github.com/bionetslab/netmap and is compatible with the scverse system. Scripts to reproduce our results can be found at https://github.com/bionetslab/netmap-evaluation.

## 1 Introduction

Gene regulatory networks (GRNs) [1] are graphical representations of complex regulatory programs in the cell. A regulatory relationship between regulators, most often transcription factors (TFs), and their target genes is represented as a direct edge between the two nodes. Gene regulatory networks are dynamic and context specific; for example, two cell types are expected to have two distinct GRNs describing the different regulatory programs taking place [2]. Standard methods to detect such cell type specific GRNs first cluster the highly variable genes, identify cell types of interest based on marker genes, and then process each cluster individually by using a GRN inference method on each homogeneous cluster separately [3, 4]. These methods produce a single GRN for each cluster, disregarding dynamic cell-level changes of regulatory activity. Moreover, there is the possibility of propagating clustering and cell annotation errors to the GRN inference, as any misassignment of cells to clusters will bias the regulatory relationships inferred for the affected cell types [5].

In this work, we present a method called NetMap for producing GRN-based embeddings for every cell rather than creating networks at the cluster level. This allows us to keep the cell resolution throughout the analysis and interact with the GRN much like one would interact with the raw single-cell expression object. NetMap is based on an autoencoder framework as used in [6, 7] coupled with explainable AI methods [8], followed by clustering and downstream processing. The key distinction from standard GRN inference approaches is that NetMap performs GRN inference at the single-cell level prior to any clustering step, thereby bypassing the need for predefined cell type labels altogether. Clustering and downstream analysis are then carried out on gene regulatory signal at rather than on the raw gene data. This avoids the propagation of clustering and cell annotation biases into the GRN inference step and entails that cell types uncovered by NetMap inherently correspond to shared regulatory programs rather than shared marker gene expression.

The inference of single-cell GRNs is an unresolved research field. The conceptually closest method is scGeneRAI [9], which uses masking autoencoders coupled with layerwise relevance propagation (LRP) to obtain attribution scores relating each input gene to the output (target) gene. The output of scGeneRAI is a cell-by-edge object which can be further refined by selecting high ranking edges. The single-cell foundation model LINGER [10] leverages prior information with a pretraining stage on bulk expression and regulatory data from ENCODE repository and a refinement stage using single-cell multiome data. In the inference phase, LINGER uses Shapley values to predict the importance of input features for the targets at cell type level. Cell-level GRNs can then be derived post hoc by reweighting the population-level GRN with each cell’s gene expression and chromatin accessibility. They thus share a common topology and differ only in edge weights. scPRINT [11] is another foundation model which uses ESM-2 embeddings, genomic location, and the gene expression of atlas-scale data to pretrain a model using several pretraining objectives. At inference time, the attention matrices of the transformer are extracted to generate GRNs at the resolutions of cell types, and — according to the authors’ evaluation — outperforms other foundation models (scGPT [12], preprint version of Geneformer *v2* [13]) that had previously been proposed for GRN inference in this downstream task. Moreover, classical GRN inference methods originally designed for bulk gene expression data can be used for cell type-level GRN inference by subsetting the input to the cells of the target cell type. For this work, we include GRNBoost2 [4] as such a baseline.

NetMap uses an autoencoder to embed the data into a low dimensional space, and then uses GradientShap to trace back the informative features for each output node individually. This results in a fully connected GRN, which can then be filtered to remove irrelevant edges. The filtered network is then processed to find cluster-specific regulons, and apply knowledge-based filtering steps, such as mapping known transcriptional networks. Additionally, the GRNs can be used directly to obtain probabilistic cell type labels, by running the PageRank algorithm on the cell level graphs. Hence, it is possible to bypass standard cell type annotation completely and directly use NetMap without prior labeling. We also include a PageRank-based gene scoring that aggregates the most relevant genes to a cell type GRN level. This allows us to show commonalities between cell types rather than focusing only on the differences. Further strengths of the NetMap framework include fast training an inference and convenient interfaces for downstream processing of cell-level GRNs. NetMap relies solely on scRNA-seq data and hence can be used on any dataset, without inducing a strong bias due to prior knowledge incorporation. Functions of the scverse [14] ecosystem, such as statistical testing and plotting, can directly be applied to the GRN. Additionally, we provide several network visualizations, enabling the user to investigate the dynamic changes in the networks.

## 2 Results

### 2.1 Overview of the NetMap method

NetMap is built on a shallow bottleneck autoencoder that reconstructs raw count data by optimizing the parameters of a negative binomial (NB) or zero-inflated negative binomial (ZINB) distribution, accommodating the overdispersion and excess zeros characteristic of scRNA-seq data [15]. The features of gene expression data are usually highly correlated; therefore, multiple models are trained and added to an ensemble. For each model, we use GradientShap to identify the most informative input genes for each target gene and average the predictions to obtain robust cell-level GRNs. This is illustrated in Figure 1a. An in-depth description of the autoencoder architecture and training is provided in Section 4.1.

**Fig. 1:**
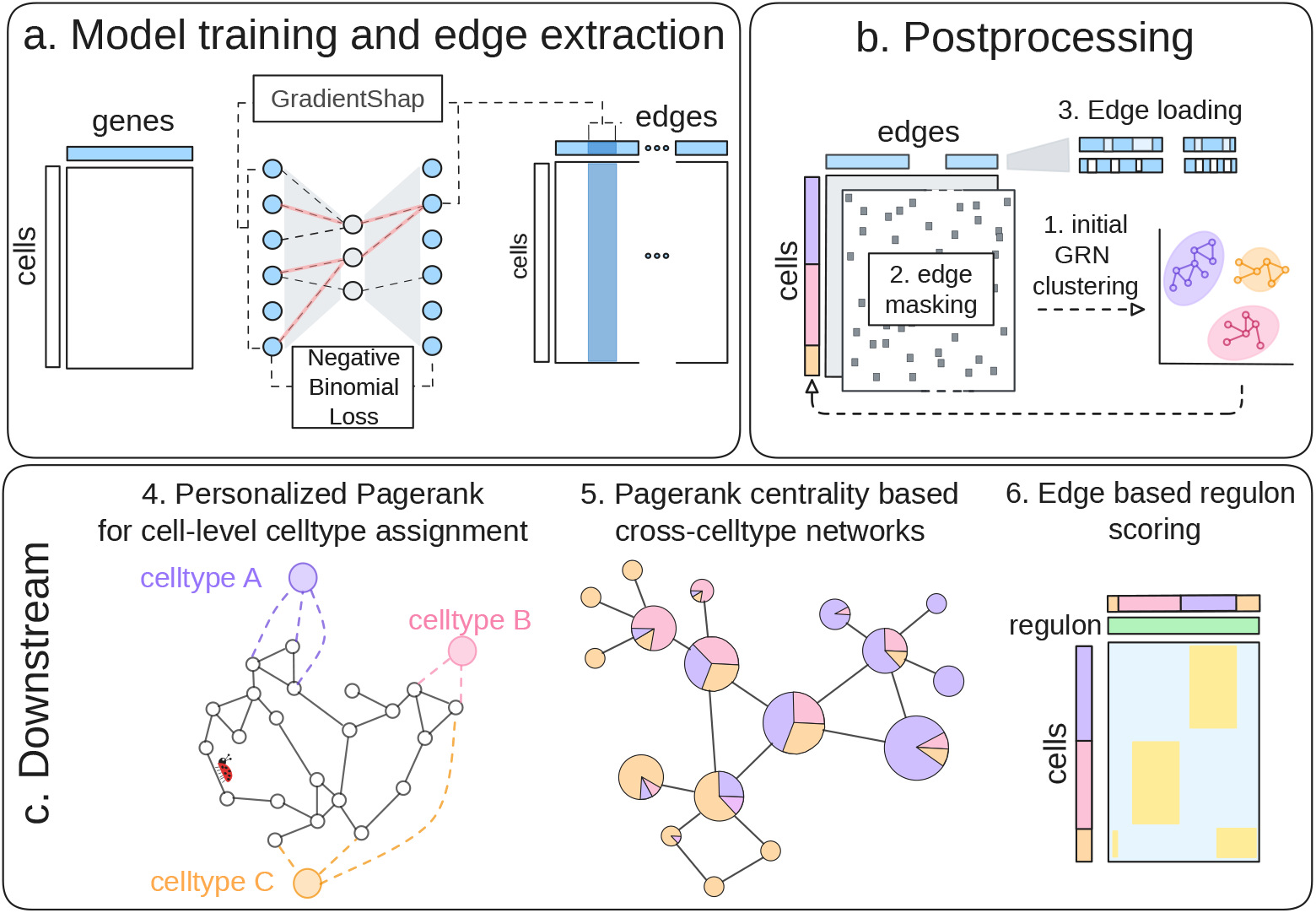
Overview over the NetMap pipeline. (a) A bottleneck autoencoder is trained and GradientShap is used to obtain edges for every source and target for a user defined number of models. (b) 1. The basic preprocessing includes clustering of the top *k*% edges; 2. edge masking based on the gene expression, and 3. loading the top most important edges per cluster. (c) Downstream analysis; 4. cell states can be assigned using personalized PageRank; 5. the central genes can be extracted and 6. regulons can be extracted per cluster and source gene.

The raw output, a dense cell-level GRN data frame, is subsequently processed through a pipeline consisting of clustering, sparsification, edge selection, and regulon construction. We apply Leiden clustering directly to the GRN, using the top 10% of edges with the highest global weights, analogous to the selection of highly variable genes in Scanpy (Figure 1b). In the sparsification step, we apply a missingness mask which suppresses edges which are unlikely to be biologically active in this cell: If gene A and gene B are both not expressed in the cell, the edge is unlikely to be present. This step suppresses spurious interactions driven solely by imputation from other cells and cell populations. This step, does not require the clustering as an input. A more detailed description of the masking step can be found in Section 4.2.1. Optionally, an external GRN can be used to mark potentially relevant edges in the edge object (Section 4.2.2). Then, the top edges are discarded and the interesting edges are loaded, which are the top scoring edges per cluster (Figure 1b, details in Section 4.2.3).

At this point it is still not necessary to annotate the clusters, as this step just selects cluster-wise informative edges. However, optionally, a known cell type annotation can be transferred between the the gene expression object metadata and the GRN object using an in-built label transfer functionality described in Section 4.5.3. As a more advanced alternative which takes the structure of the GRN into account, cell types can be assigned using a customized version of the PageRank algorithm, which allows probabilistic assignment of the cell identities and can also be used to score arbitrary gene sets. The PageRank algorithm is additionally used to retrieve important genes in the network by their centrality. This step is described in Section 4.3.1 and illustrated in Figure 1c.

Finally, as described in Section 4.3.2, regulon construction is performed by treating each source gene as a potential regulator and selecting its top-ranked target genes, yielding cluster- and gene-specific regulons. A combined regulon score is computed as the mean of the constituent edge weights. Customized visualization functions are described in Section 4.3.3. NetMap’s final output consists of two objects: a full GRN object, which can be flexibly filtered and visualized, and a signature object providing an aggregated, regulon-level summary of regulatory activity on a cluster level (Figure 1c).

### 2.2 NetMap recovers ground-truth clusters and GRNs in simulated and real data

We first evaluated NetMap using synthetic, realistically sized single-cell gene expression datasets generated using scMultiSim [16], in which each cluster was assigned its own specific GRN. In this novel benchmarking strategy, we induced cluster-specific GRNs into the simulated data, such that each cluster possesses a characteristic network to be recovered (on-target edges). Conversely, for a cluster *c*_*i*_, edges belonging to the GRNs of other clusters *c*_*k*_ with *k ≠ i*, should ideally not be recovered (off-target edges). The data generation is described in detail in Section 4.5.1.

We then performed a hyperparameter search to find the best performing NetMap configuration. For the parameter sweep, we recorded the percentage recovered edges (recall), compared the clustering performance to the groundtruth clustering, and we measured the runtime of the method (details in the Supplementary Material). The best performing method on synthetic data was a shallow autoencoder (zero-inflated, or standard negative binomial) with a hidden layer size of 64, with a latent dimensionality of 8, using 0.1 dropout, coupled with GradientShap with a background vector of zeros. We compared NetMap to scGeneRAI, a conceptually close competitor using layer-wise-relevance propagation. Like NetMap, scGeneRAI can be used for *de novo* clustering and GRN inference, i. e., it can jointly infer both the clustering and the GRN. Foundation models were excluded from this initial evaluation, as the pretraining negates the possibility to use synthetic data. As a naïve baseline, we compared NetMap to GRNBoost2 using the ground-truth labels as the clustering information, thus making the inference task easier for GRNBoost2. Results with the best performing configuration of GRNBoost2 are shown in Figure 2a-d. These results have been computed on newly generated synthetic data which were not used for the hyperparameter tuning.

**Fig. 2:**
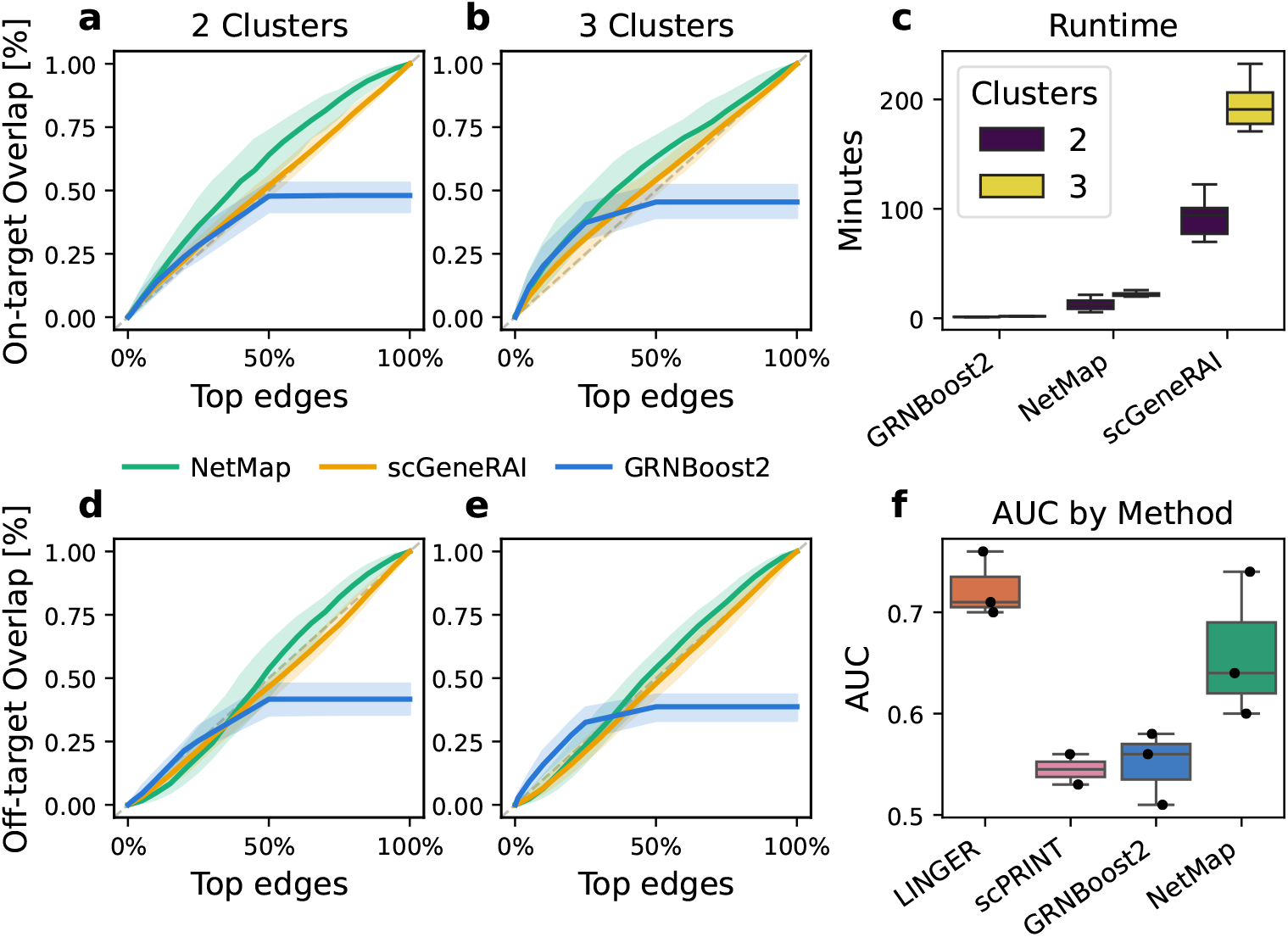
Results of the evaluation of NetMap on synthetic and real-world data. (a,b) On-target recovery of edges for cluster specific GRNs with 2 and 3 simulated clusters respectively. On-target edges are recovered for the cluster of interest and should be high. The dashed gray line corresponds to random performance. (d, e) Off-target recovery of edges of a GRN belonging to a different cluster. The recovery of off-target edges should be low. (c) Runtime of the method in minutes. A cluster leads to increase in number of genes (+ 60-150 genes) and number of cells (+500 cells). (f) Comparison of the foundation models LINGER and scPRINT and the baseline GRNBoost2 with NetMap on real scRNA-seq data.

NetMap outperforms both scGeneRAI and GRNBoost2 in terms of on-target recall, as shown in Figure 2a and b. NetMap achieves near-random performance for the off-target edges which, is a desirable result (Figure 2d and e). The precision is overall low across all methods, a known limitation when benchmarking on realistically sized synthetic datasets with many genes [1]. Clusterings were retrieved perfectly for all synthetic benchmarks, with a score of 1.0, indicating the strong effect of the induced clusters in the synthetic data. There was no clear trend for the noiseless and noisy datasets, so the results shown are averaged by the number of clusters in the dataset (see Figure S4). In terms of runtime, NetMap is significantly faster than scGeneRAI, and although slower than GRNBoost2 operates within practically feasible runtimes, as shown in Figure 2c. The xAI methods scale quadratically with the number of genes, meaning scGeneRAI becomes impractical with standard sized single-cell expression data. NetMap has a significantly lower runtime footprint and hence is applicable to realistically sized datasets.

To evaluate NetMap on real data, we benchmarked it against the state-of-the-art foundation models LINGER and scPRINT [11] as well as the GRNBoost2 baseline, using a publicly available peripheral blood mononuclear cell (PBMC) dataset from 10X Genomics, which provides paired scRNA-seq and scATAC-seq measurements from the same cells [17]. scGeneRAI was excluded due to its huge runtime requirements based on the synthetic benchmark (Figure 2c). We also excluded the single-cell mode of LINGER, as an initial test run on 100 cells revealed a runtime of 187 minutes. In this mode, LINGER requires 1-2 minutes per cell, plus additional steps before and after the cell network construction, making it unsuitable for realistic applications. We constructed three evaluation cases (Table 3), each consisting of two distinct cell types, where one corresponds to a population with experimentally validated regulatory interactions. Predicted transcription factor-target interactions inferred by each method were compared against ChIP-seq binding evidence as an independent ground-truth reference.

A critical methodological distinction separates NetMap from all competing approaches: LINGER, GRNBoost2, and scPRINT all require cell type clustering as a prerequisite, meaning they operate on pre-filtered, homogeneous cell populations assigned to the target cell type. NetMap, by contrast, infers per-cell GRNs directly from the full mixed dataset without any prior clustering. Averaged across cells, these per-cell GRNs recover cell type structure at the level of gene interactions with a clustering score of 0.99 or higher as shown in Table 1, demonstrating that NetMap captures cell type identity through regulatory structure alone.

**Table 1:** Clustering scores for application cases. NetMap achieved almost perfect clustering scores in all three application cases.

| Case | Cell Type 1 | Cell Type 2 | Evaluated TF | Clustering score |
| --- | --- | --- | --- | --- |
| Case 1 | Classical monocytes | Memory CD4 T-cells | IRF4 | 1.0 |
| Case 2 | Intermediate monocytes | Naïve CD4 T-cells | SPI1 | 0.99 |
| Case 3 | Intermediate monocytes | Naïve CD4 T-cells | FOXP3 | 0.99 |

GRN recovery performance against the ChIP-seq-based ground truth was evaluated using AUC which is shown in Figure 2f. Overall, LINGER achieved the highest median AUC of 0.71 across the three evaluation cases. However, this result reflects the considerable advantages LINGER operates under: pre-clustered cell type-specific input, chromatin accessibility signals, and regulatory element priors that provide direct evidence of transcription factor binding. Chromatin accessibility is generally less available than single-cell RNA-seq data, rendering LINGER somewhat less broadly applicable. Despite relying solely on gene expression data from unclustered mixed populations, without chromatin accessibility, regulatory priors, or cell type annotations, NetMap achieved a median AUC of 0.65, outperforming GRNBoost2 and scPRINT, both of which also benefit from pre-clustered input yet still trail NetMap in performance.

### 2.3 NetMap uncovers stable and biologically plausible regulons in PBMC data

To demonstrate the general applicability of our approach, as well as the utility of our downstream refinements, we applied the NetMap method to a set of four PBMC datasets sequenced in two replicates with two sequencing technologies, 10x and BD Rhapsody (BD Biosciences, Franklin Lakes, NJ). We ran a preprocessing pipeline with standard quality control steps, including doublet removal, and filtered genes based on their expression across annotated cell types to retain genes with sufficiently high coverage for reliable model fitting. We ran the NetMap inference and postprocessing pipeline with default parameters and extracted the clusterings and regulons de-novo using the raw GRN objects. In the remainder of the manuscript the replicates will be named BD-1, BD-2, 10X-1, and 10X-2. The application of NetMap on these datasets resulted in four signature objects obtained for the same underlying biology. These objects contain the averaged weights of all edges belonging to the regulons obtained from selecting the top targets per source gene for each cluster, while removing non unique edges.

In Figure 3a and b we show the ground truth labeling done on the gene expression data and the GRN clustering with the transferred labeling respectively. Clusters were matched to the annotation derived from the gene expression data using a built-in maximum cardinality matching method, and clustering concordance was assessed by computing the clustering score reflecting the fraction of cells which were assigned the same labels in the GRN object and in the GEX object. Overall, the match was very close with mainly a few closely related cells like CD4+ and CD8+ T-cells switching cluster assignment. We then proceeded to perform GRN-based cell labeling using NetMap’s built-in PageRank method. In Figure 3c, we show a qualitative plot with the assignment of the cells to a category. Most cells remain identically assigned, however, some natural killer (NK) cells got labeled as CD8+ T-cells. Given their shared program of cytotoxicity, NetMap likely picks up on a shared regulatory signature between NK and CD8+ T cells. Figure 3d shows the averaged assignments between the ground truth and the PageRank predictions across the four datasets, indicating that the cells are generally assigned to the same category. In Figure S5, a similar plot showing the concordance of the GRN Leiden clustering with the GRN clustering indicates an even higher degree of similarity between the clusterings, mismatching a few CD8+ and CD4+ cells where the annotation is more granular and the cell identities become less well defined.

**Fig. 3:**
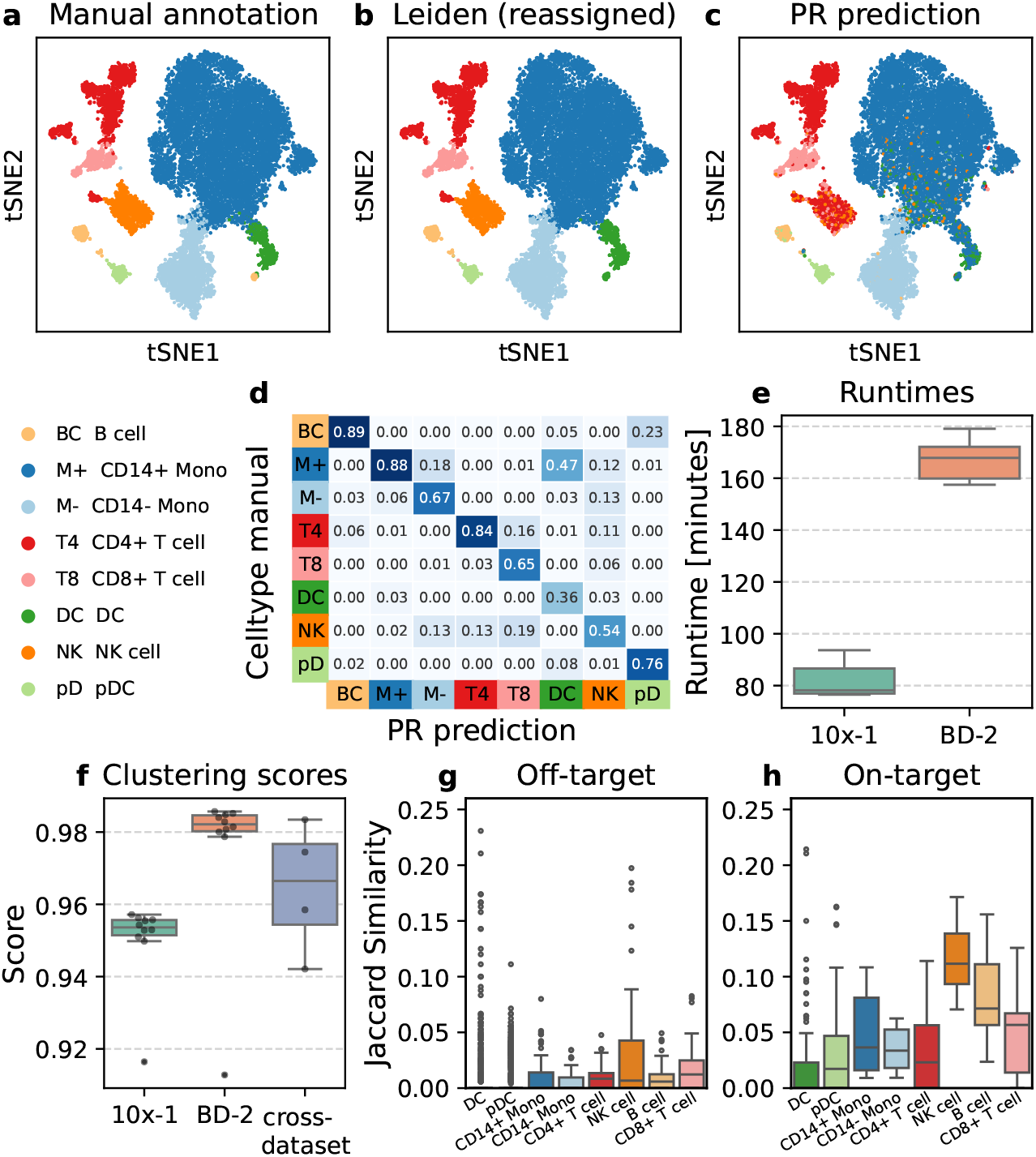
Application of NetMap on PBMC datasets. (a, b, c) tSNE plots showing the manual GEX labeling for BD-2 (a), the transferred labeling for BD-2 (b), and the PageRank prediction of the cell types of BD-2 (c). (d) Heatmap of the averaged and column normalized PR assignments over the four datasets. (e) Runtimes for 10x-1 (7K cells, 1K genes) and BD-2 (15K cells, 1K genes) repeat runs with different seeds. (f) Clustering scores for 10x-1 and BD-2 repeat runs with different seeds, as well as clustering scores for the four datasets. (g, h) Marker gene overlaps for off-target and on-target (cell type definition matches marker gene set label) top ranking regulons extracted from all four datasets.

The runtime at roughly 1000 genes (10x-1: 7K cells, 1K genes, BD-2 15K cells, 1K genes) is stable across repeats and depends largely on the number of cells in the dataset as they are relatively equal (Figure 3e). The clustering scores are also reproducible across replications of the same experiment with different seeds and across the technical replicates as shown in Figure 3f, where the left and middle box show the clustering coefficients obtained by running NetMap on the 10x-1 and one of the BD-2 runs 10 times respectively, and the right box shows the clustering scores obtained for the four replicates, with one run per replicate. NetMap mapped over 95% of the cells identically to the gene expression based clustering, and the mappings are consistent across multiple runs, with a slightly better score achieved on average for BD-2. Overall, the clustering scores are satisfying across different technical replicates and across independent computational runs.

To highlight the biological relevance of the recovered regulons, we assessed the overlap between publicly available marker gene data sets from PanglaoDB [18] and CellMarker [19] and the top regulons extracted for each cluster. Reference marker sets were matched as closely as possible to the manual cell type annotations. We then computed the Jaccard overlap for each cluster derived high ranking gene sets with all marker sets. In Figure 3g and h, we show the overlap of the off-target marker set (where the marker sets and cell type are not the same) and the overlaps of the marker sets with the closest match (where the cell type label matches the cell type marker set). Generally, the overlap between matching sets is higher, indicating that the clusters we obtain respect the known biology. It is to be noted that the definitions of the cell types in the PanglaoDB and CellMarker databases are literature-derived and not standardized, leading to high disagreements between publications. See the Figures S6 and S7 for for clustered heatmaps showing the lack of agreement between cell type definitions and Table S2 for our attempt at matching the literature-derived definitions to high level cell type marker sets based on the Jaccard overlap.

### 2.4 NetMap uncovers biologically plausible evolving signatures in exhausted T-cells

Lastly, we apply the new NetMap method on a dataset of antigen-specific CD8+ T-cells from P14 mice containing samples at 4 different stages: naïve cells, activated cells 24h after a unique high dose stimulation with gp33 peptide and exhausted T-cells at 72h (D3) and 96h (D4) after daily low dose peptide stimulation [20]. We preprocessed the dataset to retain only viable cells, and then used the standard settings of the NetMap method to obtain the GRN. The four samples form relatively distinct clusters on the expression embedding (Figure 4a), which is expected due to the strong signal induced by the experimental setup. The GRN embedding also captures these signatures, as shown in Figure 4b. We transferred the labels in a cluster-wise manner, using the in-built `unify_group_labelling` function. The clustering matched up to 95% of the cells to the identical condition, with the majority of the mismatches stemming from switching up the two exhausted states (Figure 4c). Due to the progressive nature of the exhaustion process, this is an expected result.

**Fig. 4:**
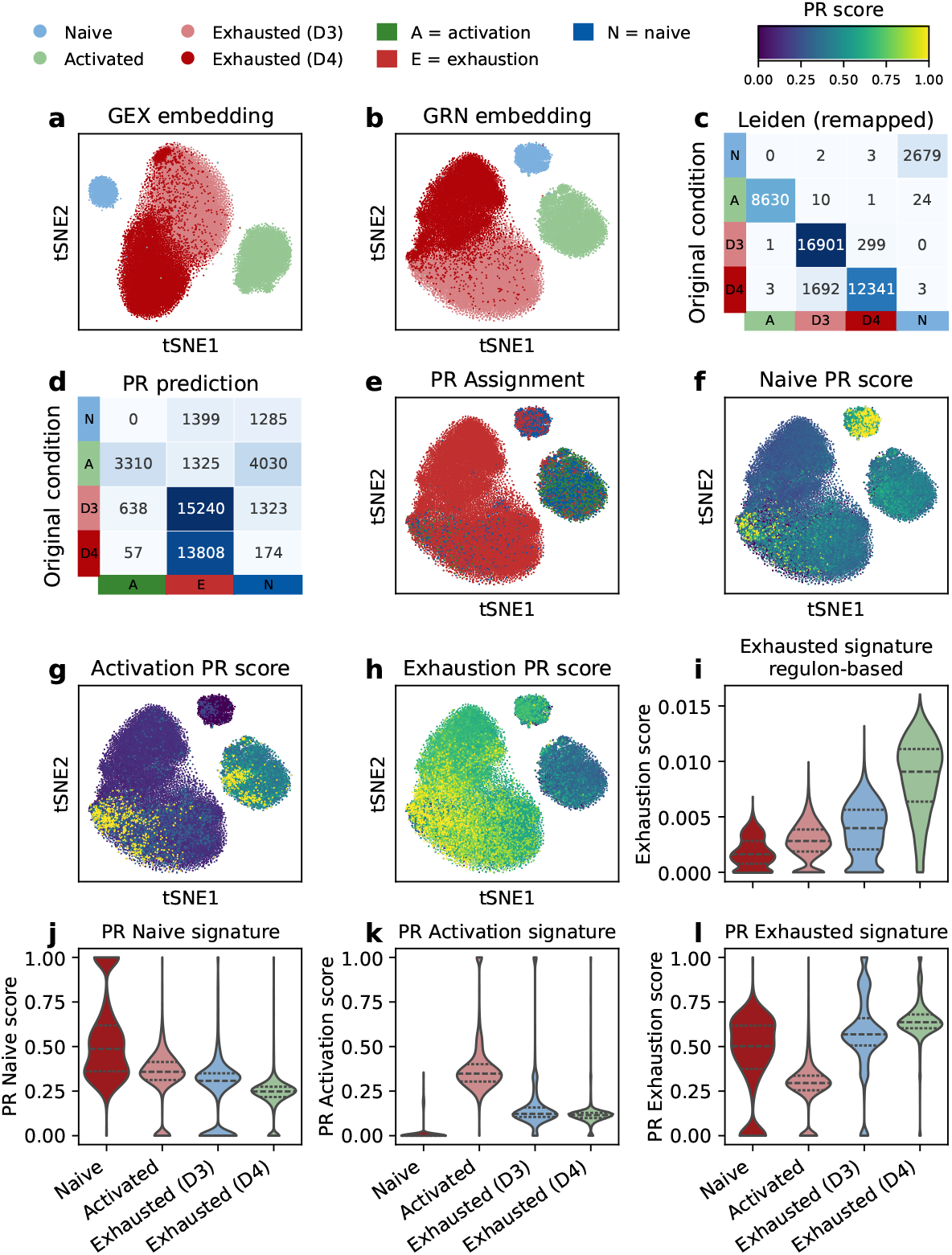
Analysis of the T-cell states during activation and exhaustion. (a) tSNE plots showing the gene expression-based low dimensional embedding and clustering and the (b) GRN-based low dimensional embedding and Leiden clustering with transferred labels. (c) Comparison of the ground truth (dataset) labels with the Leiden clustering of the GRN. (d) Heatmap showing the comparison of the PageRank (PR) based cell assignment with the ground truth labeling. (e) Visualization of the PageRank based cell assignment. (f-h) Qualitative plots showing the PR based score for the three signatures. (i) Regulon based exhaustion score split by batch. (j-l) Violin plots showing the scores (assignment pseudo-probabilities) split by batch.

We also performed the PageRank cell state assignment, using three signatures representing naïve, activated and exhausted states (see Table 6), and compared the assigned labels to the ground truth labeling (i. e., the experimental batch). The majority of the cells were assigned a cell type that matched their ground truth labels (Figure 4d and e). Some were assigned differently, which is only possible because the assignment is done on a per-cell basis, instead of a per-cell type basis.

Next, we analyzed signatures representing the naïve, activated and exhausted states in the T-cells by aggregating edge scores based on a reference gene set (see Section 4.8.2 for details), using the PageRank pseudo-probabilities as well as a regulon-based scoring scheme. As expected, the naïve signature showed a strong signal on naïve cells and low scores on activated or exhausted cells (Figure 4f and j). On the other hand, the activation signature showed the highest score on the activated cells and the lowest scores on naïve cells (Figure 4g and k). We further observe a slight decrease on the activation signal from D3 to D4 of the exhaustion subsets, corresponding to a gradual loss of proper T-cell functioning. At the same time, the exhausted signature is the strongest on the exhausted subsets (Figure 4g, k, h, l and i). When using the regulon-based score, where the GRNs have been explicitly separated (Figure 4i), a more gradual pattern can be observed, which is difficult to identify based on the PageRank assignment, given the limited number of marker genes and the uncertainty regarding the expected changes between D3 and D4. For context, we provide plots in Figure S10 showing similar scores computed on the gene expression data using `sc.tl.score genes`. The pattern is analogous, indicating that both NetMap and the standard analysis align well regarding the identification of high-level biological patterns. The previous analyses show that the networks are not only able to represent genuine biological themes at the signature level, but also that the network structure inferred by NetMap itself encodes these dynamics, making NetMap a suitable tool for the unbiased analysis of complex datasets.

Lastly, we extracted an aggregated network based on the PageRank scoring. The nodes were selected based on their PageRank importance per cell type (top 20 genes) and then merged into one global network where the relative contributions of the cell-types to the gene nodes are indicated via pie-charts (Figure 5). We then analyzed the top 20 most important genes of the global network (Table S11), using the webtool GeneAgent [21], an agentic gene set enrichment tool. We obtained the following summary, which aligns well with the biological question: “In summary, this system of interacting proteins collectively contributes to T cell activation, survival, migration, and effector functions. The interplay between signaling molecules, transcription factors, adhesion proteins, and cytoskeletal regulators ensures a coordinated immune response.”

**Fig. 5:**
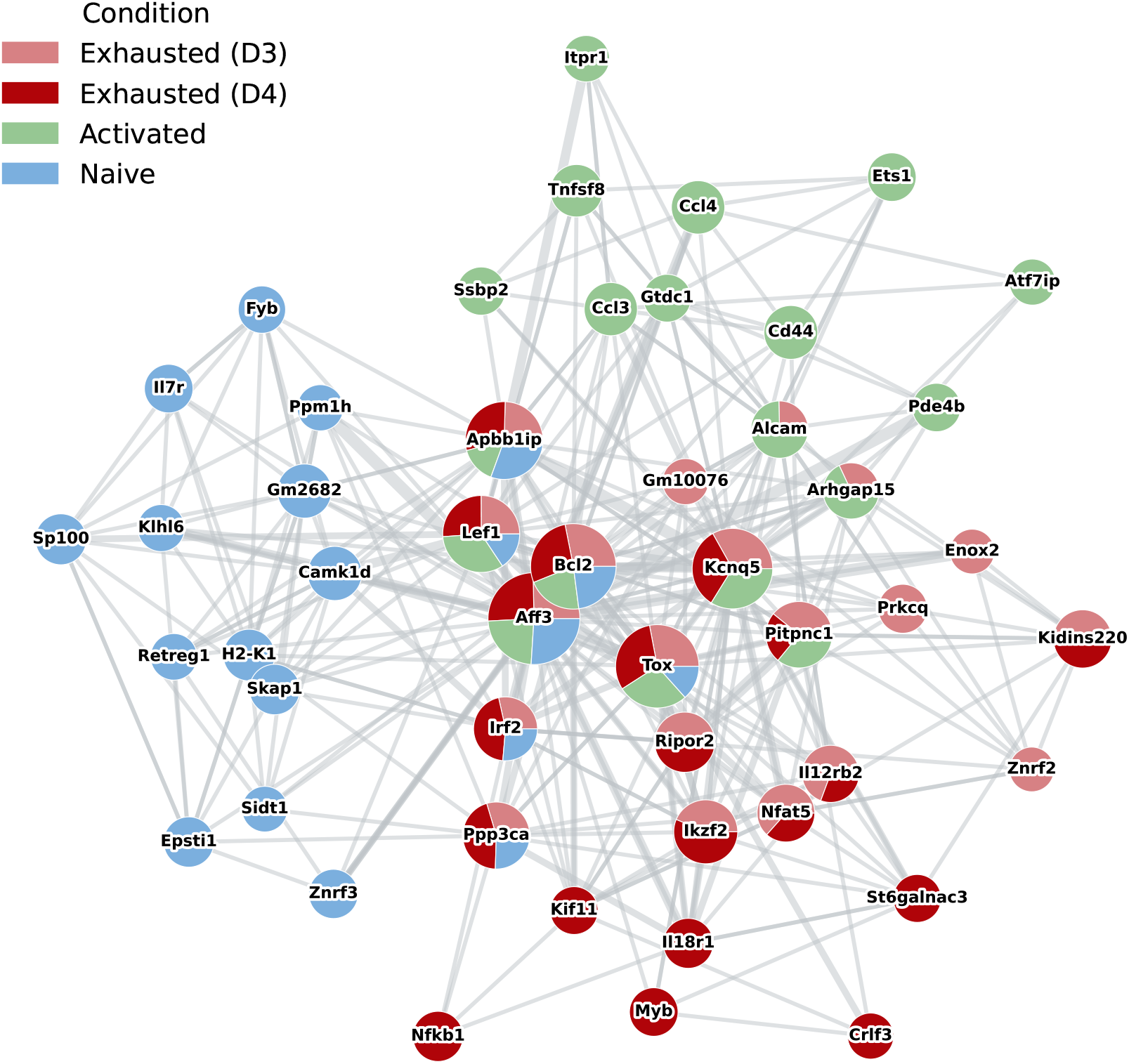
Network generated using the NetMap PageRank version, selecting at most 20 genes per cell and allowing 20 nodes per cell type in the final network. The pie chart shows the relative contribution of the cell type to the gene node with respect to the total importance of the gene in the graph, given the gene appears in the top 20 genes of that cell type. It’s size is proportional to the total importance of the gene. The edge thickness indicates how many conditions an edge was found in.

For comparison, we also produced a ranked list of marker genes across the three states (naïve, activated, exhausted) with the Wilcoxon rank sum test, using only one global state for the exhausted cells to avoid canceling out differential dynamics due to the two relatively similar datasets (see Section 4.8.3 for details). Using the top 20 ranked genes as an input to GeneAgent, we obtained the following summary: “In summary, this system of interacting proteins is primarily involved in chromosome dynamics, immune regulation, and associated processes such as transcriptional control, calcium signaling, and apoptosis. The interplay between these functions highlights the system’s role in maintaining cellular and immune homeostasis.”

We take this as indication that NetMap provides a benefit when analyzing complex datasets in a holistic manner compared to marker analysis, as it was able to detect and up-rank relevant genes. The reports generated by GeneAgent are provided in the Supplementary Material, where we also provide an additional report where the two exhausted states are treated as individual conditions (also here, using NetMap yields more specific results than Wilcoxon rank sum test).

To verify the results with a classical gene set enrichment method, we also ran g:Profiler2 [22] on the same input as used for GeneAgent (results in Tables S6–S10). The terms which are found for the NetMap gene sets are very specific. As an example, when using g:Profiler2 on the top 20 high ranking PageRank genes for the activated condition, we obtain the following terms: *alpha-beta T-cell differentiation* and *T-cell activation*. In contrast, we obtain fewer and less specific terms for the top 20 marker genes obtained from the Wilcoxon rank sum test.

As an example of visualization, Figure 5 shows the combined network indicating shared and specific genes for the four conditions. All subnetworks are connected via transcription factor nodes which have been chosen by their centrality in the network, not by manual curation, indicating their strong driver potential. The naïve and activated states have distinct subnetworks, with the former being marked by Il7r, a gene characteristic of less differentiated and resting cells [23], and the latter identifying genes strongly induced after stimulation (Ccl3, Ccl4, Tnfsf8 and Cd44) as important nodes [24]. In contrast, the exhausted states D3 and D4 share more network nodes, including known exhaustion-associated genes like Nfat5 and Irf2 [25, 26]. The expression of the top 20 genes identified by PageRank shows the clear dynamic changes across the four states (Figure S9), highlighting their relevance in the activation and exhaustion processes.

## 3 Discussion

In this article, we present NetMap, a novel clustering and GRN identification tool. Unlike other workflows, NetMap performs simultaneous clustering and GRN identification using the explainability method GradientShap and can use the GRN directly on cell-level basis to assign cell states.

Using synthetic data, NetMap was compared against scGeneRAI, a conceptually close competitor, and GRNBoost2 which was run with a simplified setup. NetMap outperformed scGeneRAI and GRNBoost2 on the GRN inference sub task. On realistic data, NetMap outperforms the pretrained model scPRINT, but performs worse on the GRN inference task than LINGER. However, the performance gap between NetMap and LINGER is well-accounted for by LINGER’s use of additional data modalities (e.g. a baseline GRN and ATAC-seq data, as well as pretraining) and the requirement for a clustered input.

NetMap achieved notably faster runtimes than scGeneRAI already on small synthetic datasets. This promotes xAI-based combined GRN inference and clustering into the realm of practical feasibility, instead of remaining proof-of-concept work.

NetMap does not assume any prior knowledge and operates solely on the gene expression data, but external data can be used to prune the gene regulatory networks to interpretable modules, e.g. using prior knowledge on transcription factors. Despite using only gene expression data as the input, NetMap achieves favorable performance in comparison to the other methods. This is particularly interesting for datasets of tissues which are underrepresented in the large databases online and may be less well represented by foundation models.

One limitation of xAI methods is that they assume independence of the variables. As gene expression data has high correlation between features, the attribution is split between correlated variables, and therefore the ‘sharpness’ of the signal decreases, making it overall more difficult to distinguish signal from noise. Nonetheless, we observed that in small-scale simulated data, the xAI method is able to distinguish the relevant from the irrelevant edges for each cluster, as we observe higher recall for the clusters in which we should find the edges, compared to the ones where we should not find any edges.

NetMap combines GRN inference with true network analysis in form of the PageRank algorithm, which allows us to overcome a limitation of other GRN inference methods which have to revert to gene set analysis shortly after the network creation step. In this work, we directly employ the network structure in the PageRank algorithm and are hence able to identify important nodes in the datasets. One caveat here is that largely unequal gene set sizes can lead to diluted probabilities in the probabilistic assignment. We correct for this issue by allowing a gene set size correction, however, we recommend to keep the gene sets similarly-sized. Sometimes, the marker gene sets are used in a subjective manner when assigning cell types, such that some marker genes for particular cell types are also present at relatively high expression levels in ‘unrelated’ cell types. In the PageRank, ambiguous markers necessarily dilute the labeling, because the algorithm does not have our implicit knowledge about the coordination of gene expression for cell identity, so the markers have to be chosen with this observation in mind.

In the PBMC dataset with four technical replicates we were able to demonstrate that the clusters and GRNs respect known biological themes, such as the cluster identity, and the overlap with known markers. Unfortunately, the definition of the cell types in the literature is quite divergent. Many genes appear to be used for several, sometimes biologically different cell types (see Figures S6 and S7 for heatmaps of Jaccard overlaps between the cell types most relevant for the blood datasets). This leads to a higher number of outliers for the overlap of gene sets with de-novo derived regulons from NetMap.

We also analyzed a dataset investigating the T-cell activation and exhaustion process and were able to show very clearly the dynamic behavior of the GRNs as the T-cells become increasingly exhausted. NetMap identified driver genes such as Tox without manual selection process. Both gene set enrichment using gprofiler2 and agentic enrichment using GeneAgent showed that the biological themes were very closely matched to the known biology. Compared to a standard cluster-wise Wilcoxon differential analysis, the results yielded by NetMap are more specific. While we were able to pick up on programs such as cytotoxicity and general immune systems using the genes ranked by Wilcoxon rank sum test, NetMap was able to pinpoint more specific processes such T-cell activation and differentiation in most cases. This shows that network based discovery of biological themes is possible using the NetMap method. Additionally, the genes we identified as the top most important genes in the dataset include *Tox*, a known regulator of T-cell exhaustion [27], and *Bcl2*, a gene implicated in T cell survival, activation and exhaustion [28], both having a central role in the network.

To summarize, with NetMap, we provide a tool for fast and convenient GRN inference and analysis, which allows users to move from a gene-centric representation of their data to an edge-centric view without prior annotation of the cell types. NetMap delivers state-of-the-art performance at a reasonable runtime for standard datasets. The tool can be installed as a python package and will be easy-to-use for practitioners who are already familiar with the scverse ecosystem.

## 4 Methods

### 4.1 Description of the NetMap model and training process

The input for NetMap is a gene expression matrix *D*^*n×g*^ with *n* the number of cells and *g* the number of genes. The model is a shallow bottleneck autoencoder which takes *D* as input and reconstructs the raw counts while optimizing the parameters of a negative binomial distribution. After training we use GradientShap to find predictive variables from the input space for a given target gene. The data is normalized to equal count depth and log transformed before training.

More precisely, the shallow bottleneck autoencoder consists of an encoder with a low-dimensional output size, and two decoders which we use to estimate the one to estimate the parameter *µ*, and one for the parameter *θ* of the negative binomial distribution.

Let *L*(*d*_*i*_, *d*_*o*_) be a dense layer with input dimension *d*_*i*_ and output dimension *d*_*o*_. The Encoder is described as *E* (*g, l*) = *L*(*g, d*_*h*_) *° L*(*d*_*h*_, *l*), the decoder is defined as *D*(*l, g*) = *L*(*l, d*_*h*_) *° L*(*d*_*h*_, *g*). Following the implementation of DCA in [6] we optimize the parameters of the negative binomial or zero-inflated negative binomial distribution for each gene and cell. The reconstruction loss of the cell’s gene expression vector *y* is computed using the following formula. *µ* and *θ* are estimated by using the two decoders.

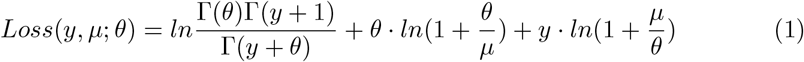

For internal layers, we use `ReLU` as the activation function, for the last layer we use `Sigmoid`. There are optional dropout layers for regularization. The number and dimensions of the hidden layers can be defined by the user, but we found the shallow autoencoders generally outperformed deeper architectures for the GRN inference task. Hyperparameter search results for both the architecture and explainability method are reported in the Supplementary Material (Table S1 and Figures S1-S3), where we show the hyperparameter we evaluated, and the on-target, off-target and clustering coefficients for different setups respectively.

The optimizer is Adam, with a default learning rate of 1*e*^*−*4^. By default, we set a high number of epochs, but implement early-stopping for the training loop by splitting the data into a training and validation set randomly with a ratio of 80:20. We terminate the training prematurely if for at least *e* epochs the validation loss did not decrease by more than 0.001. By default the patience is *e* = 10, with a validation every 10 training steps. These parameters can be adjusted by the user. Due to the high collinearity of the input features and inherent stochasticity, the models can pick up on different features during different runs, which results in random fluctuations of the model. Therefore, in order to improve robustness, we train several versions of the model using different random initialization, and random training and validation split. Since, we do not use a hold-out set for final testing, the models see the entirety of the dataset during the training, but we avoid overfitting the model in the given run.

Following training the model, we use the GradientShap method from the captum library [8] on each model to obtain an attribution matrix for each target. We use only the decoder which was used to estimate *µ*, the mean expression for the attribution, by disabling the other outputs for the forward pass in inference mode. We call *G*^*n×e*^ the final output GRN with *e* = *g*^2^ as we obtain values for every potential pair of genes. Let *s*_*n*,(*i,k*)_ be the feature attribution for gene *g*_*i*_ in the input space to gene *g*_*k*_ in the output space. For each target gene *g*_*k*_, we obtain the full attribution matrix containing values for every input gene *g*_*i*_: *G*_*n*,(*i,k*)_ = *s*_*n*,(*i,k*)_. This way, for every target gene *g*_*k*_, and every model *m* we obtain a complete GRN *G*^*m*^. We then obtain the final explainability matrix over all *m* as 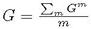.

### 4.2 Postprocessing

After obtaining the final attribution scores per target matrices, we filter the output with masks computed based the raw data to be overlayed on the output. This is straightforwardly achieved as we use the AnnData [29] format which lets us define additional layers.

#### 4.2.1 Missingness mask

The raw input may contain a high number of missing values which can arise either because the cell does not express the gene or because the sequencing technology did not capture the gene. The autoencoder will fill in the blanks by imputing the data. However, this can be a problem when there is a structure in the missingness, for example, a gene is significantly more often expressed in cluster *c*_1_ than in cluster *c*_2_. The autoencoder will fill in the missing values in cluster *c*_2_ based on the values in *c*_1_. This leads to a smooth representation, but may be an over-imputation in some cases. Similarly, the explainability method cannot distinguish these cases and will produce an attribution score for each source and each target. Therefore, we provide two masking strategies for the postprocessing of the raw GRN embeddings, one naïve masking and a knn-masking. In the naïve masking, for every edge *e*_*n*,(*i,k*)_, the edge (*i, k*) in cell *n* the raw GRN is masked where either gene *g*_*i*_ or gene *g*_*k*_ are not expressed in cell *n*. However, as this may be overly strict, we provide a user-configurable knn-version of this procedure, where an edge *e*_*n*,(*i,k*)_ is only masked if it is not expressed in *n* and additionally not expressed in *l* (user defined) out of *r* neighbors in the knn-graph denoted as *KNN*_*r*_(*n*) (computed using `sc.tl.neighbors`). This mask is added as a layer in the GRN output object. The strict mask *M*_*m*_ and the lenient mask *M*_*l*_ are defined below.

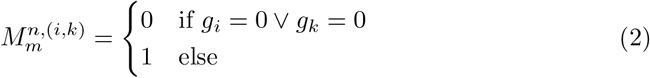

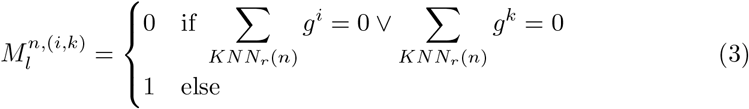

#### 4.2.2 External annotation

We provide external data masks to be used as an additional filtering method. This approach has been made popular by the SCENIC [3] suite. There are several external resources which can be included to streamline the analysis of the NetMap output. We natively provide a TF annotation (*A*_*F*_) and an external GRN mask (*A*_*GRN*_). The annotations are annotated in the variable object of the data. *A*_*TF*_ indicates, whether a source gene is a TF from a provided, or user defined set of regulators *R* or all sources in a GRN (e.g. collecTRI [30] or TTrust [31]). The *A*_*GRN*_ indicates whether an edge has been found in an external or user defined preexisting *G*_*ℛN*_. This mask can be quite strict because our knowledge of transcription factor stored in databases is not vast enough yet.

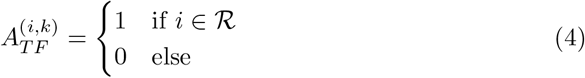

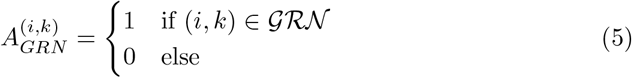

#### 4.2.3 Clustering and edge ranking

The raw GRN object contains cell specific GRNs for all cells, which is an extremely large data object. This information is difficult to interpret directly and needs to be further processed, by clustering and edge annotation. We first cluster the data using the standard workflow of PCA embedding, knn-graph construction and Leiden clustering [32] using the top *p*%, (*p* = 10% by default) of the data. The top scoring variables maintain the structure of the data while allowing most users to use the tool, as the high number of edges will likely exceed the computational resources of many users. After clustering, we use the missingness mask in combination with the cluster labels to determine plausible edges, by computing the proportion of cells an edge is expressed in per cluster, yielding the cluster-wise mask *A*_*p*_. An edge is subsequently selected if it is a relevant edge in at least one cluster resulting in the global edge mask *A*_*EDGE*_:

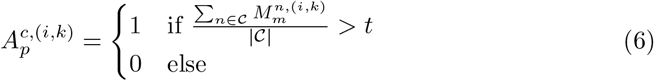

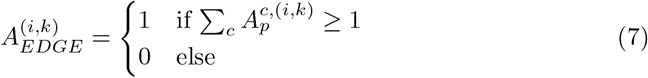

### 4.3 Downstream analysis

Following reconstruction of cell-specific gene–gene interaction networks, we conducted a series of downstream analyses designed to annotate the object, identify regulatory factors responsible for driving cluster-specific transcriptional programs, and facilitate visualization and interpretation of the resulting regulatory architectures.

#### 4.3.1 PageRank based cell type annotation of the GRN object using marker genes

Inspired by the idea described in Cellwalker2 [33], we create a PageRank based annotation method using the GRN graph directly on a cell-by-cell basis. Let *G* = *{g*_1_, …, *g*_*n*_*}* denote the set of genes in the sparsified GRN and *T* = *{t*_1_, …, *t*_*q*_} the set of candidate cell types. For a given cell *c*, let *W*_*c*_ *∈* ℝ^*n×n*^ be the signed, sparsified gene–gene edge weight matrix inferred for that cell. Negative edges are removed to ensure non-negativity, and the remaining edges are re-weighted by their inverse document frequency (IDF) across all cells, so that edges observed in fewer cells receive proportionally larger weight:

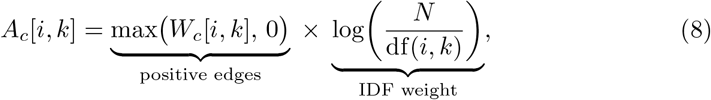

where *N* is the total number of cells and df(*i, k*) is the number of cells in which the edge (*g*_*i*_, *g*_*k*_) is present in the sparsified GRN. A bipartite adjacency matrix *A*_*m*_ ∈ {0, 1}^*n×q*^ encodes uniform, positive-marker relationships between genes and cell types:

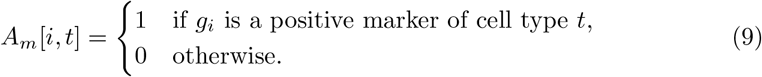

The gene-gene and gene-cell type adjacencies are combined into a single block adjacency matrix over the joint node set *G ∪ T*, where the genes are connected with each other, and the cell types with the genes. The hierarchy or dependency of the cell types is not considered. The adjacency matrix

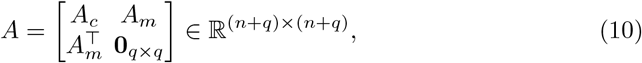

is row-normalized into a transition matrix *P* = *D*^*−*1^*A*, with *D* = diag(*A***1**).

The restart (personalization) vector *v ∈* ℝ^*n*+*q*^ is supported only on gene nodes and set proportional to each gene’s degree in the cell-specific gene–gene graph *A*_*c*_:

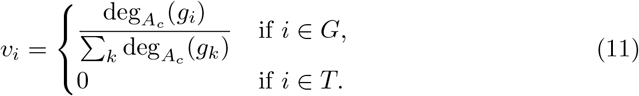

Starting from *π*^(0)^ = *v*, the visiting probability vector is updated for *R* iterations (*R*=200 by default) with restart probability *α* (*α*=0.2 by default):

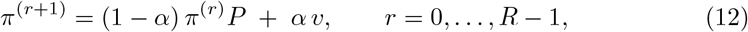

yielding the stationary (or *R*-step approximate) node-visit distribution *π* = *π*^(*R*)^ *∈* ℝ^*n*+*q*^ .

The entries of *π* corresponding to cell type nodes are extracted and renormalized to sum to one, yielding a probability-like score for each cell type:

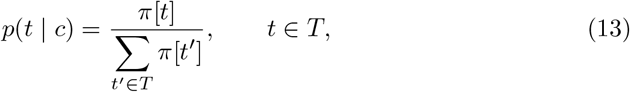

and the discrete cell type label is obtained by taking the arg max over these probabilities:

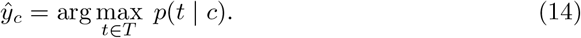

#### 4.3.2 Computation and ranking of signatures

Since the object can contain thousands of edges which on an individual level may have a low signal, we aggregate multiple edges into regulons per cell type. The regulons can be aggregated at the source level and at the target level. To do so, we treat each gene as the focal point and collect the top *k* sources or targets by attribution value accounting for the fraction of cells this edge is expressed in based on the missingness mask we computed earlier. The parameters (*k*, the number of genes selected and *p* the percentage of cells an edge is expressed in) of this step allow the user to be more restrictive or more exploratory. By default the edges are deduplicated, meaning a regulon only contains unique edges with respect to the other regulons. These edges can be combined easily into signatures by computing the mean of the edges. We choose the mean to avoid emphasizing larger regulons. These signatures are computed per cell, hence we can use standard testing from scanpy to compute a rank of the signatures. Since the regulons are computed per cell type and per focus gene, we can obtain multiple regulons with the same focus gene, one for each cell type. To summarize, we compute signatures which comprise several edges into one score, which can be visualized using scanpy functionalities.

#### 4.3.3 Integrated visualization of regulatory programs

The NetMap object is fully compatible with the scanpy ecosystem, meaning many plots can be used without modification. This includes the t-SNE plots as well as various other plots. By transferring the embedding coordinates, of either object, the gene and edge values can be plotted in the same space. We provide a modified version of the gene expression dotplot to account for the regulon composition. In the scanpy dotplot, the dotsize is proportional to the number of cells where the gene is expressed. In the NetMap version, we override the sizing functionality of the dotplot to show the regulon dots proportional to the number of participating edges. To do so, for a signature *U*_*s*_ consisting of all edges *S* = {*e*_*s,t*_ }in the set *S* of top edges starting from a source *s*, we sum up the clusterwise edge counts, normalize them by the number of cells in the cluster and finally normalize the value by dividing by the max across all clusters and regulons. This enables a faithful representation of the regulon activity in the dotplot. Lastly, we add a scatter plot option which allows to visualize the regulon scores and annotating the axis with the regulon GRN they contain.

For the NetMap visualization, we provide a custom network plot, which depict the nodes as little pie charts, showing the relative contribution of a cell type to the node centrality. Node size encodes the total importance of a gene *I*_*g*_ = Σ_*c*_ *f*_*g,c*_ as the sum across cell types of *f*_*g,c*_ over the fraction of a given cell type’s cells in which the gene appears among the top-*n* PageRank genes for that cell. The wedges partition this importance into *f*_*g,c*_*/I*_*g*_, the relative contribution of each cell type to the gene’s total recurrence, colored by cell type; edge thickness is proportional to the number of cell types in which the corresponding gene–gene interaction was independently found among the top mask-supported GRN edges.

### 4.4 Technical details

NetMap is implemented as a python package using pytorch [34], captum and the scverse ecosystem, including scanpy and anndata. The gene expression matrix is passed as a torch tensor and the model is trained on the GPU. We implement the inference process in a target wise manner on the GPU, and move the matrix of per-target attribution values to disk, as the matrix becomes extremely large. To do so, we use the `h5py` library reducing memory overhead during the inference, without substantial runtime cost. The output of the method is stored in an AnnData object, a standard format for single-cell analysis that facilitates downstream processing. The returned AnnData object only holds the observation data frames and the variable annotation, including the sum and mean of each variable, and a reference to the file where the GRN is stored. This allows the user to directly load the top edges using the annotation in the variable without having to load the entire object which is likely infeasible due to memory restrictions, when proceeding to the downstream analysis. Furthermore, the user can use an index base data loading method. Suitable functions are implemented in the NetMap library. Users are encouraged to discard the full GRN object and only keep the top edges after the analysis to restore disk space. Most plotting functions can be directly used from the scanpy library. We provide additional tooling for the plotting of the networks.

### 4.5 Evaluation on synthetic data

To demonstrate that NetMap is capable of recovering cell type-specific GRNs, we first evaluated its capacities using synthetic data. Using scMultisim [16] which takes as an input a GRN, we generate synthetic gene expression (GEX) data. Then we apply the NetMap workflow on this synthetic data with the goal to reconstruct the GRN that was originally used for the data generation. In brief, we first argue that scMultisim is suitable for generating differential networks. Then we describe how to obtain small subnetworks corresponding to a joint regulatory program of several cell types, and finally how to modify these GRNs to get differential behavior for the cell types we want to model.

#### 4.5.1 Synthetic data generation

scMultiSim [16] is a recently developed data generation tool for multimodal omics data based on a gene regulatory network (GRN). The tree controls the number of cell populations generated and whether to simulate differentiation trajectories. The GRN models the dependencies of the genes and is provided as a table with a source and a target gene and an edge weight. As we are interested in modeling a realistic dataset, we used a publicly available transcription factor network, collecTRI as a starting point. CollecTRI [30] contains a large number of transcription factor target interactions. We are only interested in the empirical structure of those networks. This approach avoids handcrafting networks which may not be realistic, however it of course introduces a certain research bias. We use a greedy modularity optimization method to partition the network into modules of a desired size.

Natively, scMultisim uses one underlying GRN which controls all the cell populations. The algorithm first generates cell (CIV) and gene identity vectors (GIV) which can be interpreted as a latent space. Then the effect of the GRN is added on top of these CIV and GIVs. Consequently, when fixing the random seed in several consecutive runs this latent space will be (almost) identical. In order to generate two cell populations which differ only due to a GRN, running the algorithm with the same random seed and differential GRNs which induce the desired effect into the generated gene expression data. After fixing the randomness of scMultiSim such that the networks where the only factors changing in repeated runs of scMultisim, we generated one dataset per GRN and subsequently combine them into the final multi-cluster datasets. We simulated datasets with 2 and 3 clusters respectively. Additionally, we included a background GRN which is switched on in all clusters to simulate a common ‘housekeeping’ gene regulatory program. To each cluster *c*_*i*_ we want to introduce into the data, we randomly assign a module *GRN*_*i*_ with non-zero edge weights. In each other cluster *c*_*k*_, we set the edge weights of *GRN*_*i*_ to a pseudocount (0.01). This way, we ensure that scMultiSim is run with the same genes, but differing edge weights for each network. We choose scMultiSim parameters which maximise the effect of the GRN in the data, compared to other effects which can be simulated by scMultisim. Finally, we also introduce a joint regulatory program, which consists of a GRN with identical edge weights for all clusters, in order to generate a more realistic dataset. To summarize, we produce data where each cluster has its own GRN while keeping the rest of the data generation process fixed. scMultisim can generate data which includes realistic noise, therefore we generate two versions of each dataset one with, and one without noise. In total, we simulate datasets with four configurations, 2 and 3 clusters with and without noise respectively.

#### 4.5.2 Compared methods

We compare NetMap to its closest competitor scGeneRAI [9] as well as GRNBoost2 with a relaxed setting as a naïve baseline. For NetMap, the best configuration identified during the hyperparameter optimization (Negative Binomial, or zero-inflated negative binomial with one hidden layer with 64 neurons and a latent size of 8, and using GradientShap) is used as the compared method. The hyperparameter search strategy is described in the Supplementary Material. scGeneRAI is a method which is closely related to the NetMap approach. The method is an end-to-end framework using a masking based strategy, in which arbitrary genes are masked, and the input is reconstructed. They use an inflated autoencoder architecture, where the latent dimension is larger than the input/output dimensions. For each target (output neuron) the most important input nodes are identified by using layer-wise relevance propagation. The final output of the model is an average over several runs. As a naïve baseline, we choose GRNBoost2 [4] as a fast competitor, which is still among the top-performing methods in recent benchmark [35, 36]. GRNBoost2 is designed to infer the GRN of a single homogeneous group of samples. Therefore, we simplify the validation procedure running the tool on each cluster individually, using the ground truth cluster labels. This returns one GRN per cluster, however the cluster is predefined, making the task overall easier.

#### 4.5.3 Evaluation metrics

##### Clustering comparison

The NetMap pipeline directly clusters the GRN embedding. The clustering should be broadly comparable to an embedding created based on the gene expression data, using standard preprocessing pipelines, especially for datasets with a strong signal. Therefore, we can compare the gene expression based clustering to the NetMap clustering and compute a score for how well the novel clustering corresponds to the expected clustering. We use the maximum linear sum assignment (MLSA) score to match the novel clusters to the synthetic ground truth labeling and compute the score as the MLSA-score divided by the number of cells to be comparable across data set sizes. A high score between the NetMap clustering and the GEX-clustering indicates that both representations pick up on the same signal.

##### GRN comparison metrics

Additionally, to assess how accurately the tool reconstructs the network which we used to generate the gene expression data, we compute standard confusion tables (true positive (TP): edge was present in ground truth and in inferred network, false positive (FP): edge was found in inferred network but was not present in ground truth, false negative (FN): edge was present in synthetic network but was not found by NetMap), where we disregard TN (edge neither in ground truth nor inferred GRN) as they represent the overwhelming majority of edges.

Recall that we generated data where each cluster is defined by its characteristic GRN meaning that we need to evaluate each dataset with multiple GRNs. We define several metrics to assess how well our method reconstructs GRNs and how specific these are to the dataset at hand. For each cluster we compute the on-target recall *O*_*on−target*_, indicating how well we can recover the edges that we should find. The higher this value, the better. We also compute the ‘off-target’ recall *O*_*off−target*_ for each GRN, computing the metrics for the clusters where the GRN is switched off in the data simulation. This measures the degree to which edges that should not be recovered are nonetheless identified, hence the lower the number, the better. We use these two metrics because the precision tends to be quite low for real-sized datasets in most GRN benchmarks. Let *G*_*i*_ = (*V, E*) with *E* = (*g*_*i*_, *g*_*k*_, *w*_*c*_) be the predicted GRN for cluster *c*_*i*_ with *w*_*c*_ the number of cells the edge was found in, and *GRN*_*i*_ the groundtruth network induced in cluster *c*_*i*_ in the simulation process. We factor in the number of cells an edge has been found in, and clip the recall such that no more edges than |*c*_*i*_| are allowed to contribute positively to the recall for cluster *c*_*i*_. The weighted recall is defined as follows:

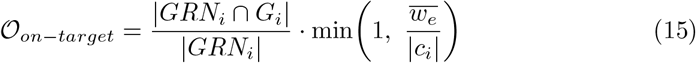

and *O*_*off−target*_ analogously.

In the parameter sweep, the area over/under the diagonal of the recall curve is computed to compare the different combinations with each other. The diagonal indicates random performance, i.e. the number of edges we would expect to be recovered by chance. Using the diagonal as a reference, we can indicate whether we find more edges than expected by chance, but also whether we find fewer edges than expected by chance, which is useful to evaluate the cluster specificity.

#### 4.5.4 Summary of the evaluation procedure on synthetic data

To summarize, we evaluate the method using synthetic data generated using scMultiSim, with groundtruth networks generated by clustering the collecTRI transcription factor network. We use the NetMap and scGeneRAI methods in combination with Leiden clustering to obtain the clusters, but we use the cluster groundtruth GRNBoost2. We compute on-target and off-target scores to ensure that our method indeed retrieves information pertinent to the clusters. Table 2 summarizes all synthetic datasets used in this evalution.

**Table 2:**
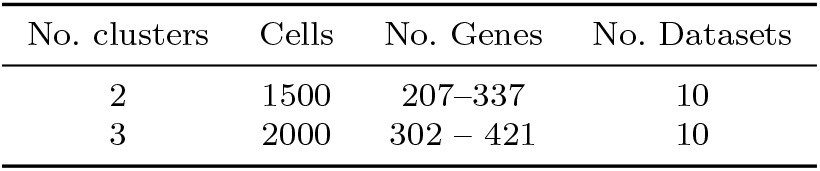
Summary of the generated datasets for evaluation using synthetic ground truths.

**Table 3:** Dataset composition and evaluated transcription factor per benchmarking case. Each case consists of two cell populations drawn from the PBMC multiome dataset, with ChIP-seq binding targets of the evaluated TF serving as the ground-truth reference.

| Case | Cell Type 1 | Cell Type 2 | Cells (n) | Genes (n) | Evaluated TF |
| --- | --- | --- | --- | --- | --- |
| Case 1 | Classical monocytes | Memory CD4 T-cells | 514 | 1,500 | IRF4 |
| Case 2 | Intermediate monocytes | Naïve CD4 T-cells | 3,388 | 1,500 | SPI1 |
| Case 3 | Intermediate monocytes | Naïve CD4 T-cells | 1,999 | 2,000 | FOXP3 |

### 4.6 Evaluation against competing methods on real data

To benchmark NetMap against state-of-the-art GRN inference methods LINGER and scPRINT, which we were unable to benchmark on synthetic data, due to data requirements or pretraining, we use real data, again including GRNBoost2 as a baseline. The three methods each require cell type cluster labels as input and are applied independently to pre-filtered, cell type-specific subpopulations. LINGER additionally utilized the paired scATAC-seq modality alongside predefined regulatory element annotations. NetMap, by contrast, receives the full dataset without any cell type annotations, clustering, or prior population structure. All methods were evaluated on a publicly available PBMC dataset from 10*×* Genomics providing paired scRNA-seq and scATAC-seq measurements from the same cells [17]. Predicted transcription factor– target interactions were compared against ChIP-seq binding evidence from blood cells sourced from the Cistrome database [17, 37] as an independent ground-truth reference.

#### 4.6.1 Datasets

Three evaluation cases were constructed from the PBMC multiome dataset, each pairing a monocyte subtype with a T-cell subtype to represent biologically distinct regulatory programs. Cases 2 and 3 share the same cell types but differ in the number of highly variable genes used as input features, allowing the effect of feature set size to be assessed independently of cell type composition. For each case, evaluation was performed with respect to a single transcription factor with available ChIP-seq binding evidence in the relevant cell type. All cases are summarized in Table 3.

Highly variable genes were selected as input features in all cases. Only scRNA-seq data (raw counts) were provided as input to NetMap, GRNBoost2, and scPRINT. LINGER additionally utilizes the paired scATAC-seq modality alongside predefined regulatory element annotations. For each case, the ground-truth reference for benchmarking was obtained from identifying the targets of one transcription factor in the corresponding cell type using ChIP-seq experiments.

#### 4.6.2 Compared methods

Four methods were evaluated: NetMap, LINGER, GRNBoost2, and scPRINT. As noted above, LINGER, GRNBoost2, and scPRINT were each provided with groundtruth cluster labels and applied separately to pre-filtered, cell type-specific subpopulations, while NetMap was applied to the full mixed dataset with no cell type information. Direct performance comparisons should be interpreted in light of this asymmetry. NetMap was applied to the full unclustered dataset using the best configuration identified during hyperparameter optimization: a Negative Binomial or zero-inflated Negative Binomial model with one hidden layer of 64 neurons, a latent size of 8, and GradientShap as the explainability method (Supplementary Material). No cell type labels, chromatin accessibility data, or regulatory priors were provided.

LINGER was run using its standard `LINGER` mode, which employs lifelong machine learning based on neural network models pre-trained on atlas-scale external bulk data from ENCODE and subsequently refined on the single-cell multiome data. Input data were prepared as paired scRNA-seq and scATAC-seq AnnData objects, with cells filtered by ground-truth cluster labels prior to inference, providing the method with cell type-homogeneous input. Pseudo-bulk profiles were generated from the single-cell data per sample and overlapped with the general GRN using the human reference genome hg38. The resulting cell type-specific *trans*-regulatory score matrix was used for evaluation against ChIP-seq ground truth.

GRNBoost2 was run using the `arboreto` implementation (`arboreto.algo.GRNBoost2`). Raw count data were loaded as an AnnData object, library-size normalized using `sc.pp.normalize total`, and log-transformed with `sc.pp.log1p`. The normalized expression matrix was converted to a dense format and provided as input. In contrast to the other methods, GRNBoost2 was applied exclusively to the target cell type subpopulation, filtered by ground-truth cluster labels prior to inference. Unlike in a standard GRNBoost2 call, where all the transcription factors are used as potential regulators, only the evaluated transcription factor for each case (IRF4, SPI1, and FOXP3 for Cases 1, 2, and 3, respectively) was supplied as the candidate regulator, restricting inference to the edges originating from that single TF. The resulting ranked edge list was pivoted into a TF *×* target gene importance matrix, which was used for evaluation against the ChIP-seq ground truth. scPRINT was run using its pre-trained large foundation model, loaded from checkpoint with a standard transformer configuration and no precomputed gene embeddings. Raw count data were preprocessed using Ensembl identifiers, requiring a minimum of 500 valid gene identifiers, and with cell and gene count filtering disabled (`filter_gene_by_counts=False`, `filter_cell_by_counts=False`). Gene symbols were mapped to Ensembl identifiers prior to preprocessing; genes without a valid mapping were excluded. GRN inference was performed using the `GNInfer` task, configured to select the most variable genes across cell types (`how=‘‘most var across’’`), with softmax preprocessing of the attention matrix (`preprocess=‘‘softmax’’`), no head aggregation (`head_agg=‘‘none’’`), and expression passed directly through the model without a denoising forward pass (`forward_mode=‘‘none’’`). The top 1,500 genes were used (`num_genes=1500`), with a maximum of 1,024 cells per cell type (`max_cells=1024`), a batch size of 8, and all transformer layers included. Inference was performed separately for each target cell type using ground-truth cluster labels. Attention matrices across all heads were aggregated by averaging (`grn.varp[‘GRN’].mean(-1)`), and the resulting gene*×* gene attention weight matrix was used for evaluation against ChIP-seq ground truth.

#### 4.6.3 Performance metrics

We used the built-in clustering transfer method to match the pre-annotated clusters to the GRN clusters and computed the clustering score. Performance for all methods was assessed using the area under the receiver operating characteristic curve (AUC), computed against ChIP-seq binding evidence as ground truth. As ground truth we used the putative target genes of each transcription factor derived from ChIP-seq data in the Cistrome database[37], applying the same benchmarking procedure as LINGER. For each transcription factor, the Cistrome gene scores were aggregated to the maximum score per gene symbol, the genes were ranked by this score, and the 500 top-ranked genes were labelled as true targets. All other genes in a method’s output were labelled as non-targets, including genes with lower-ranked binding evidence and genes absent from the Cistrome dataset. Each method was then evaluated on the regulatory scores it assigned to the target genes of that transcription factor, with higher scores taken to indicate stronger regulation, and the AUC was computed from the resulting receiver operating characteristic curve.

### 4.7 Application on PBMC datasets

#### 4.7.1 Data preprocessing

To demonstrate how NetMap can uncover plausible regulons on realistic data sets, we ran NetMap on four datasets of PBMCs stemming from a sequencing platform comparison [38], where the same sample was sequenced using different technologies. We processed the raw data using nf-core/scrnaseq setting kallisto as the aligner for the 10X Genomics datasets and with the BD Rhapsody provided pipeline for the BD Rhapsody datasets. Then we used scanpy and scrublet [39] to filter the data and remove doublets. The annotation was performed using celltypist [40] and corrected using marker genes. Specifically, we generated a manual consensus from the high- and low-resolution celltypist models, to obtain a coarse cell type mapping and used marker genes, to obtain a finer annotation for the T-cells and the Monocytes. After annotating the data, we removed genes which were not among the highly variable top 2500 genes, and further removed genes which were expressed in less than 5% of the cells. We lastly removed genes which have a high ubiquitousness index (*>* 0.1 for human) according to PanglaoDB [18] which removes ribosomal and mitochondrial genes. The ubiquitousness index was manually parsed from the website. Table 4 shows the sizes of the datasets after filtering.

**Table 4:**
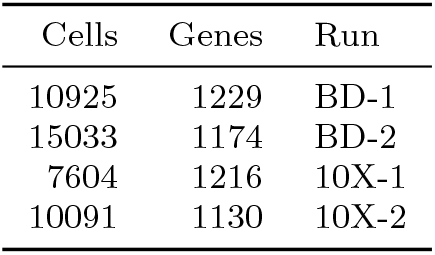
Blood dataset summary.

| Cells | Genes | Run |
| --- | --- | --- |
| 10925 | 1229 | BD-1 |
| 15033 | 1174 | BD-2 |
| 7604 | 1216 | 10X-1 |
| 10091 | 1130 | 10X-2 |

#### 4.7.2 NetMap embedding

We used NetMap with default training parameters. Afterwards, we selected the top 10% edges, based on the aggregated sum of the feature attributions, and applied PCA and Leiden clustering to the dataset like in the standard workflow. We compared the clustering we obtained from the standard gene expression data embedding pipeline (PCA, tSNE, kNN, Leiden clustering) with the clustering we obtain from the NetMap, using NetMap’s built-in functionality and manually adjusted the resolution to be able to meaningfully map the clusters from NetMap to the gene expression clustering. We then compute the missingness mask for all edges present in the object, and determine if they are candidate edges in any cluster (edge possible in more than 50% of the edges). We then load the candidate edges for further processing and discard the previously loaded edges. We then apply the edge selection per regulon, by selecting the top 30 targets per source for each regulon and further removing the edges which are selected for multiple clusters at the same time (the deduplication is optional, as it is possible that edges are shared between regulons). The regulons are aggregated by taking the mean edge value and a new signature object is created. This signature was processed with the standard scanpy processing. Additionally, we label the cell types using the PageRank label assignment to obtain the probabilistic assignment using default parameters with the cell type markers for the PBMC data shown in Table 5.

**Table 5:** PBMC cell type marker genes.

| Cell type | Marker genes |
| --- | --- |
| B cell | CD19, CD79A, CD21, MS4A1, PAX5 |
| CD4+ T cell | CD4 |
| CD8+ T cell | CD8A, CD8B, GZMA, GZMK |
| NK cell | NKG7, NCAM1, FCGR3A, GZMB |
| CD14+ Mono | VCAN, S100A8, S100A9 |
| CD14- monocyte | FCGR3A, CDKN1C |
| DC | CD1C, CST3, FLT3 |
| pDC | PTCRA, SMIM5, LAMP5, JCHAIN |

**Table 6:**
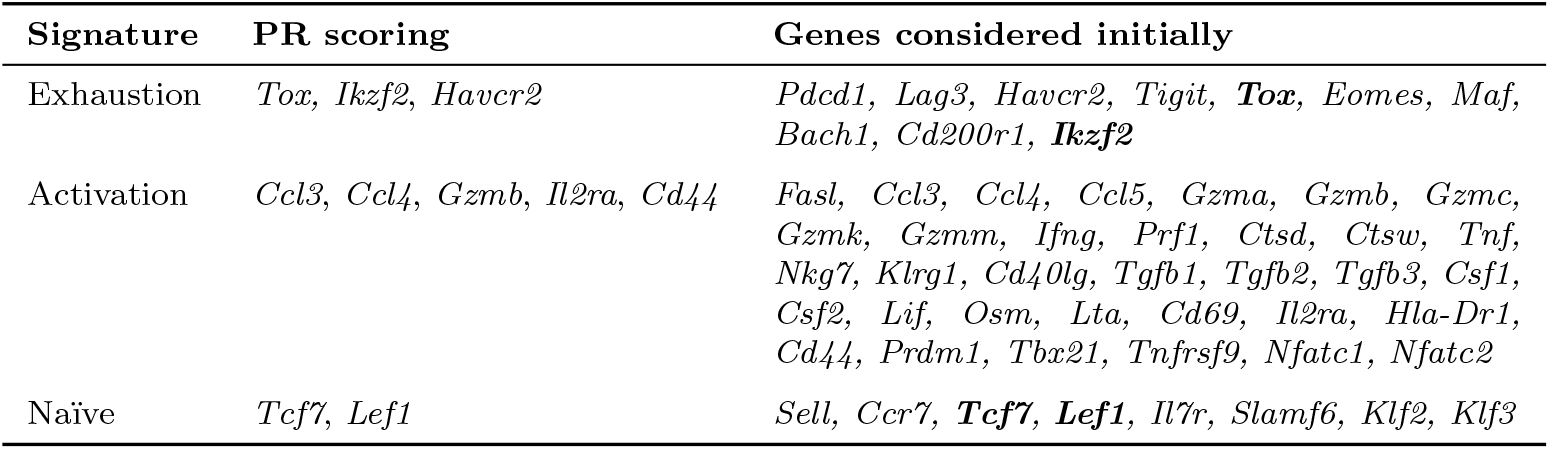
Gene signatures used for scoring. The genes in the column PR scoring are used for the PageRank. For completeness the genes considered initially are shown in the third column.

#### 4.7.3 Regulon validation strategies

We then proceed to compute metrics to show that the NetMap signature object contains biologically plausible information. We first download the marker gene databases PanglaoDB [18] and CellMarker [19] and compute the Jaccard overlaps between the marker sets and the top ranked regulon sources and their top targets separately. PanglaoDB is a curated database containing one marker set per cell type and Cell-Marker is a collection of marker sets from the literature without a consensus labeling. We subset the datasets by selecting the tissues of interest (Blood/Immune cell types). For PanglaoDB we regrouped the subtypes into broader categories, e.g. Naïve B-cells and Memory B-cells into the supercategory B-cells. The CellMarker database does not provide consensus annotations, so we clustered the marker gene sets based on their Jaccard overlap to obtain groups of semantically similar marker sets despite their inconsistent labeling. We first computed the pairwise Jaccard index between all sets of marker genes. Then, we used hierarchical clustering to obtain clusters of marker genes. We then fit a quantile regression between the tree cophenetic distance and the Jaccard index and selected the tree distance threshold such that the average Jaccard distance was greater than 0.2. We extracted the labels per cluster and manually selected a consensus description for the gene sets. This way, we retained the raw information of the original annotations, while being able to group the sets according to the cell type categories we had in our data. The label correspondences can be found in Table S2. We obtained the ubiquitousness index files by extracting the table from the website using a scraping script. We include the script in our validation github repository and the file in the NetMap resources.

### 4.8 Application on exhausted T-Cell dataset

#### 4.8.1 NetMap embedding

We obtained scRNA-seq data from a mouse time course experiment with 4 conditions: Naïve T-cells, activated T-cells, T-cells after 72h and 96h from GEO (GSE268165) [20]. Due to the lower sequencing depth of the naïve subset, we performed stringent QC and selected the subset of barcodes which are within 3 median absolute deviation (MAD) for the counts, the number of RNA features and the percentage mitochondria using scuttle. Additionally, a hard threshold was set to include only barcodes with at least 500 genes, and the mitochondrial content falling between 5 and 25%. As autoencoders impute the data, we additionally set a threshold and only included genes which were expressed in at least 3% of the cells. We further removed genes with a high ubiquitousness index of 0.3 (for mouse). Due to this strict filtering, we manually included genes which we know are relevant for the dataset found in Table 6. We note however, that these are extremely lowly expressed (see Figure S8). We then trained the NetMap with 10 models and standard parameters. As per our downstream process, we then used the top 10% of the edges by global weight to cluster the object, verify that we obtain the expected clustering, which in this case was expected to conform to the time point, as the effect of the experiment is strong. As this was the case we then used candidate edges in which the edge was expressed in at least 1% of the cluster’s cells to retain as much information as possible for the downstream analysis. This led to the exclusion of edges of the genes which were expressed at low level.

#### 4.8.2 Network based signatures and PageRank centralities

We perform the downstream analysis by using dimensionality reduction and tSNE to visualize the GRN object in a low dimensional space and cluster the data using the Leiden algorithm. We then use the label transfer function (`unify_group_labelling`) to match the GRN based Leiden clusters with the experimental batches and compute the contingency matrix. Based on the Leiden clusters, we retrieve the gene centric regulons, which we then used to compute the signature scores using all incident edges to the signature marker genes defined in Table 6. We chose to use all edges instead of the deduplicated ones in this instance, because we expect the two exhausted states to be quite similar in gene expression pattern. Using a deduplication procedure would artificially remove edges which have a stronger effect in both D3 and D4 compared to the Naïve and Activated states, and hence would be erroneously removed. We then proceeded to compute network based signatures using the signatures defined in table 6 and the standard settings of the PageRank function with 200 iterations, IDF correction and the normalization by the number of markers used in the PageRank call. We then compare the labeling obtained by scoring each cell with the ground truth labels which correspond to the experimental batch using a contingency table.

#### 4.8.3 Enrichment analysis

Using the PR probabilities, we obtain the ranked list of the top 20 genes which are descriptive for the entire dataset. This information is automatically returned by the graph creation method, which provides a global gene weight ‘importance’, as well as cluster probabilities, indicating the participation of nodes of a phenotype in a particular node. Using this list, we performed the GeneAgent search. In order to compare these results to the standard Wilcoxon rank sum test based gene lists, we aggregated the ranks of all conditions into one global gene rank: For each condition, we performed differential expression testing using the Wilcoxon rank-sum test and ranked genes from most upregulated to most downregulated. For each gene and condition, we computed a distance metric *d* = min(*r −*1, *n −r*), where *r* is the gene’s rank and *n* is the total number of genes, to obtain a proxy degree to which a gene is differentially regulated. Per-condition distances were then summed to yield a single score per gene, and genes were re-ranked by this aggregated distance. We then took the top 20 genes with the lowest rank and performed a GeneAgent search. We performed two searches for the Wilcoxon setup, one where we use the four annotated conditions (Naïve, Activated, Exhausted (D3) and Exhausted (D4)) as groups, and one where we simplified the grouping to Naïve, Activated, and Exhausted. The rationale behind this is, that the two exhausted conditions are more similar to each other than the other ones. Since we are interested in the global dynamics of the dataset from Naïve to Exhausted. We also applied gprofiler2 on Wilcoxon ranked list per condition individually, as well as on the per-condition high importance genes from the PageRank call, which are slightly different sets that the global high ranking genes.

## Supporting information

Netmap Supplement

## Acknowledgements

All authors were funded by the German Federal Ministry of Research, Technology and Space (BMFTR, 031L0309A, 031L0309B, 031L0309C), DBB and AH were funded the German Research Foundation (DFG, CRC1755 CAS-CAID, project Z01), and the German Cancer Aid (70116894). DBB was funded by the German Research Foundation (DFG, 516188180). MO and JB were funded by the German Federal Ministry of Research, Technology and Space (BMFTR, 031L0309B, NetMap). JB was funded by the European Union under grant REPO4EU (nr. 101057619). Views and opinions expressed are however those of the author(s) only and do not necessarily reflect those of the European Union or the European Research Executive Agency. Neither the European Union nor the granting authority can be held responsible for them. This work was also partly supported by the Swiss State Secretariat for Education, Research, and Innovation (SERI) under contract No. 22.00115. MO and JB were funded by the Deutsche Forschungsgemeinschaft (DFG, CRC 1700, project SP01).

## Data and code availability

- The NetMap method is available as a python package on github https://github.com/bionetslab/netmap and can be installed using pip. We provide pixi environments for the easy deployment of the method.
- The code used for the benchmarking is available in a separate github repository https://github.com/bionetslab/netmap-evaluation.
- We package the code for the data simulation as an R package which can be installed from github (https://github.com/AnneHartebrodt/grn2gex). The main functionalities include the clustering of a user defined network and the production of the data as specified above using a subset of the functionalities scMultiSim provides.
- CollecTRI has been downloaded via the decoupler [41] package.
- The list of human transcription factors has been obtained from https://resources.aertslab.org/cistarget/tf_lists/.

## References

[1] Pratapa, A., Jalihal, A. P., Law, J. N., Bharadwaj, A. & Murali, T. M. Benchmarking algorithms for gene regulatory network inference from single-cell transcriptomic data. Nature Methods 17, 147–154 (2020). URL 10.1038/s41592-019-0690-6.

[2] Zhang, S. et al. Inference of cell type-specific gene regulatory networks on cell lineages from single cell omic datasets. Nature Communications 14, 3064 (2023).

[3] Bravo González-Blas, C. et al. Scenic+: single-cell multiomic inference of enhancers and gene regulatory networks. Nature Methods 20, 1355–1367 (2023). URL 10.1038/s41592-023-01938-4.

[4] Moerman, T. et al. Grnboost2 and arboreto: efficient and scalable inference of gene regulatory networks. Bioinformatics 35, 2159–2161 (2018). URL 10.1093/bioinformatics/bty916.

[5] Tang, J., Wang, C., Xiao, F. & Xi, R. Single-cell gene regulatory network analysis for mixed cell populations. Quantitative Biology 12, 375–388 (2024).

[6] Eraslan, G., Simon, L. M., Mircea, M., Mueller, N. S. & Theis, F. J. Single-cell rna-seq denoising using a deep count autoencoder. Nature Communications 10 (2019). URL 10.1038/s41467-018-07931-2.

[7] Lopez, R., Regier, J., Cole, M. B., Jordan, M. I. & Yosef, N. Deep generative modeling for single-cell transcriptomics. Nature Methods 15, 1053–1058 (2018). URL 10.1038/s41592-018-0229-2.

[8] Kokhlikyan, N. et al. Captum: A unified and generic model interpretability library for pytorch (2020). URL https://arxiv.org/abs/2009.07896.

[9] Keyl, P. et al. Single-cell gene regulatory network prediction by explainable ai. Nucleic Acids Research 51, e20–e20 (2023). URL 10.1093/nar/gkac1212.

[10] Yuan, Q. & Duren, Z. Inferring gene regulatory networks from single-cell multiome data using atlas-scale external data. Nature Biotechnology 43, 247–257 (2024). URL 10.1038/s41587-024-02182-7.

[11] Kalfon, J., Samaran, J., Peyré, G. & Cantini, L. scprint: pre-training on 50 million cells allows robust gene network predictions. Nature Communications 16, 3607 (2025).

[12] Cui, H. et al. scgpt: toward building a foundation model for single-cell multiomics using generative ai. Nature Methods 21, 1470–1480 (2024). URL 10.1038/s41592-024-02201-0.

[13] Chen, H. et al. Quantized multi-task learning for context-specific representations of gene network dynamics (2024). URL 10.1101/2024.08.16.608180.

[14] Virshup, I. et al. The scverse project provides a computational ecosystem for single-cell omics data analysis. Nature Biotechnology 41, 604–606 (2023). URL 10.1038/s41587-023-01733-8.

[15] Eraslan, G., Simon, L. M., Mircea, M., Mueller, N. S. & Theis, F. J. Single-cell rna-seq denoising using a deep count autoencoder. Nature communications 10, 390 (2019).

[16] Li, H., Zhang, Z., Squires, M., Chen, X. & Zhang, X. scmultisim: simulation of single-cell multi-omics and spatial data guided by gene regulatory networks and cell–cell interactions. Nature methods 22, 982–993 (2025).

[17] Yuan, Q. & Duren, Z. Inferring gene regulatory networks from single-cell multiome data using atlas-scale external data. Nature Biotechnology 43, 247–257 (2025).

[18] Franzén, O.Gan, L.-M. & Björkegren, J. L. M. Panglaodb: a web server for exploration of mouse and human single-cell rna sequencing data. Database 2019 (2019). URL 10.1093/database/baz046.

[19] Hu, C. et al. Cellmarker 2.0: an updated database of manually curated cell markers in human/mouse and web tools based on scrna-seq data. Nucleic Acids Research 51, D870–D876 (2022). URL 10.1093/nar/gkac947.

[20] Tseng, T.-Y. et al. Transcriptional dynamics of cd8+ t-cell exhaustion in immune checkpoint inhibitor resistance at single-cell resolution. Molecular Cancer 24 (2025). URL 10.1186/s12943-025-02468-7.

[21] Wang, Z. et al. Geneagent: self-verification language agent for gene-set analysis using domain databases. Nature Methods 22, 1677–1685 (2025). URL 10.1038/s41592-025-02748-6.

[22] Kolberg, L. et al. g:profiler—interoperable web service for functional enrichment analysis and gene identifier mapping (2023 update). Nucleic Acids Research 51, W207–W212 (2023). URL 10.1093/nar/gkad347.

[23] Carrette, F. & Surh, C. D. Il-7 signaling and cd127 receptor regulation in the control of t cell homeostasis. Seminars in Immunology 24, 209–217 (2012). URL 10.1016/j.smim.2012.04.010.

[24] Richard, A. C. et al. T cell cytolytic capacity is independent of initial stimulation strength. Nature Immunology 19, 849–858 (2018). URL 10.1038/s41590-018-0160-9.

[25] Tillé, L. et al. Activation of the transcription factor nfat5 in the tumor microen-vironment enforces cd8+ t cell exhaustion. Nature Immunology 24, 1645–1653 (2023). URL 10.1038/s41590-023-01614-x.

[26] Lukhele, S. et al. The transcription factor irf2 drives interferon-mediated cd8+ t cell exhaustion to restrict anti-tumor immunity. Immunity 55, 2369–2385.e10 (2022). URL 10.1016/j.immuni.2022.10.020.

[27] Khan, O. et al. Tox transcriptionally and epigenetically programs cd8+ t cell exhaustion. Nature 571, 211–218 (2019). URL 10.1038/s41586-019-1325-x.

[28] Liu, L. et al. Bcl-2 expression promotes immunosuppression in chronic lymphocytic leukemia by enhancing regulatory t cell differentiation and cytotoxic t cell exhaustion. Molecular Cancer 21 (2022). URL 10.1186/s12943-022-01516-w.

[29] Virshup, I., Rybakov, S., Theis, F. J., Angerer, P. & Wolf, F. A. anndata: Access and store annotated data matrices. Journal of Open Source Software 9, 4371 (2024). URL 10.21105/joss.04371.

[30] Müller-Dott, S. et al. Expanding the coverage of regulons from high-confidence prior knowledge for accurate estimation of transcription factor activities. Nucleic Acids Res. 51, 10934–10949 (2023).

[31] Han, H. et al. Trrust v2: an expanded reference database of human and mouse transcriptional regulatory interactions. Nucleic Acids Research 46, D380–D386 (2017). URL 10.1093/nar/gkx1013.

[32] Heumos, L. et al. Best practices for single-cell analysis across modalities. Nature Reviews Genetics 24, 550–572 (2023). URL 10.1038/s41576-023-00586-w.

[33] Hu, Z., Przytycki, P. F. & Pollard, K. S. Cellwalker2: Multi-omic discovery using hierarchical cell type relationships. Cell Genomics 5, 100886 (2025). URL 10.1016/j.xgen.2025.100886.

[34] Paszke, A. et al. Pytorch: An imperative style, high-performance deep learning library (2019). URL https://arxiv.org/abs/1912.01703.

[35] Pratapa, A., Jalihal, A. P., Law, J. N., Bharadwaj, A. & Murali, T. M. Benchmarking algorithms for gene regulatory network inference from single-cell transcriptomic data. Nature Methods 17, 147–154 (2020). URL 10.1038/s41592-019-0690-6.

[36] Zinati, Y., Takiddeen, A. & Emad, A. Groundgan: Grn-guided simulation of single-cell rna-seq data using causal generative adversarial networks. Nature Communications 15 (2024). URL 10.1038/s41467-024-48516-6.

[37] Liu, T. et al. Cistrome: an integrative platform for transcriptional regulation studies. Genome biology 12, R83 (2011).

[38] De Simone, M. et al. A comprehensive analysis framework for evaluating commercial single-cell rna sequencing technologies. Nucleic Acids Research 53 (2024). URL 10.1093/nar/gkae1186.

[39] Wolock, S. L., Lopez, R. & Klein, A. M. Scrublet: Computational identification of cell doublets in single-cell transcriptomic data. Cell Systems 8, 281–291.e9 (2019). URL https://www.sciencedirect.com/science/article/pii/S2405471218304745.

[40] Domínguez Conde, C. et al. Cross-tissue immune cell analysis reveals tissue-specific features in humans. Science 376 (2022). URL 10.1126/science.abl5197.

[41] Badia-i Mompel, P. et al. decoupler: ensemble of computational methods to infer biological activities from omics data. Bioinformatics Advances 2 (2022). URL 10.1093/bioadv/vbac016.

