## Supplementary material for "Concurrent de novo inference of cell types and gene regulatory networks with NetMap": Netmap Supplement

### 1 Evaluation of Netmap configurations

In order to find a good default configuration and understand the effect of different hyperparameter choices and explainable AI methods, we performed a hyperparameter search using the synthetic data. While each configuration runs quite quickly on the smaller sized synthetic data the space of potential hyperparameters is quite extensive. We set a budget of 3 weeks to evaluate a range of architecture configurations, hyperparameters and 3 explainability methods (Deconvolution, GradientShap, GuidedBackprop) which exhibited feasible run times in a preliminary test. This results in 108 tested configurations which are summarized in table Table 1. The final test ran on 19 simulated datasets with varying size and number of clusters, but not all configurations produced a result on every dataset, due to the runtime restriction. We selected the configuration which performed the best on average in the on-target evaluation, which is presented in Figure 1. In Figure 2, we show the corresponding off-target values in the same order as for Figure 1. While we do recover a couple of edges from the 'wrong' clusters, we still see a better signal in the on-target than the off-target regime. Lastly, in Figure 3, we show that the top setups also achieve decent clustering results. We observe, that the xAI appears to be the most important factor in the performace, as almost all top-50 results use GradientShap in the explainablity step. While the top performers had a wider hidden layer and a shallower architecture, overall no striking trend can be found in our hyperparameter setup. We suspect that the data set characteristics play a more important role.

### 2 Supplementary Figures: Synthetic data

| **Hyperparameter** | **choices** |
|---|---|
| Loss function | Negative Bionomial Likelihood (NB), Zero Inflated NB |
| Encoder/Decoder Hidden Layer size | 16, 32, 64 |
| Latent dimensionality | 10 |
| Encoder/Decoder Hidden Layers | 1,2,3 |
| Feature Attribution method | GradientShap (GS), Deconvolution (DC), GuidedBackprop (GB) |
| Dropout | 0, 0.1 |
| Background for GS | zeros, randomized, data |

**Table 1** Parameter choices for the sweep

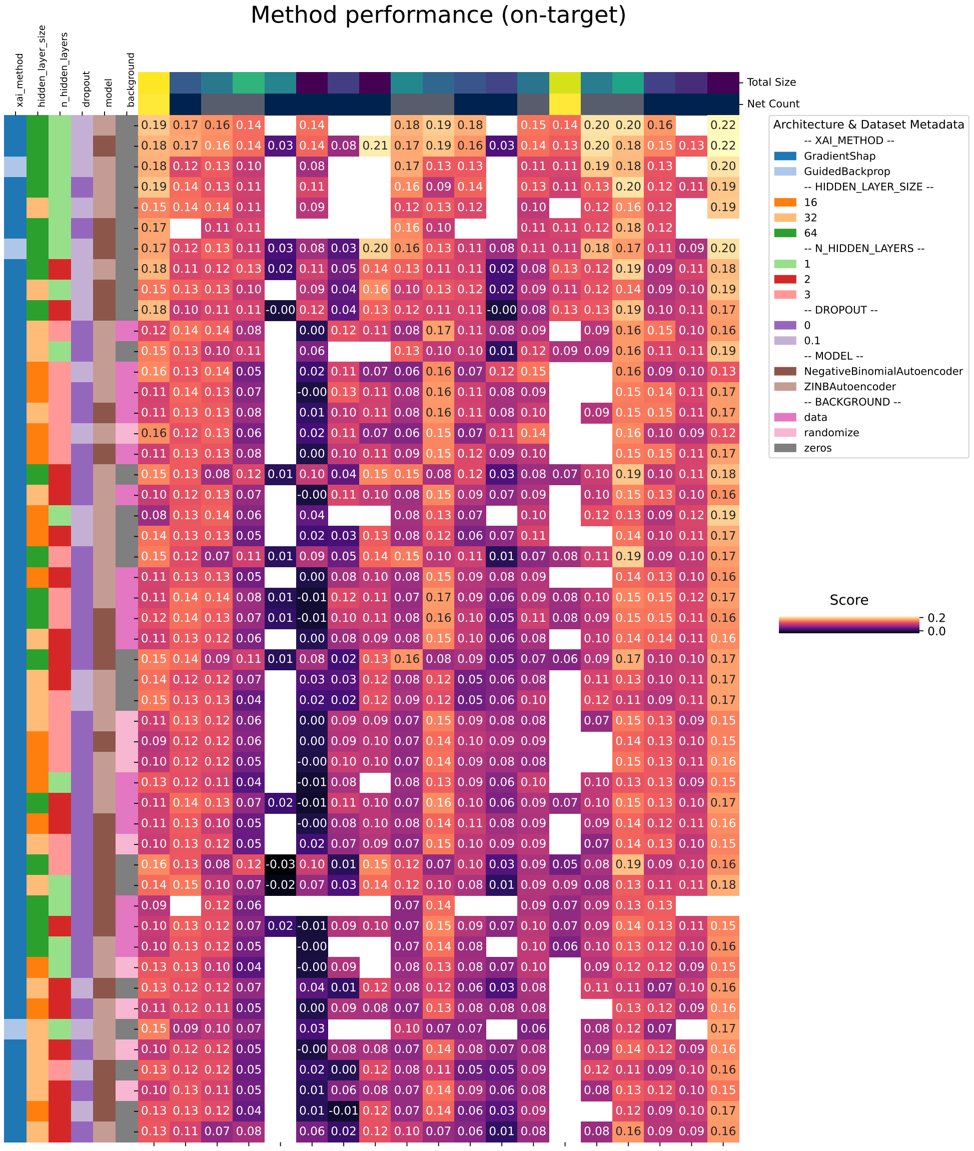


**Fig. 1** On-target recovery of the tested configuration ordered by the row-wise mean. The row-wise annotation shows the method configurations, the column annotation shows the number of networks (clusters) which where used in the test run (there is one background network identical to all clusters).

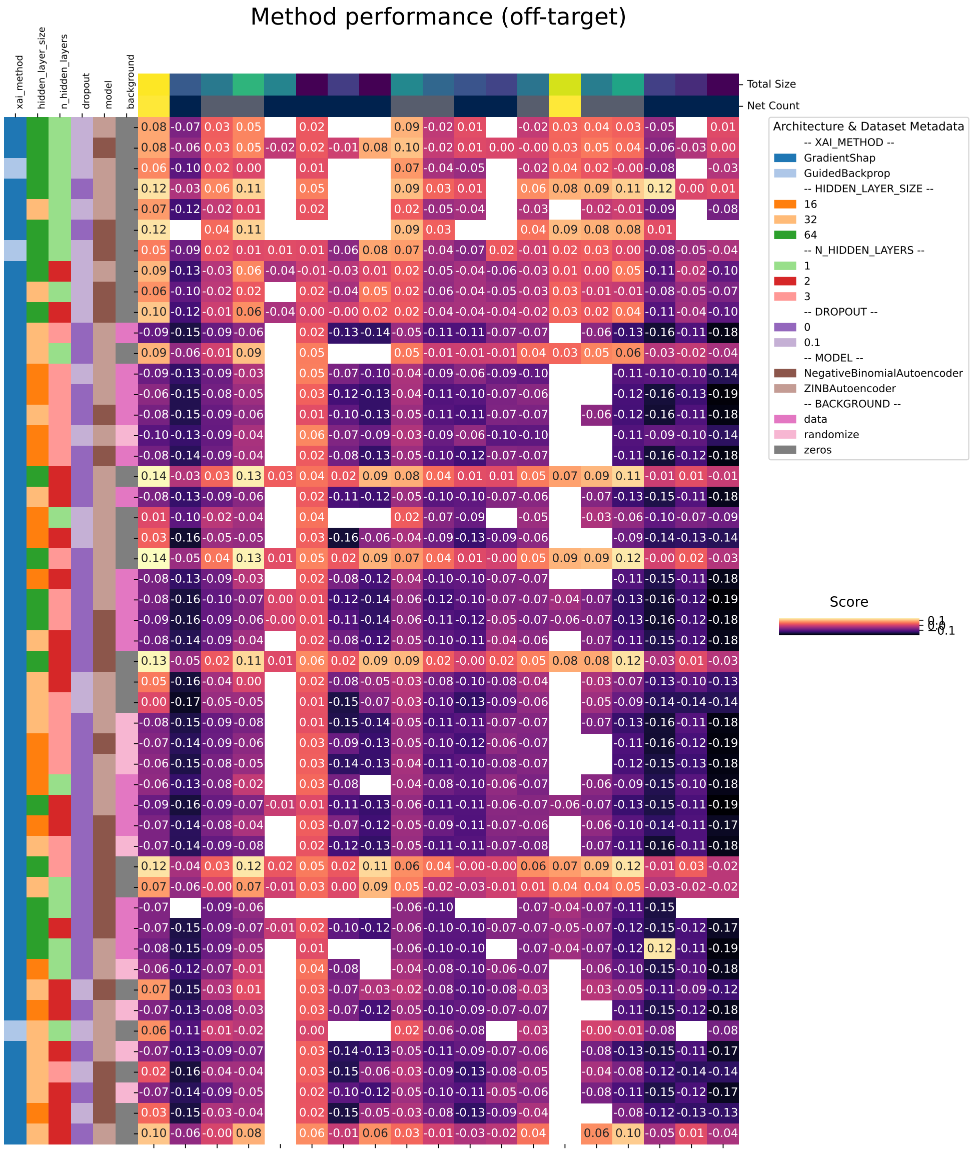


**Fig. 2** On-target recovery of the tested configuration. The order of the rows is identical to the order in Figure 1. The row-wise annotation shows the method configurations, the column annotation shows the number of networks (clusters) which where used in the test run (there is one background network identical to all clusters).

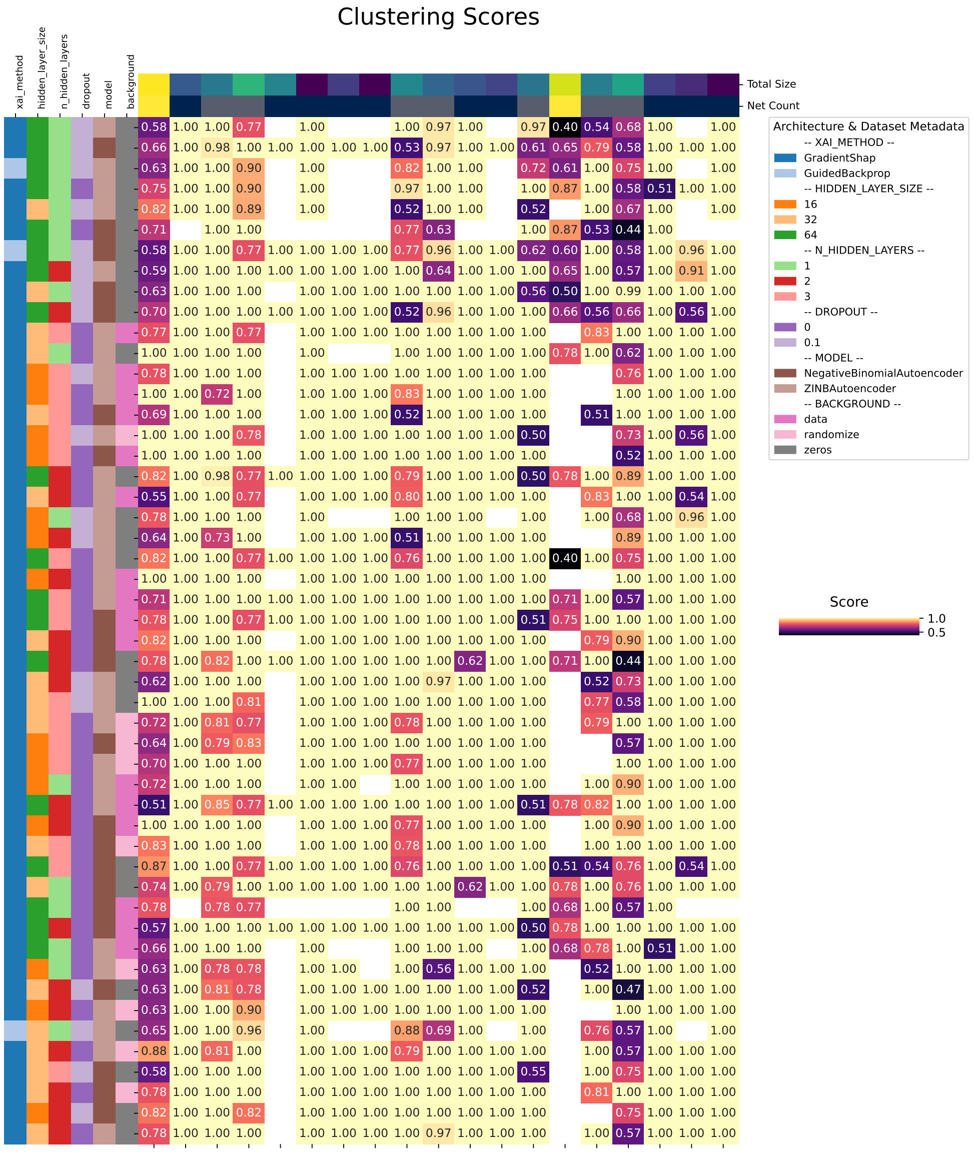


**Fig. 3** Clustering scores of the tested configuration. The order of the rows is identical to the order in Figure 1. The row-wise annotation shows the method configurations, the column annotation shows the number of networks (clusters) which where used in the test run (there is one background network identical to all clusters).

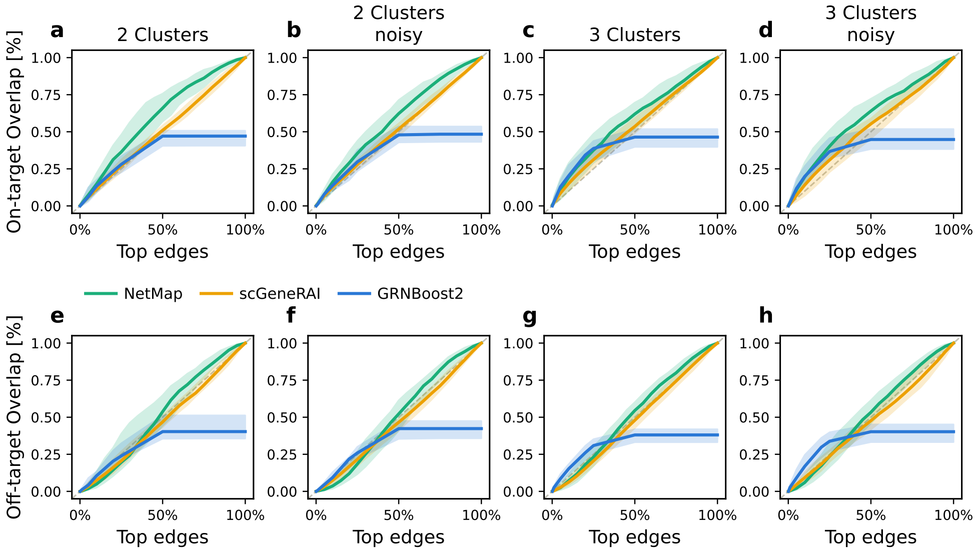


**Fig. 4** Performance on synthetic data split by data configuration

### 3 Supplementary Figures: PBMC

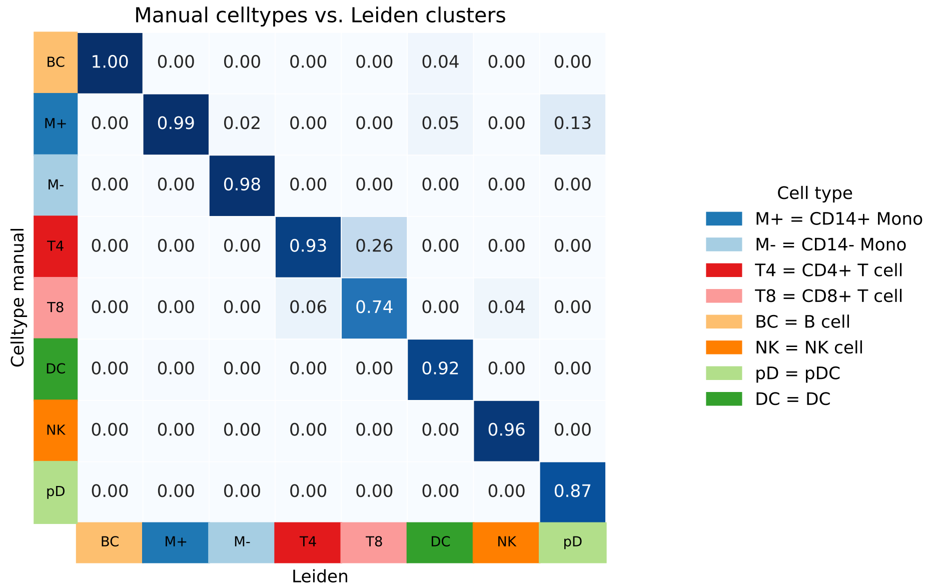


**Fig. 5** Comparison of the GRN clustering with the ground truth clustering. The clusters are mapped onto each other using LSA assignment. The values are averaged across the four datasets. Leiden clustering of the GRN is generally highly concordant with the leiden clustering of the gene expression data, except for some closely related cells switching categories.

**Table 2**: Cell Type Cluster Annotations and High-Level Groupings

| **Index** | **Cluster** | **Cluster Name** | **Cluster High Level** |
|---|---|---|---|
| Abnormal myeloid cell | 11 | Neutrophils | Neutrophils |
| Activated B cell | 2 | Activated B cells | B cells |
| Activated CD4+ T cell | 20 | Naive/Cytotoxic T cell | T cells |
| Activated CD8+ T cell | 23 | Activated T cells/ Effector T cells | T cells |
| Activated T cell | 16 | Activated/T helper cells | T cells |
| Activated memory B cell | 15 | Not specific | Not specific |
| Activated memory CD8+ T cell | 4 | Activated T cell | T cells |
| Activated memory T cell | 4 | Activated T cell | T cells |
| Activated naive CD8+ T cell | 4 | Activated T cell | T cells |
| Activated naive T cell | 4 | Activated T cell | T cells |
| Activated natural killer cell | 4 | Activated T cell | T cells |
| Age-associated B cell | 15 | Not specific | Not specific |
| Aged memory B cell | 2 | Activated B cells | B cells |
| Antibody Secreting B cell | 1 | Memory B cells | B cells |
| Astrocyte | 15 | Not specific | Not specific |
| Atypical memory B cell | 2 | Activated B cells | B cells |
| B cell | 3 | B cells/ Dendritic cells | B cells/ Dendritic cells |
| B cell lineage | 2 | Activated B cells | B cells |

Continued on next page

**Table 2 – Continued from previous page**

| Index | Cluster | Cluster Name | Cluster High Level |
|---|---|---|---|
| Basophil | 10 | Stem cell | Stem cell |
| Beta cell($\beta$ cell) | 15 | Not specific | Not specific |
| CAR-T cell | 15 | Not specific | Not specific |
| CD14 monocyte | 6 | Monocytes | Monocytes |
| CD14+ monocyte | 6 | Monocytes | Monocytes |
| CD16 monocyte | 6 | Monocytes | Monocytes |
| CD16+ monocyte | 6 | Monocytes | Monocytes |
| CD20+ B cell | 1 | Memory B cells | B cells |
| CD3/CD28-stimulated NK cell | 24 | T cells/ NK cells | T cells/ NK cells |
| CD3/CD28-stimulated T cell | 15 | Not specific | Not specific |
| CD4 T cell | 17 | Exhausted/proliferative T cells | T cells |
| CD4+ T cell | 22 | Effector T cells/ NK cells | T cells/ NK cells |
| CD4+ T helper cell | 18 | T cells (?) | T cells |
| CD4+ central memory like T (Tcm-like) cell | 20 | Naive/Cytotoxic T cell | T cells |
| CD4+ cytotoxic T lymphocyte | 15 | Not specific | Not specific |
| CD4+ cytotoxic T1 cell | 20 | Naive/Cytotoxic T cell | T cells |
| CD4+ cytotoxic T2 cell | 20 | Naive/Cytotoxic T cell | T cells |
| CD4+ recently activated effector memory or effector T cell (CTL) | 23 | Activated T cells/ Effector T cells | T cells |
| CD4+ tumor antigen-specific T (Tas) cell | 15 | Not specific | Not specific |
| CD56+ natural killer cell | 24 | T cells/ NK cells | T cells/ NK cells |
| CD8 T cell | 18 | T cells (?) | T cells |
| CD8+ T cell | 22 | Effector T cells/ NK cells | T cells/ NK cells |
| CD8+ mucosal-associated invariant T cell(CD8+ MAIT) | 22 | Effector T cells/ NK cells | T cells/ NK cells |
| CD8+ recently activated effector memory or effector T cell (CTL) | 23 | Activated T cells/ Effector T cells | T cells |
| CD8+ terminally differentiated effector memory re-expressing CD45RA T cell(CD8+ TEMRA) | 16 | Activated/T helper cells | T cells |
| CD8+ tumor antigen-specific T (Tas) cell | 15 | Not specific | Not specific |
| CD8+ intraepithelial cell | 15 | Not specific | Not specific |
| Cancer cell | 22 | Effector T cells/ NK cells | T cells/ NK cells |
| Cancer stem cell | 15 | Not specific | Not specific |
| Central memory CD4+ T cell | 19 | Central memory T cells | T cells |
| Central memory CD8+ T cell | 19 | Central memory T cells | T cells |
| Central memory T cell | 22 | Effector T cells/ NK cells | T cells/ NK cells |
| Cholangiocyte | 15 | Not specific | Not specific |
| Circulating memory T cell | 13 | Naive circulating T cells | T cells |
| Classical monocyte | 6 | Monocytes | Monocytes |
| Conventional T(Tconv) cell | 14 | T reg cells | T cells |
| Conventional dendritic cell 1(cDC1) | 15 | Not specific | Not specific |
| Conventional dendritic cell 2(cDC2) | 15 | Not specific | Not specific |
| Cycling cell | 8 | Cycling cells | Cycling cells |
| Cytotoxic CD4+ T cell | 21 | Cytotoxic T cell | T cells |
| Cytotoxic CD4+ T2 cell | 21 | Cytotoxic T cell | T cells |
| Cytotoxic CD8 T cell | 21 | Cytotoxic T cell | T cells |
| Cytotoxic CD8+ T cell | 24 | T cells/ NK cells | T cells/ NK cells |
| Cytotoxic T cell | 23 | Activated T cells/ Effector T cells | T cells |
| Dendritic cell | 3 | B cells/ Dendritic cells | B cells/ Dendritic cells |
| Dendritic cell lineage | 15 | Not specific | Not specific |
| Dermal cell | 15 | Not specific | Not specific |
| Dividing plasma B cell | 3 | B cells/ Dendritic cells | B cells/ Dendritic cells |
| Early activated CD4+ T cell | 16 | Activated/T helper cells | T cells |
| Early activated CD8+ T cell | 16 | Activated/T helper cells | T cells |
| Early effector T cell | 18 | T cells (?) | T cells |
| Effector CD4+ T cell | 23 | Activated T cells/ Effector T cells | T cells |
| Effector CD8 T cell | 24 | T cells/ NK cells | T cells/ NK cells |
| Effector CD8+ T cell | 23 | Activated T cells/ Effector T cells | T cells |
| Effector T(Teff) cell | 7 | Effector T cell | T cells |
| Effector memory CD4 T cell | 22 | Effector T cells/ NK cells | T cells/ NK cells |
| Effector memory CD4+ T cell | 22 | Effector T cells/ NK cells | T cells/ NK cells |
| Effector memory CD8 T cell | 7 | Effector T cell | T cells |
| Effector memory CD8+ T cell | 22 | Effector T cells/ NK cells | T cells/ NK cells |
| Effector memory T cell | 22 | Effector T cells/ NK cells | T cells/ NK cells |

Continued on next page

**Table 2 – Continued from previous page**

| Index | Cluster | Cluster Name | Cluster High Level |
|---|---|---|---|
| Effector regulatory T(eTreg) cell | 24 | T cells/ NK cells | T cells/ NK cells |
| Endothelial cell | 15 | Not specific | Not specific |
| Epithelial cell | 15 | Not specific | Not specific |
| Erythroid cell | 15 | Not specific | Not specific |
| Exhausted CD4+ T cell | 17 | Exhausted/proliferative T cells | T cells |
| Exhausted CD8+ T cell | 17 | Exhausted/proliferative T cells | T cells |
| Fibroblast | 15 | Not specific | Not specific |
| Finally highly effector (TEMRA) memory T cell | 24 | T cells/ NK cells | T cells/ NK cells |
| Follicular B cell | 15 | Not specific | Not specific |
| Follicular regulatory T(Tfr) cell | 15 | Not specific | Not specific |
| Gamma delta($\gamma\delta$) T cell | 22 | Effector T cells/ NK cells | T cells/ NK cells |
| Germinal center B cell | 15 | Not specific | Not specific |
| Granulocyte | 11 | Neutrophils | Neutrophils |
| Hematopoietic progenitor cell | 15 | Not specific | Not specific |
| Hematopoietic stem cell | 10 | Stem cell | Stem cell |
| IgA memory B cell | 1 | Memory B cells | B cells |
| IgG memory B cell | 1 | Memory B cells | B cells |
| IgM memory B cell | 1 | Memory B cells | B cells |
| Immature B cell | 3 | B cells/ Dendritic cells | B cells/ Dendritic cells |
| Immature erythroid cell | 15 | Not specific | Not specific |
| Immature neutrophil | 5 | Neutrophils | Neutrophils |
| Intermediate monocyte | 6 | Monocytes | Monocytes |
| Intermediate monocyte cell | 6 | Monocytes | Monocytes |
| Intermediate transition memory B cell | 1 | Memory B cells | B cells |
| Leukocyte | 12 | Myeloid cells | Myeloid cells |
| Lymphocyte | 24 | T cells/ NK cells | T cells/ NK cells |
| Lymphoid cell | 16 | Activated/T helper cells | T cells |
| M1 macrophage | 15 | Not specific | Not specific |
| M2 macrophage | 15 | Not specific | Not specific |
| Macrophage | 6 | Monocytes | Monocytes |
| Malignant cell | 15 | Not specific | Not specific |
| Mast cell | 15 | Not specific | Not specific |
| Mature B cell | 3 | B cells/ Dendritic cells | B cells/ Dendritic cells |
| Mature dendritic cell | 6 | Monocytes | Monocytes |
| Mature neutrophil | 5 | Neutrophils | Neutrophils |
| Mature progenitor cell | 3 | B cells/ Dendritic cells | B cells/ Dendritic cells |
| Megakaryocyte | 9 | Megakaryocytes | Megakaryocytes |
| Memory B cell | 3 | B cells/ Dendritic cells | B cells/ Dendritic cells |
| Memory CD4+ T cell | 22 | Effector T cells/ NK cells | T cells/ NK cells |
| Memory CD8 T cell | 18 | T cells (?) | T cells |
| Memory CD8+ T cell | 23 | Activated T cells/ Effector T cells | T cells |
| Memory T cell | 22 | Effector T cells/ NK cells | T cells/ NK cells |
| Memory-like B cell | 1 | Memory B cells | B cells |
| Mesenchymal cell | 15 | Not specific | Not specific |
| Microglial cell | 15 | Not specific | Not specific |
| Monocyte | 6 | Monocytes | Monocytes |
| Monocyte derived dendritic cell | 6 | Monocytes | Monocytes |
| Monocyte precursor | 15 | Not specific | Not specific |
| Monocyte progenitor cell | 6 | Monocytes | Monocytes |
| Mucosa-associated invariant T (MAIT) cell | 22 | Effector T cells/ NK cells | T cells/ NK cells |
| Myeloid cell | 6 | Monocytes | Monocytes |
| Myeloid dendritic cell | 15 | Not specific | Not specific |
| Myeloid dendritic cell 1 | 12 | Myeloid cells | Myeloid cells |
| Myeloid derived suppressor cell (MDSC) | 15 | Not specific | Not specific |
| Myeloid progenitor cell | 15 | Not specific | Not specific |
| Myofibroblast | 15 | Not specific | Not specific |
| Naive B cell | 3 | B cells/ Dendritic cells | B cells/ Dendritic cells |
| Naive CD4 T cell | 19 | Central memory T cells | T cells |
| Naive CD4+ T cell | 20 | Naive/Cytotoxic T cell | T cells |
| Naive CD8 T cell | 13 | Naive circulating T cells | T cells |
| Naive CD8+ T cell | 20 | Naive/Cytotoxic T cell | T cells |
| Naive T(Th0) cell | 22 | Effector T cells/ NK cells | T cells/ NK cells |
| Naive cytotoxic T cell | 18 | T cells (?) | T cells |
| Natural killer T(NKT) cell | 22 | Effector T cells/ NK cells | T cells/ NK cells |
| Natural killer cell | 22 | Effector T cells/ NK cells | T cells/ NK cells |
| Natural regulatory T cell | 15 | Not specific | Not specific |

Continued on next page

**Table 2 – Continued from previous page**

| Index | Cluster | Cluster Name | Cluster High Level |
|---|---|---|---|
| Neutrophil | 11 | Neutrophils | Neutrophils |
| Neutrophil lineage | 15 | Not specific | Not specific |
| Neutrophil progenitor cell | 11 | Neutrophils | Neutrophils |
| Non-classical monocyte | 6 | Monocytes | Monocytes |
| Pan-B cell | 2 | Activated B cells | B cells |
| Pan-T cell | 16 | Activated/T helper cells | T cells |
| Patelet | 9 | Megakaryocytes | Megakaryocytes |
| Peripheral blood mononuclear cell (PBMC) | 2 | Activated B cells | B cells |
| Peripheral immune cell | 12 | Myeloid cells | Myeloid cells |
| Plasma cell | 3 | B cells/ Dendritic cells | B cells/ Dendritic cells |
| Plasmablast | 3 | B cells/ Dendritic cells | B cells/ Dendritic cells |
| Plasmacytoid dendritic cell(pDC) | 3 | B cells/ Dendritic cells | B cells/ Dendritic cells |
| Platelet | 9 | Megakaryocytes | Megakaryocytes |
| Polymorphonuclear myeloid-derived suppressor(PMN-MDSC) cell | 11 | Neutrophils | Neutrophils |
| Pre-neutrophil | 5 | Neutrophils | Neutrophils |
| Precursor cell | 10 | Stem cell | Stem cell |
| Precursor memory cell | 15 | Not specific | Not specific |
| Pro-inflammatory macrophage | 15 | Not specific | Not specific |
| Pro-neutrophil | 15 | Not specific | Not specific |
| Pro-tumor type-2 pericyte | 15 | Not specific | Not specific |
| Progenitor cell | 10 | Stem cell | Stem cell |
| Proliferative CD4+ T cell | 17 | Exhausted/proliferative T cells | T cells |
| Proliferative CD8+ T cell | 17 | Exhausted/proliferative T cells | T cells |
| Proliferative T cell | 18 | T cells (?) | T cells |
| Proliferative cell | 8 | Cycling cells | Cycling cells |
| Proliferative lymphocyte | 8 | Cycling cells | Cycling cells |
| Red blood cell (erythrocyte) | 15 | Not specific | Not specific |
| Regulatory B(Breg) cell | 14 | T reg cells | T cells |
| Regulatory CD4 T cell | 14 | T reg cells | T cells |
| Regulatory CD4+ T cell | 17 | Exhausted/proliferative T cells | T cells |
| Regulatory CD8+ T cell | 18 | T cells (?) | T cells |
| Regulatory T(Treg) cell | 15 | Not specific | Not specific |
| Responding conventional T cell | 24 | T cells/ NK cells | T cells/ NK cells |
| Revertant memory CD8+ T(TEMRA) cell | 16 | Activated/T helper cells | T cells |
| Smooth muscle cell | 15 | Not specific | Not specific |
| Stem cell | 10 | Stem cell | Stem cell |
| T cell | 22 | Effector T cells/ NK cells | T cells/ NK cells |
| T cell large granular lymphocytic leukemia cell | 24 | T cells/ NK cells | T cells/ NK cells |
| T follicular helper(Tfh) cell | 16 | Activated/T helper cells | T cells |
| T helper 1(Th1) cell | 24 | T cells/ NK cells | T cells/ NK cells |
| T helper 17(Th17) cell | 22 | Effector T cells/ NK cells | T cells/ NK cells |
| T helper 2(Th2) cell | 22 | Effector T cells/ NK cells | T cells/ NK cells |
| T helper 9(Th9) cell | 16 | Activated/T helper cells | T cells |
| T helper(Th) cell | 22 | Effector T cells/ NK cells | T cells/ NK cells |
| Terminal effector CD8+ T cell | 21 | Cytotoxic T cell | T cells |
| Tissue resident macrophage | 15 | Not specific | Not specific |
| Tissue resident memory T(TRM) cell | 12 | Myeloid cells | Myeloid cells |
| Transitional B cell | 3 | B cells/ Dendritic cells | B cells/ Dendritic cells |
| Transitional CD8+ T cell | 7 | Effector T cell | T cells |
| Tumor regulatory T cell | 14 | T reg cells | T cells |
| Yolk sac-derived myeloid-biased progenitor cell(YSMP) | 12 | Myeloid cells | Myeloid cells |

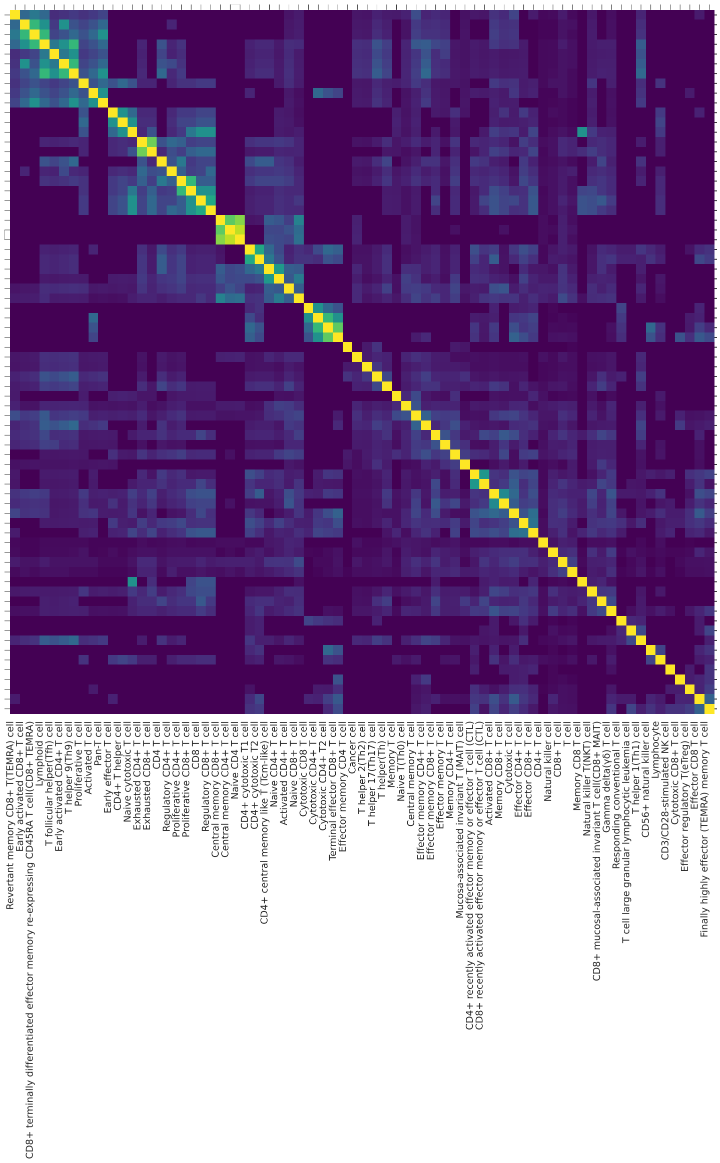


**Fig. 6** Clustered Heatmap of the Jaccard similarities between the marker genes sets of the cellmarker database. This group broadly represents the T-cell like immune cells. The heatmap is symmetrical and the dendrograms row labels have been removed for readability purposes. Generally speaking, the jaccard similarities indicate some commonalities between the definitions, however, the clusters are not strictly confined to the diagnoal, indiating that there are inconsistent definitions. This serves to illustrate, that, while the sets may indicate a general biological theme, they are unsuited to pinpoint exactly the function of the clusters without further investigation.

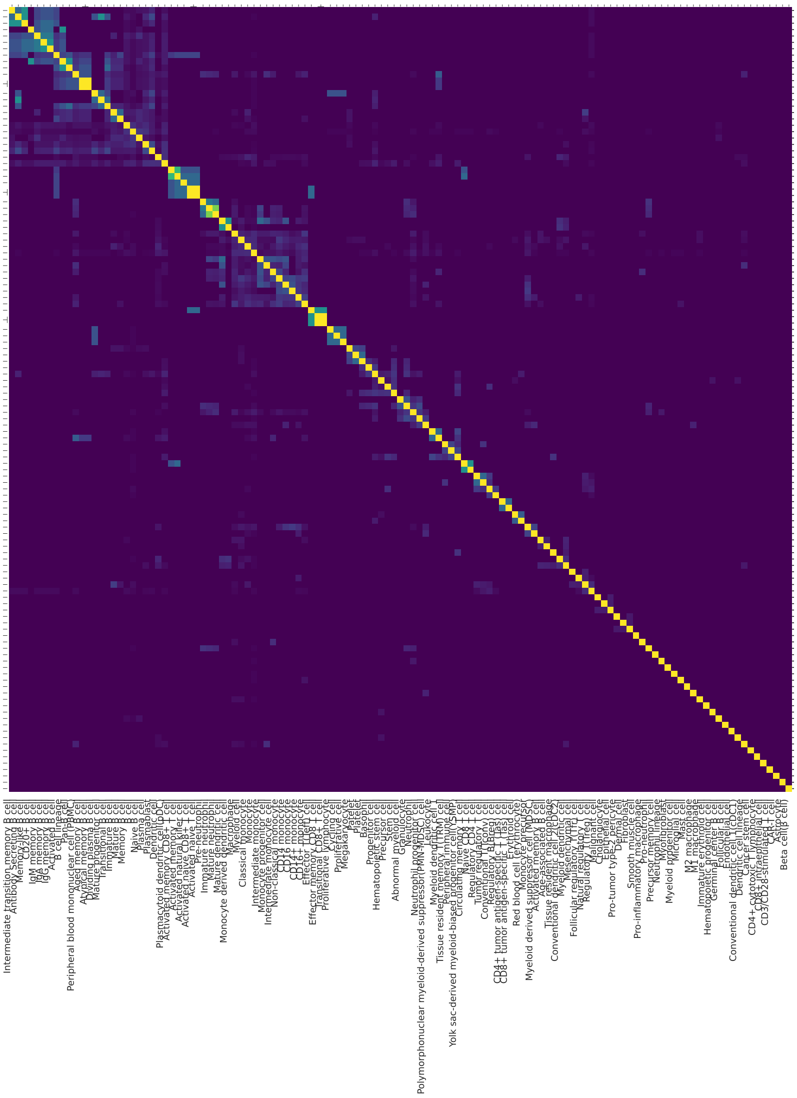

**Fig. 7** Clustered Heatmap of the Jaccard similarities between the marker genes sets of the cellmarker database. This group broadly represents the non-T-cell like immune cells. The heatmap is symmetrical and the dendrograms row labels have been removed for readability purposes. Generally speaking, the jaccard similarity of the gene sets tends to be low even for cell types which by their cluster titel indicate a similar cell type. Some T-cell like groups are mixed into the 'non-T-cell' like set as well. This serves to illustrate, that, while the sets may indicate a general biological theme, they are unsuited to pinpoint exactly the function of the clusters without further investigation.

### 4 CD8+ Tcells

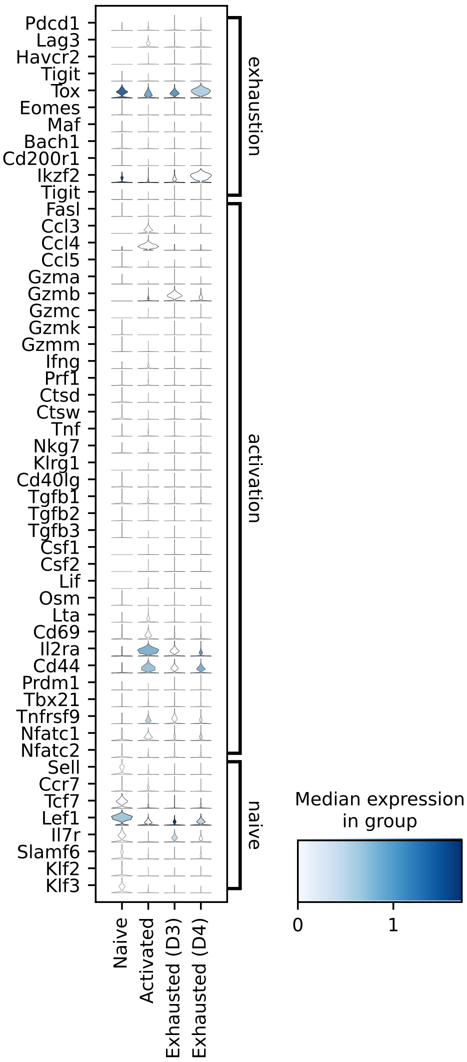


**Fig. 8** Expression of marker genes in mouse gene expression data split by condition (Naive, Activated, Exhausted (D3) and Exhausted (D4)

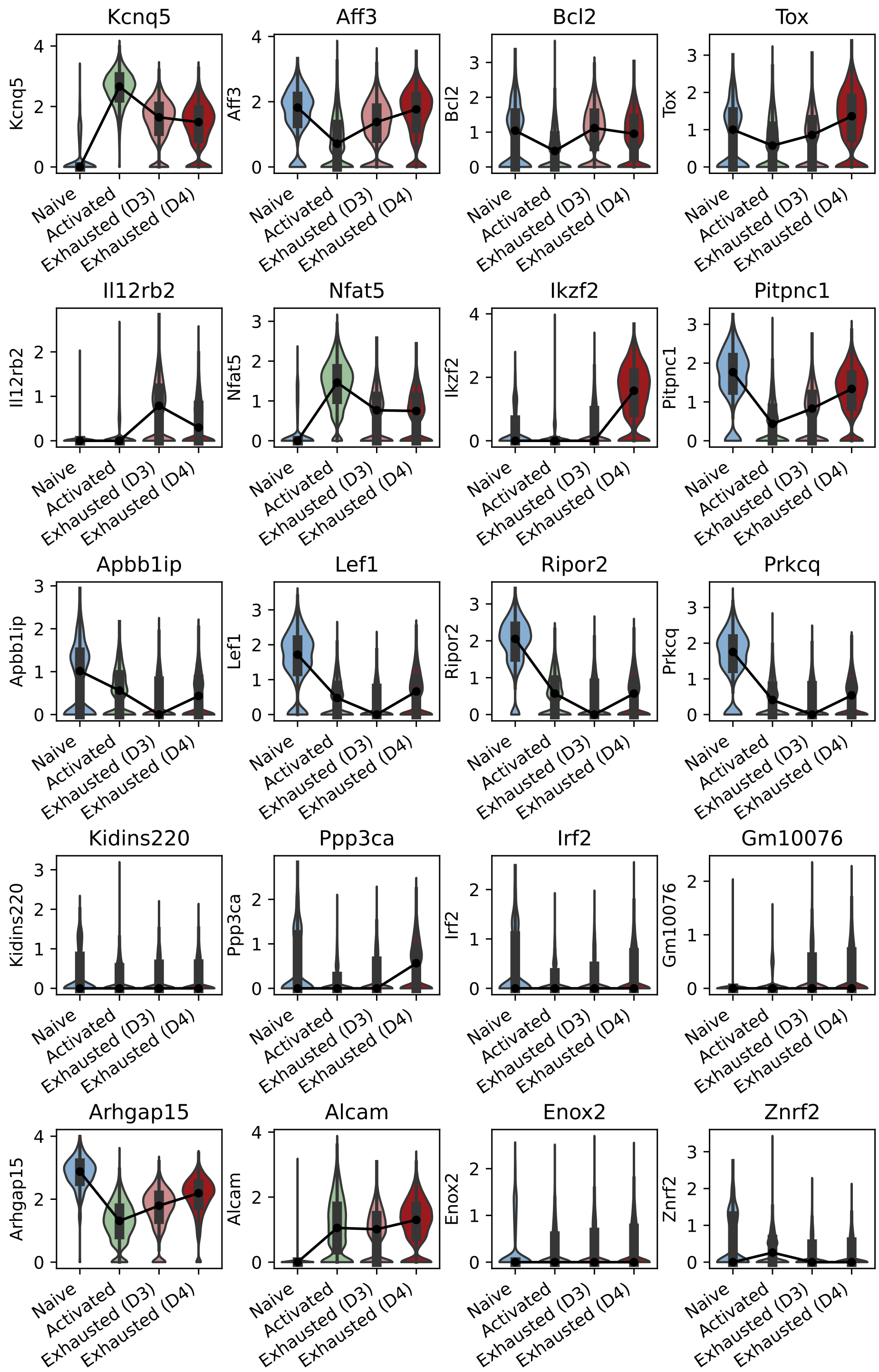


**Fig. 9** Gene expression distribution of the top 20 genes by PR centrality grouped by condition.

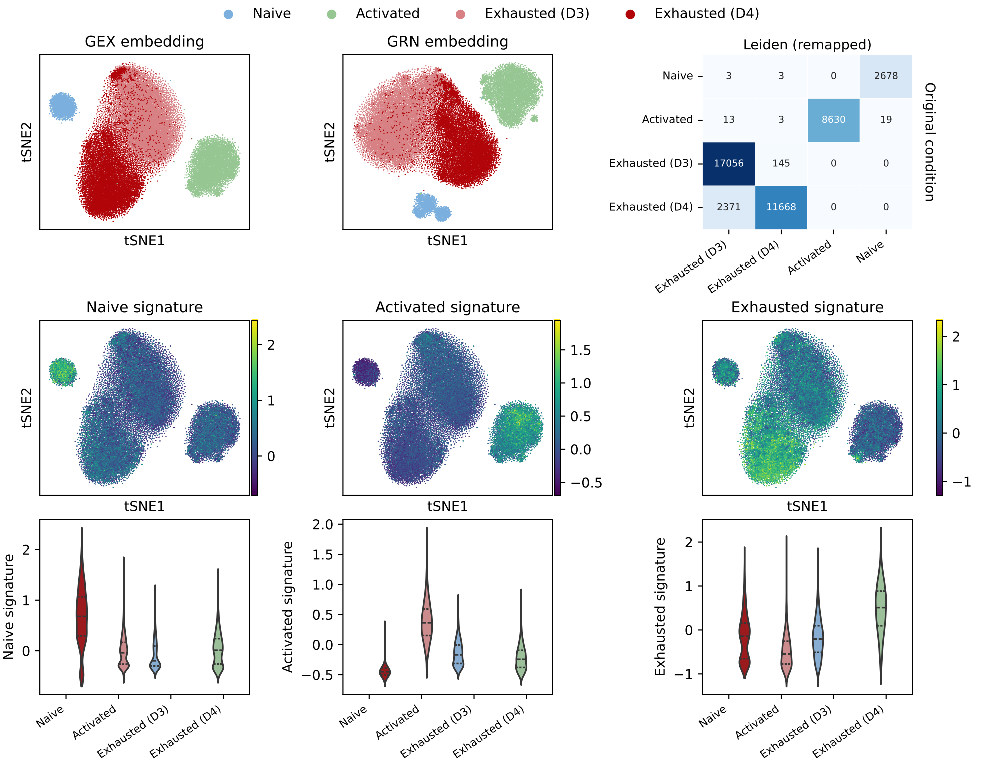


**Fig. 10** Analysis of CD8+ T gene expression data. A Gene expression embedding and clustering which has been produced using a standard workflow including QC, scaling, PCA, kNN embedding, Leiden clustering followed by tSNE embedding. B Embedding of the GRN for comparison, where we took the Netmap embedding of the data and used PCA, leiden and tSNE to produce the plot. C Clustering concordance of the Gene expression based Leiden clustering with the ground truth labels of the data. Most cells are assigned to the experimental day, except for cells from D3 and D4 of the experiment which are not all assigned, due to the stronger overlap between the gene expression clusters. D-F expression based scores computed from marker gene signatures. G-K Violin plots of the scores by day. We observe that the naive cluster has the strongest expression of the naive signature, the activated cluster the strongest expression of the activation signature, with a gradual decrease of the activation signature in D3 and D4. Conversely, we observe a gradual increase of the Exhausted signature upon activation.

**Table 3** Netmap Activated

| Term name | GO Term ID | P-value |
|---|---|---|
| Bcl2 homodimer complex | CORUM:1988 | 0.05 |
| immune system process | GO:0002376 | 0.00 |
| alpha-beta T cell differentiation | GO:0046632 | 0.00 |
| T cell activation | GO:0042110 | 0.00 |
| leukocyte activation | GO:0045321 | 0.00 |
| alpha-beta T cell activation | GO:0046631 | 0.00 |
| CD8-positive, alpha-beta T cell differentiation | GO:0043374 | 0.00 |
| cell activation | GO:0001775 | 0.00 |
| CD8-positive, alpha-beta T cell lineage commitment | GO:0043375 | 0.00 |
| lymphocyte activation | GO:0046649 | 0.00 |
| CD8-positive, alpha-beta T cell activation | GO:0036037 | 0.00 |
| cell morphogenesis | GO:0000902 | 0.00 |
| T cell differentiation | GO:0030217 | 0.00 |
| leukocyte differentiation | GO:0002521 | 0.01 |
| anatomical structure morphogenesis | GO:0009653 | 0.01 |
| somatic diversification of T cell receptor genes | GO:0002568 | 0.01 |
| T cell receptor V(D)J recombination | GO:0033153 | 0.01 |
| somatic recombination of T cell receptor gene segments | GO:0002681 | 0.01 |
| hemopoiesis | GO:0030097 | 0.01 |
| regulation of hemopoiesis | GO:1903706 | 0.01 |
| lymphocyte differentiation | GO:0030098 | 0.02 |
| regulation of cell development | GO:0060284 | 0.02 |
| immune system development | GO:0002520 | 0.03 |
| immune response | GO:0006955 | 0.03 |
| mononuclear cell differentiation | GO:1903131 | 0.04 |
| cell migration | GO:0016477 | 0.05 |
| molecular function regulator activity | GO:0098772 | 0.03 |
| channel regulator activity | GO:0016247 | 0.04 |
| transporter regulator activity | GO:0141108 | 0.05 |
| Colorectal cancer | KEGG:05210 | 0.01 |
| Prostate cancer | KEGG:05215 | 0.02 |
| Cholinergic synapse | KEGG:04725 | 0.02 |
| Parathyroid hormone synthesis, secretion and action | KEGG:04928 | 0.02 |
| Gastric cancer | KEGG:05226 | 0.04 |
| RUNX3 regulates WNT signaling | REAC:R-MMU-8951430 | 0.01 |
| Regulation of MITF-M-dependent genes involved in cell cycle and proliferation | REAC:R-MMU-9825892 | 0.01 |
| Repression of WNT target genes | REAC:R-MMU-4641265 | 0.02 |
| MITF-M-dependent gene expression | REAC:R-MMU-9856651 | 0.05 |
| Comprehensive IL 17A signaling | WP:WP5242 | 0.03 |

**Table 4** Netmap Naive

| Term name | GO Term ID | P-value |
|---|---|---|
| Bcl2 homodimer complex | CORUM:1988 | 0.05 |
| positive regulation of immune system process | GO:0002684 | 0.00 |
| regulation of immune system process | GO:0002682 | 0.00 |
| immune system process | GO:0002376 | 0.00 |
| CD8-positive, alpha-beta T cell lineage commitment | GO:0043375 | 0.00 |
| immune response | GO:0006955 | 0.00 |
| T cell activation | GO:0042110 | 0.00 |
| positive regulation of lymphocyte activation | GO:0051251 | 0.00 |
| leukocyte activation | GO:0045321 | 0.00 |
| positive regulation of leukocyte activation | GO:0002696 | 0.00 |
| positive regulation of cell activation | GO:0050867 | 0.00 |
| positive regulation of response to stimulus | GO:0048584 | 0.00 |
| regulation of response to stimulus | GO:0048583 | 0.00 |
| leukocyte differentiation | GO:0002521 | 0.00 |
| cell activation | GO:0001775 | 0.00 |
| regulation of cell adhesion | GO:0030155 | 0.01 |
| positive regulation of cell adhesion | GO:0045785 | 0.01 |
| positive regulation of leukocyte differentiation | GO:1902107 | 0.01 |
| positive regulation of hemopoiesis | GO:1903708 | 0.01 |
| lymphocyte activation | GO:0046649 | 0.01 |
| positive regulation of cell development | GO:0010720 | 0.01 |
| regulation of immune response | GO:0050776 | 0.01 |
| regulation of lymphocyte activation | GO:0051249 | 0.01 |
| immune system development | GO:0002520 | 0.01 |
| developmental process | GO:0032502 | 0.02 |
| positive regulation of leukocyte cell-cell adhesion | GO:1903039 | 0.02 |
| mononuclear cell differentiation | GO:1903131 | 0.02 |
| regulation of leukocyte activation | GO:0002694 | 0.02 |
| regulation of cell activation | GO:0050865 | 0.04 |
| positive regulation of cell-cell adhesion | GO:0022409 | 0.04 |
| B cell proliferation | GO:0042100 | 0.04 |
| lymphocyte proliferation | GO:0046651 | 0.04 |
| T cell differentiation in thymus | GO:0033077 | 0.04 |
| hemopoiesis | GO:0030097 | 0.05 |
| mononuclear cell proliferation | GO:0032943 | 0.05 |
| adaptive immune response based on somatic recombination of immune receptors built from immunoglobulin superfamily domains | GO:0002460 | 0.05 |
| Wnt signaling pathway | KEGG:04310 | 0.03 |
| Herpes simplex virus 1 infection | KEGG:05168 | 0.04 |
| Kaposi sarcoma-associated herpesvirus infection | KEGG:05167 | 0.05 |
| Factor: FKLF; motif: BGGGNGGVMD | TF:M01837 | 0.01 |

**Table 5** Netmap Exhausted (D3)

| Term name | GO Term ID | P-value |
|---|---|---|
| Bcl2 homodimer complex | CORUM:1988 | 0.05 |
| T cell activation | GO:0042110 | 0.00 |
| regulation of cell adhesion | GO:0030155 | 0.00 |
| CD8-positive, alpha-beta T cell lineage commitment | GO:0043375 | 0.00 |
| regulation of lymphocyte activation | GO:0051249 | 0.00 |
| lymphocyte activation | GO:0046649 | 0.00 |
| cell-cell adhesion | GO:0098609 | 0.00 |
| regulation of leukocyte activation | GO:0002694 | 0.00 |
| positive regulation of lymphocyte activation | GO:0051251 | 0.00 |
| cell adhesion | GO:0007155 | 0.00 |
| regulation of cell activation | GO:0050865 | 0.00 |
| leukocyte activation | GO:0045321 | 0.00 |
| positive regulation of leukocyte activation | GO:0002696 | 0.00 |
| cell morphogenesis | GO:0000902 | 0.00 |
| regulation of leukocyte cell-cell adhesion | GO:1903037 | 0.01 |
| positive regulation of cell activation | GO:0050867 | 0.01 |
| regulation of T cell activation | GO:0050863 | 0.01 |
| positive regulation of immune system process | GO:0002684 | 0.01 |
| leukocyte cell-cell adhesion | GO:0007159 | 0.01 |
| cell activation | GO:0001775 | 0.01 |
| alpha-beta T cell activation | GO:0046631 | 0.02 |
| positive regulation of cell adhesion | GO:0045785 | 0.02 |
| regulation of cell-cell adhesion | GO:0022407 | 0.02 |
| positive regulation of leukocyte cell-cell adhesion | GO:1903039 | 0.03 |
| Th1 and Th2 cell differentiation | KEGG:04658 | 0.00 |
| Factor: FPM315; motif: SRGGGAGGAGGN | TF:M01587 | 0.01 |
| B cell receptor signaling pathway | WP:WP274 | 0.03 |

**Table 6**: Netmap Exhausted (D4)

| Term name | GO Term ID | P-value |
|---|---|---|
| Bcl2 homodimer complex | CORUM:1988 | 0.05 |
| response to cytokine | GO:0034097 | 0.00 |
| lymphocyte activation | GO:0046649 | 0.00 |
| response to peptide | GO:1901652 | 0.00 |
| positive regulation of transcription by RNA polymerase II | GO:0045944 | 0.00 |
| T cell activation | GO:0042110 | 0.00 |
| leukocyte activation | GO:0045321 | 0.00 |
| positive regulation of macromolecule biosynthetic process | GO:0010557 | 0.00 |
| positive regulation of immune system process | GO:0002684 | 0.00 |
| positive regulation of biosynthetic process | GO:0009891 | 0.00 |
| CD8-positive, alpha-beta T cell lineage commitment | GO:0043375 | 0.00 |
| immune system development | GO:0002520 | 0.00 |
| cell activation | GO:0001775 | 0.00 |
| positive regulation of DNA-templated transcription | GO:0045893 | 0.00 |
| positive regulation of RNA biosynthetic process | GO:1902680 | 0.00 |
| positive regulation of RNA metabolic process | GO:0051254 | 0.00 |
| positive regulation of macromolecule metabolic process | GO:0010604 | 0.00 |
| hematopoietic or lymphoid organ development | GO:0048534 | 0.00 |
| alpha-beta T cell differentiation | GO:0046632 | 0.01 |
| positive regulation of cellular process | GO:0048522 | 0.01 |
| positive regulation of metabolic process | GO:0009893 | 0.01 |
| regulation of immune system process | GO:0002682 | 0.01 |
| leukocyte differentiation | GO:0002521 | 0.01 |
| cellular response to cytokine stimulus | GO:0071345 | 0.01 |
| regulation of transcription by RNA polymerase II | GO:0006357 | 0.01 |
| immune system process | GO:0002376 | 0.01 |
| G1/S transition of mitotic cell cycle | GO:0000082 | 0.01 |
| positive regulation of biological process | GO:0048518 | 0.01 |
| regulation of gene expression | GO:0010468 | 0.01 |
| positive regulation of response to stimulus | GO:0048584 | 0.02 |
| transcription by RNA polymerase II | GO:0006366 | 0.02 |
| regulation of cell adhesion | GO:0030155 | 0.02 |
| regulation of macromolecule biosynthetic process | GO:0010556 | 0.02 |
| alpha-beta T cell activation | GO:0046631 | 0.02 |
| cell cycle G1/S phase transition | GO:0044843 | 0.02 |
| lymphocyte differentiation | GO:0030098 | 0.02 |
| regulation of biosynthetic process | GO:0009889 | 0.02 |
| regulation of lymphocyte activation | GO:0051249 | 0.03 |
| positive regulation of cell differentiation | GO:0045597 | 0.05 |
| regulation of DNA-templated transcription | GO:0006355 | 0.05 |
| regulation of RNA biosynthetic process | GO:2001141 | 0.05 |

Continued on next page

Table 6 – continued from previous page

| Term name | GO Term ID | P-value |
|---|---|---|
| cytosol | GO:0005829 | 0.00 |
| DNA-binding transcription factor activity | GO:0003700 | 0.00 |
| DNA-binding transcription activator activity, RNA polymerase II-specific | GO:0001228 | 0.00 |
| DNA-binding transcription activator activity | GO:0001216 | 0.01 |
| DNA binding | GO:0003677 | 0.01 |
| transcription regulator activity | GO:0140110 | 0.01 |
| double-stranded DNA binding | GO:0003690 | 0.03 |
| RNA polymerase II transcription regulatory region sequence-specific DNA binding | GO:0000977 | 0.04 |
| sequence-specific DNA binding | GO:0043565 | 0.04 |
| Inflammatory bowel disease | KEGG:05321 | 0.00 |
| Th1 and Th2 cell differentiation | KEGG:04658 | 0.01 |
| Prostate cancer | KEGG:05215 | 0.01 |
| Neurotrophin signaling pathway | KEGG:04722 | 0.02 |
| Factor: Tcfap2b; motif: NYGCCYNARGGCAN; match class: 1 | TF:M02820_1 | 0.01 |
| Factor: TCFAP2B; motif: NTWGCCKSMGGCRNN; match class: 1 | TF:M02924_1 | 0.01 |
| Factor: FPM315; motif: SRGGGAGGAGGN | TF:M01587 | 0.02 |
| Factor: AP-2alphaA; motif: ANNGCCTNAGGSNNT | TF:M01045 | 0.02 |
| Factor: WT1; motif: NNGGGNGGGSGN; match class: 1 | TF:M07436_1 | 0.04 |
| Factor: RNF96; motif: BCCCGCRGCC; match class: 1 | TF:M01199_1 | 0.05 |
| Apoptosis | WP:WP1254 | 0.00 |
| Oxidative damage response | WP:WP1496 | 0.04 |

**Table 7** Wilcoxon Activated

| Term name | GO Term ID | P-value |
|---|---|---|
| immune response | GO:0006955 | 0.00 |
| inflammatory response | GO:0006954 | 0.00 |
| positive regulation of natural killer cell chemotaxis | GO:2000503 | 0.01 |
| regulation of T-helper 1 cell cytokine production | GO:2000554 | 0.02 |
| extracellular space | GO:0005615 | 0.00 |
| cytokine activity | GO:0005125 | 0.00 |
| tumor necrosis factor receptor binding | GO:0005164 | 0.05 |

**Table 8** Wilcoxon Naive

| Term name | GO Term ID | P-value |
|---|---|---|
| cellular response to peptide | GO:1901653 | 0.02 |

**Table 9** Wilcoxon Exhausted (D3)

| Term name | GO Term ID | P-value |
|---|---|---|
| cell killing | GO:0001906 | 0.00 |
| granzyme-mediated programmed cell death signaling pathway | GO:0140507 | 0.01 |
| cytolytic granule | GO:0044194 | 0.01 |

**Table 10** Wilcoxon Exhausted (D4)

| Term name | GO Term ID | P-value |
|---|---|---|
| inflammatory response | GO:0006954 | 0.01 |
| regulation of leukocyte mediated immunity | GO:0002703 | 0.02 |
| interleukin-18-mediated signaling pathway | GO:0035655 | 0.03 |
| cellular response to interleukin-18 | GO:0071351 | 0.03 |
| response to interleukin-18 | GO:0070673 | 0.05 |
| interleukin-18 receptor complex | GO:0045092 | 0.00 |
| external side of plasma membrane | GO:0009897 | 0.01 |
| receptor complex | GO:0043235 | 0.04 |
| interleukin-18 receptor activity | GO:0042008 | 0.00 |
| NAD+ nucleosidase activity, cyclic ADP-ribose generating | GO:0061809 | 0.02 |
| Viral protein interaction with cytokine and cytokine receptor | KEGG:04061 | 0.00 |
| Interleukin-18 signaling | REAC:R-MMU-9012546 | 0.00 |

### 5 Top 20 PR genes obtained using Netmap

The following three sections contain the input and output of GeneAgent. The ouput has been copied from the webtool. The inputs including the rankings can be found under input. The generation of the input tables is described in detail in the methods section of the main text. In brief, we use the top 20 genes (nodes) by node importance from the Netmap method; and aggregate the Wilcoxon ranks for all conditions to obtain a global rank for the Wilxocon based tests. Claude has been used to structure the raw output into sections without further content modification.

#### Input

Table 11 shows the top 20 genes from the PageRank call used as an input for GeneAgent.

| Gene Name | importance | Exhausted (D3) | Exhausted (D4) | Activated | Naive |
|---|---|---|---|---|---|
| Kcnq5 | 2.951318 | 0.977908 | 0.975141 | 0.998269 | NaN |
| Aff3 | 3.796726 | 0.961572 | 0.969371 | 0.878823 | 0.986960 |
| Bcl2 | 3.320430 | 0.933609 | 0.930266 | 0.694634 | 0.761923 |
| Tox | 3.161977 | 0.884774 | 0.983902 | 0.870052 | 0.423249 |
| Il12rb2 | 1.213374 | 0.841695 | 0.371679 | NaN | NaN |
| Nfat5 | 1.253901 | 0.795535 | 0.458366 | NaN | NaN |
| Ikzf2 | 1.718438 | 0.751061 | 0.967377 | NaN | NaN |
| Pitpnc1 | 1.906619 | 0.746585 | 0.474749 | 0.685286 | NaN |
| Apbb1ip | 2.745637 | 0.675368 | 0.811169 | 0.420427 | 0.838674 |
| Lef1 | 2.684630 | 0.672635 | 0.699836 | 0.891518 | 0.420641 |
| Ripor2 | 1.440048 | 0.665136 | 0.774913 | NaN | NaN |
| Prkcq | 0.605604 | 0.605604 | NaN | NaN | NaN |
| Kidins220 | 1.297280 | 0.603570 | 0.693710 | NaN | NaN |
| Ppp3ca | 1.949796 | 0.577873 | 0.875276 | NaN | 0.496647 |
| Irf2 | 1.715799 | 0.488576 | 0.774913 | NaN | 0.452310 |
| Gm10076 | 0.388582 | 0.388582 | NaN | NaN | NaN |
| Arhgap15 | 1.139815 | 0.364281 | NaN | 0.775534 | NaN |
| Alcam | 1.221809 | 0.309749 | NaN | 0.912060 | NaN |
| Enox2 | 0.290565 | 0.290565 | NaN | NaN | NaN |
| Znrf2 | 0.286204 | 0.286204 | NaN | NaN | NaN |

**Table 11** Input to Gene Agent: the top 20 PageRank genes

### Output

Analysis of your gene set is completed. You can copy the results or copy the retrieval URL and keep it to retrieve the results later. Analysis of gene set: Kcnq5, Aff3, Bcl2, Tox, Il12rb2, Nfat5, Ikzf2, Pitpnc1, Apbb1ip, Lef1 ...

#### Step 1: Initial generation for gene set

##### Process: T-cell Activation and Differentiation Regulation

The system of interacting proteins encoded by the provided gene set appears to play a central role in regulating T-cell activation, differentiation, and associated signaling pathways. This conclusion is based on the functional annotations and known interactions of the encoded proteins.

Kcnq5 encodes a potassium channel subunit that regulates membrane potential, which is critical for T-cell activation by modulating calcium signaling. Proper ion flux is essential for downstream signaling cascades in T-cells.

Aff3 encodes a transcriptional coactivator that influences gene expression during immune responses. It likely contributes to the transcriptional regulation of genes involved in T-cell differentiation.

Bcl2 encodes an anti-apoptotic protein that promotes T-cell survival during activation and differentiation. This is crucial for maintaining a functional T-cell population during immune responses.

Tox encodes a transcription factor that is essential for the development and maintenance of CD4+ T-cell subsets, particularly regulatory T-cells and exhausted T-cells during chronic infections or cancer.

Il12rb2 encodes a subunit of the IL-12 receptor, which is critical for Th1 differentiation. IL-12 signaling promotes the production of interferon-gamma (IFN-?), a key cytokine in cell-mediated immunity.

Nfat5 encodes a transcription factor that regulates gene expression in response to osmotic stress and is involved in T-cell activation under hypertonic conditions, which can occur in inflamed tissues.

Ikzf2 encodes Helios, a transcription factor that plays a role in the development and function of regulatory T-cells, contributing to immune tolerance and preventing autoimmunity.

Pitpnc1 encodes a phosphatidylinositol transfer protein that may regulate phosphoinositide signaling, which is critical for T-cell receptor (TCR) signaling and vesicular trafficking.

Apbb1ip encodes a protein involved in integrin activation and cytoskeletal rearrangements, which are necessary for T-cell migration and immune synapse formation.

Lef1 encodes a transcription factor in the Wnt signaling pathway, which is important for T-cell development in the thymus and differentiation of effector T-cells.

Ripor2 encodes a cytoskeletal regulator that modulates T-cell migration and polarity, essential for effective immune responses.

Prkcq encodes protein kinase C-theta, a key signaling molecule in TCR-mediated activation of NF-?B and AP-1, which are critical for T-cell activation and cytokine production.

Kidins220 encodes a scaffold protein involved in neurotrophin and TCR signaling, suggesting a role in T-cell activation and survival.

Ppp3ca encodes the catalytic subunit of calcineurin, a phosphatase that activates NFAT transcription factors during TCR signaling, driving T-cell activation and cytokine production.

Irf2 encodes a transcription factor that regulates the expression of interferon-stimulated genes, contributing to antiviral responses and T-cell function.

Gm10076 is a less characterized gene, but its potential role in this system may involve regulatory functions in immune signaling, requiring further investigation.

Arhgap15 encodes a Rho GTPase-activating protein that regulates actin cytoskeleton dynamics, which are critical for T-cell migration and immune synapse formation.

Alcam encodes an adhesion molecule that facilitates T-cell interactions with antigen-presenting cells, enhancing TCR signaling and immune synapse stability.

Enox2 encodes an oxidoreductase that may influence redox signaling, which is known to modulate T-cell activation and function.

Znrf2 encodes a RING finger protein with potential roles in protein ubiquitination, which could regulate TCR signaling or protein turnover in T-cells.

In summary, this system of interacting proteins collectively regulates T-cell activation, differentiation, survival, migration, and effector functions. The interplay between ion channels, transcription factors, signaling molecules, and cytoskeletal regulators ensures a coordinated immune response.

#### Step 2: Self-verification of the process name

**Original_claim:** The entire gene set is involved in the regulation of T-cell activation and differentiation. Here is the entire gene set used for verification:

##Kcnq5,Aff3,Bcl2,Tox,Il12rb2,Nfat5,Ikzf2,Pitpnc1,Apbb1ip,Lef1,Ripor2,Prkcq,Kidins220,Ppp3ca,Irf2,[illegible]

**Verified_claim:** The claim that the entire gene set is involved in the regulation of T-cell activation and differentiation is **partially supported** by the evidence.

1. The enrichment analysis of the gene set indicates significant associations with biological processes such as "T cell activation" (GO:0042110) and "Th1 and Th2 cell differentiation" (KEGG:04658). These processes are directly related to T-cell activation and differentiation, providing strong evidence for the claim.
2. The pathway analysis further supports the claim by identifying pathways such as "Calcineurin-dependent NFAT signaling role in lymphocytes," "Th1 and Th2 cell differentiation," and "IL-2/STAT5 Signaling." These pathways are known to play roles in T-cell activation and differentiation.
3. However, the claim states that the "entire gene set" is involved in these processes. The evidence shows that only subsets of the gene set are directly associated with the relevant biological processes and pathways. For example:
   - "T cell activation" involves 7 genes from the set.
   - "Th1 and Th2 cell differentiation" involves 3 genes from the set.
   - Other pathways also involve specific subsets of the gene set.

Conclusion: While there is strong evidence that parts of the gene set are involved in T-cell activation and differentiation, the claim that the "entire gene set" is involved is not fully supported. The evidence provides strong support for the involvement of some genes in the set but does not confirm the involvement of all genes in the specified processes.

#### Step 3: Modification

**Process: T cell activation**

The system of interacting proteins encoded by the provided gene set appears to play a central role in T cell activation, as supported by the verified enrichment analysis. This conclusion is based on the functional annotations and known interactions of the encoded proteins.

Kcnq5 encodes a potassium channel subunit that regulates membrane potential, which is critical for T cell activation by modulating calcium signaling. Proper ion flux is essential for downstream signaling cascades in T cells.

Aff3 encodes a transcriptional coactivator that influences gene expression during immune responses. It likely contributes to the transcriptional regulation of genes involved in T cell activation.

Bcl2 encodes an anti-apoptotic protein that promotes T cell survival during activation. This is crucial for maintaining a functional T cell population during immune responses.

Tox encodes a transcription factor that is essential for the development and maintenance of CD4+ T cell subsets, particularly regulatory T cells and exhausted T cells during chronic infections or cancer.

Il12rb2 encodes a subunit of the IL-12 receptor, which is critical for Th1 differentiation. IL-12 signaling promotes the production of interferon-gamma (IFN-?), a key cytokine in cell-mediated immunity.

Nfat5 encodes a transcription factor that regulates gene expression in response to osmotic stress and is involved in T cell activation under hypertonic conditions, which can occur in inflamed tissues.

Ikzf2 encodes Helios, a transcription factor that plays a role in the development and function of regulatory T cells, contributing to immune tolerance and preventing autoimmunity.

Pitpnc1 encodes a phosphatidylinositol transfer protein that may regulate phosphoinositide signaling, which is critical for T cell receptor (TCR) signaling and vesicular trafficking.

Apbb1ip encodes a protein involved in integrin activation and cytoskeletal rearrangements, which are necessary for T cell migration and immune synapse formation.

Lef1 encodes a transcription factor in the Wnt signaling pathway, which is important for T cell development in the thymus and differentiation of effector T cells.

Ripor2 encodes a cytoskeletal regulator that modulates T cell migration and polarity, essential for effective immune responses.

Prkcq encodes protein kinase C-theta, a key signaling molecule in TCR-mediated activation of NF-?B and AP-1, which are critical for T cell activation and cytokine production.

Kidins220 encodes a scaffold protein involved in neurotrophin and TCR signaling, suggesting a role in T cell activation and survival.

Ppp3ca encodes the catalytic subunit of calcineurin, a phosphatase that activates NFAT transcription factors during TCR signaling, driving T cell activation and cytokine production.

Irf2 encodes a transcription factor that regulates the expression of interferon-stimulated genes, contributing to antiviral responses and T cell function.

Gm10076 is a less characterized gene, but its potential role in this system may involve regulatory functions in immune signaling.

Arhgap15 encodes a Rho GTPase-activating protein that regulates actin cytoskeleton dynamics, which are critical for T cell migration and immune synapse formation.

Alcam encodes an adhesion molecule that facilitates T cell interactions with antigen-presenting cells, enhancing TCR signaling and immune synapse stability.

Enox2 encodes an oxidoreductase that may influence redox signaling, which is known to modulate T cell activation and function.

Znrf2 encodes a RING finger protein with potential roles in protein ubiquitination, which could regulate TCR signaling or protein turnover in T cells.

In summary, this system of interacting proteins collectively regulates T cell activation, survival, migration, and effector functions. The interplay between ion channels, transcription factors, signaling molecules, and cytoskeletal regulators ensures a coordinated immune response.

#### Step 4: Self-verification of the analytical narratives

**Original_claim:** Kcnq5 regulates membrane potential and modulates calcium signaling, which is critical for T cell activation by ensuring proper ion flux for downstream signaling cascades.

**Verified_claim:** The claim that "Kcnq5 regulates membrane potential and modulates calcium signaling, which is critical for T cell activation by ensuring proper ion flux for downstream signaling cascades" is weakly supported based on the following evidence:

1. **Gene Summary**: Kcnq5 (potassium voltage-gated channel, subfamily Q, member 5) enables voltage-gated potassium channel activity and acts upstream of or within potassium ion transport. It is predicted to be located in the plasma membrane and is part of the voltage-gated potassium channel complex. This aligns with the regulation of membrane potential but does not directly confirm its role in calcium signaling or T cell activation.
2. **Enrichment Analysis**: Kcnq5 is associated with the "KCNQ3-KCNQ5 complex," which is significant in the context of potassium channel activity. However, this does not directly link it to calcium signaling or T cell activation.
3. **Domains and Diseases**: No specific biological domains or diseases were identified for Kcnq5 that directly connect it to calcium signaling or T cell activation.

While the evidence supports the role of Kcnq5 in regulating membrane potential through potassium ion transport, there is no direct evidence provided here to confirm its involvement in calcium signaling or T cell activation. Therefore, the claim is weakly supported.

#### Step 5: Summarization

##### Process: T cell activation

The system of interacting proteins encoded by the provided gene set plays a role in T cell activation, as supported by the verified enrichment analysis. Below is the revised summary based on the verification report.

Kcnq5 regulates membrane potential through potassium ion transport, which is essential for maintaining the electrical properties of T cells during activation. However, its direct involvement in calcium signaling or T cell activation remains unclear.

Bcl2 encodes an anti-apoptotic protein that promotes T cell survival by preventing apoptotic cell death, ensuring the maintenance of a functional T cell population during immune responses.

Il12rb2 encodes a subunit of the IL-12 receptor, which is critical for Th1 differentiation and promotes interferon-gamma (IFN-?) production, a key cytokine in cell-mediated immunity.

Ppp3ca encodes the catalytic subunit of calcineurin, a phosphatase that activates NFAT transcription factors during TCR signaling, driving T cell activation and cytokine production.

Ripor2 modulates T cell migration and polarity by inhibiting RhoA activity, which is essential for effective immune responses.

Lef1 is a transcription factor in the Wnt signaling pathway, important for T cell development in the thymus and differentiation of effector T cells.

Apbb1ip facilitates integrin activation and cytoskeletal rearrangements, which are necessary for T cell migration and immune synapse formation. Arhgap15 regulates Rho GTPases, contributing to actin cytoskeleton dynamics and supporting T cell migration.

Alcam encodes an adhesion molecule that binds to CD6, facilitating T cell interactions with antigen-presenting cells and supporting immune synapse formation.

Prkcq encodes protein kinase C-theta, which is involved in TCR signaling and promotes interleukin-2 production, contributing to T cell activation.

Kidins220 acts as a scaffold protein mediating neurotrophin signaling, but its role in TCR signaling and T cell activation remains ambiguous.

Nfat5 regulates gene expression in response to osmotic stress, which may indirectly influence T cell activation under hypertonic conditions.

Tox is involved in T cell development, but its specific role in regulatory or exhausted T cells during chronic infections or cancer is not fully established.

Aff3 functions as a transcriptional coactivator expressed in lymphoid tissues, but its direct role in T cell activation is not confirmed.

Irf2 regulates interferon-stimulated genes, contributing to antiviral responses, but its role in T cell function is not explicitly verified.

Pitpnc1 facilitates phosphoinositide signaling, which is important for lipid metabolism, but its direct involvement in TCR signaling or vesicular trafficking during T cell activation is unclear.

Enox2 influences redox signaling through NADH oxidation and protein disulfide interchange, but its role in modulating T cell activation is not established.

Znrf2 encodes a RING finger protein involved in protein ubiquitination, but its specific role in TCR signaling or protein turnover in T cells is not confirmed.

In summary, this system of interacting proteins collectively contributes to T cell activation, survival, migration, and effector functions. The interplay between signaling molecules, transcription factors, adhesion proteins, and cytoskeletal regulators ensures a coordinated immune response.

### 6 Comparison: Wilcoxon with groups Naive, Active, Exhausted

#### Input

**Table 12** Gene Aggregated Distance and Rank

| Index | Gene | Aggregated Distance | Aggregated Rank |
|---|---|---|---|
| 0 | Mki67 | 5 | 1 |
| 1 | Rad51b | 11 | 2 |
| 2 | Kcnq5 | 16 | 3 |
| 3 | Cenpp | 22 | 4 |
| 4 | Itpr1 | 26 | 5 |
| 5 | Mir155hg | 32 | 6 |
| 6 | Hist1h1b | 33 | 7 |
| 7 | Lgals1 | 37 | 8 |
| 8 | Nfkb1 | 39 | 9 |
| 9 | Gzmb | 39 | 10 |
| 10 | Themis | 40 | 11 |
| 11 | Hivep3 | 45 | 12 |
| 12 | Cenpe | 47 | 13 |
| 13 | Adamts6 | 48 | 14 |
| 14 | Ctla4 | 48 | 15 |
| 15 | Taf1d | 53 | 16 |
| 16 | Knl1 | 53 | 17 |
| 17 | Il12rb2 | 55 | 18 |
| 18 | Nfat5 | 57 | 19 |
| 19 | Btg1 | 58 | 20 |

### Output

Analysis of your gene set is completed. You can copy the results or copy the retrieval URL and keep it to retrieve the results later. Analysis of gene set: Mki67, Rad51b, Kcnq5, Cenpp, Itpr1, Mir155hg, Hist1h1b, Lgals1, Nfkb1, Gzmb ...

#### Step 1: Initial generation for gene set

**Process: Chromosome Dynamics and Immune Regulation**

The system of interacting proteins encoded by the provided gene set appears to perform a dual role, primarily in chromosome dynamics during cell division and in modulating immune responses. Below is a critical analysis of the biological processes performed by this system:

1. **Chromosome Segregation and Mitotic Regulation:** The genes Mki67, Cenpp, Cenpe, Knl1, and Rad51b are central to chromosome dynamics. Mki67 encodes a proliferation marker associated with chromatin organization during mitosis. Cenpp and Cenpe are involved in centromere and kinetochore function, essential for accurate chromosome segregation. Knl1 is a scaffold protein critical for kinetochore assembly and spindle checkpoint signaling. Rad51b participates in homologous recombination repair, ensuring genomic stability during DNA replication. Together, these proteins coordinate chromosome alignment, segregation, and repair, which are vital for cell division and genomic integrity.
2. **Immune Response Modulation:** Several genes in this set are key regulators of immune processes. Nfkb1 encodes a subunit of the NF-?B transcription factor, which controls the expression of cytokines and immune response genes. Il12rb2

encodes a receptor subunit for interleukin-12, promoting T-helper 1 (Th1) cell differentiation. Ctla4 is a critical immune checkpoint protein that downregulates T-cell activation. Mir155hg encodes a precursor for miR-155, a microRNA involved in immune cell differentiation and inflammatory responses. These genes collectively regulate immune activation, cytokine signaling, and immune tolerance.

3. **Calcium Signaling and Apoptosis:** Itpr1 encodes an inositol 1,4,5-trisphosphate receptor, which mediates calcium release from the endoplasmic reticulum. Calcium signaling is crucial for various cellular processes, including T-cell activation and apoptosis. Gzmb encodes granzyme B, a serine protease involved in inducing apoptosis in target cells during immune responses. These proteins link calcium signaling to immune cell function and programmed cell death.
4. **Chromatin Structure and Transcriptional Regulation:** Hist1h1b encodes a linker histone H1 variant, which modulates chromatin compaction and transcriptional regulation. Taf1d is a component of the TFIID complex, essential for RNA polymerase II-mediated transcription initiation. Hivep3 encodes a transcriptional regulator involved in immune and developmental gene expression. These proteins influence gene expression by modulating chromatin accessibility and transcriptional machinery.
5. **Cellular Differentiation and Proliferation:** Btg1 is a regulator of cell cycle progression and differentiation, particularly in hematopoietic and neural cells. Adamts6 encodes a metalloproteinase involved in extracellular matrix remodeling, which can influence cell migration and tissue development. Nfat5 is a transcription factor that regulates osmotic stress responses and immune cell differentiation. These proteins contribute to cellular growth, differentiation, and adaptation to environmental changes.
6. **Glycan Binding and Cell-Cell Interactions:** Lgals1 encodes galectin-1, a lectin that binds ?-galactoside sugars and modulates cell-cell and cell-matrix interactions. It plays roles in immune tolerance, apoptosis, and tumor progression. This protein highlights the system's involvement in extracellular signaling and immune regulation.
7. **T-Cell Development and Function:** Themis is a critical regulator of T-cell development and signaling, ensuring proper selection and activation of T-cells in the thymus. This function is essential for maintaining immune homeostasis and preventing autoimmunity.

In summary, this system of interacting proteins is primarily involved in chromosome dynamics, immune regulation, and associated processes such as transcriptional control, calcium signaling, and apoptosis. The interplay between these functions highlights the system's role in maintaining cellular and immune homeostasis.

#### Step 2: Self-verification of the process name

**Original_claim:** The entire gene set is involved in chromosome dynamics and immune regulation. Here is the entire gene set used for verification:

##Mki67,Rad51b,Kcnq5,Cenpp,Itpr1,Mir155hg,Hist1h1b,Lgals1,Nfkb1,Gzmb,Themis,Hivep3,Cenpe,Adamts6

**Verified_claim:** The claim that the gene set is involved in chromosome dynamics and immune regulation is strongly supported.

Evidence:

1. Enrichment analysis reveals significant associations with terms related to immune regulation, such as “T cell activation SARS CoV 2” and “Integrative Analysis of Treg Glial Interactions.” These terms indicate involvement in immune-related processes.
2. Enrichment analysis also identifies “condensed chromosome” as a significant term, which is directly related to chromosome dynamics.
3. Pathway analysis further supports the claim:
    - Immune regulation is evidenced by pathways like “Interleukin-12-mediated signaling events” (genes: GZMB, NFKB1, IL12RB2) and “T cell signal transduction” (genes: NFAT5, CTLA4, NFKB1).
    - Chromosome dynamics are supported by pathways such as “Unattached Kinetochores Signal Amplification Via A MAD2 Inhibitory Signal” and “Resolution Of Sister Chromatid Cohesion,” involving genes CENPE, KNL1, and CENPP.

These findings collectively confirm the involvement of the gene set in both chromosome dynamics and immune regulation.

**Original_claim:** The gene set plays a role in the regulation of immune responses and chromosomal processes. Here is the entire gene set used for verification:

##Mki67,Rad51b,Kcnq5,Cenpp,Itpr1,Mir155hg,Hist1h1b,Lgals1,Nfkb1,Gzmb,Themis,Hivep3,Cenpe,Ada[illegible]

**Verified_claim:** The claim that the gene set plays a role in the regulation of immune responses and chromosomal processes is strongly supported by the evidence.

1. **Enrichment Analysis:**
    - The gene set is significantly associated with “T cell activation SARS CoV 2” and “Integrative Analysis of Treg Glial Interactions,” which are directly related to immune responses.
    - The gene set is also associated with “condensed chromosome,” indicating involvement in chromosomal processes.
2. **Pathway Analysis:**
    - Immune-related pathways include “Interleukin-12-mediated signaling events” and “T cell signal transduction,” with overlapping genes such as GZMB, NFKB1, IL12RB2, NFAT5, and CTLA4.
    - Chromosomal-related pathways include “Unattached Kinetochores Signal Amplification Via A MAD2 Inhibitory Signal,” “EML4 And NUDC In Mitotic Spindle Formation,” and “Resolution Of Sister Chromatid Cohesion,” with overlapping genes such as CENPE, KNL1, and CENPP.

These findings provide strong evidence that the gene set is involved in both immune responses and chromosomal processes.

**Original_claim:** The entire gene set contributes to the coordination of chromosome behavior and immune system functions. Here is the entire gene set used for verification:

##Mki67,Rad51b,Kcnq5,Cenpp,Itpr1,Mir155hg,Hist1h1b,Lgals1,Nfkb1,Gzmb,Themis,Hivep3,Cenpe,Adamts6

**Verified_claim:** The claim that the entire gene set contributes to the coordination of chromosome behavior and immune system functions is **partially supported** by the evidence.

1. **Immune System Functions**:
   - Enrichment analysis identified pathways such as "T cell activation SARS CoV 2" and "Integrative Analysis of Treg Glial Interactions," which are directly related to immune system functions.
   - Pathway analysis revealed terms like "Interleukin-12-mediated signaling events" (involving genes GZMB, NFKB1, IL12RB2) and "T cell signal transduction" (involving genes NFAT5, CTLA4, NFKB1), both of which are critical to immune system processes.
2. **Chromosome Behavior**:
   - Enrichment analysis identified the term "condensed chromosome," which is associated with chromosome structure and behavior.
   - Pathway analysis highlighted terms such as "Unattached Kinetochores Signal Amplification Via A MAD2 Inhibitory Signal," "EML4 And NUDC In Mitotic Spindle Formation," and "Resolution Of Sister Chromatid Cohesion," all of which involve genes (CENPE, KNL1, CENPP) that play roles in chromosome segregation and mitotic processes.

While the evidence strongly supports the involvement of the gene set in both immune system functions and chromosome behavior, it does not confirm that the "entire" gene set contributes to these processes. Some genes may not be directly involved in these specific functions. Therefore, the claim is partially supported.

**Original_claim:** Here is the entire gene set used for verification:

##Mki67,Rad51b,Kcnq5,Cenpp,Itpr1,Mir155hg,Hist1h1b,Lgals1,Nfkb1,Gzmb,Themis,Hivep3,Cenpe,Adamts6

**Verified_claim:** The gene set provided has significant enrichment in several biological functions and pathways. Here are the findings:

1. **T cell activation SARS CoV 2**: This pathway is significantly enriched, indicating a potential association with immune response mechanisms, particularly in the context of SARS-CoV-2.
2. **Integrative Analysis of Treg Glial Interactions**: This enrichment suggests involvement in regulatory T cell and glial cell interactions, which are important in immune regulation and neurobiology.
3. **Condensed Chromosome**: The gene set shows enrichment in functions related to chromosomal structure, which may be relevant to cell division and genetic material organization.
4. **hsa-mir-6512-5p**: Enrichment in this microRNA pathway suggests potential regulatory roles in gene expression.
5. **Apoptosis**: The gene set is significantly associated with apoptosis, a critical process in programmed cell death and cellular homeostasis.

These enrichments provide evidence of the gene set's involvement in immune response, cellular regulation, and apoptosis. If the claim relates to these functions or pathways, the findings offer strong support. If the claim is unrelated, the evidence may be considered weak or irrelevant.

#### Step 3: Modification

##### Process: Chromosome Dynamics and Immune Regulation

The system of interacting proteins encoded by the provided gene set appears to perform a dual role, primarily in chromosome dynamics during cell division and in modulating immune responses. Below is a critical analysis of the biological processes performed by this system:

**Chromosome Segregation and Mitotic Regulation:** The genes Mki67, Cenpp, Cenpe, Knl1, and Rad51b are central to chromosome dynamics. Mki67 encodes a proliferation marker associated with chromatin organization during mitosis. Cenpp and Cenpe are involved in centromere and kinetochore function, essential for accurate chromosome segregation. Knl1 is a scaffold protein critical for kinetochore assembly and spindle checkpoint signaling. Rad51b participates in homologous recombination repair, ensuring genomic stability during DNA replication. Together, these proteins coordinate chromosome alignment, segregation, and repair, which are vital for cell division and genomic integrity.

**Immune Response Modulation:** Several genes in this set are key regulators of immune processes. Nfkb1 encodes a subunit of the NF-?B transcription factor, which controls the expression of cytokines and immune response genes. Il12rb2 encodes a receptor subunit for interleukin-12, promoting T-helper 1 (Th1) cell differentiation. Ctla4 is a critical immune checkpoint protein that downregulates T-cell activation. Mir155hg encodes a precursor for miR-155, a microRNA involved in immune cell differentiation and inflammatory responses. These genes collectively regulate immune activation, cytokine signaling, and immune tolerance.

**Calcium Signaling and Apoptosis:** Itpr1 encodes an inositol 1,4,5-trisphosphate receptor, which mediates calcium release from the endoplasmic reticulum. Calcium signaling is crucial for various cellular processes, including T-cell activation and apoptosis. Gzmb encodes granzyme B, a serine protease involved in inducing apoptosis in target cells during immune responses. These proteins link calcium signaling to immune cell function and programmed cell death.

**Chromatin Structure and Transcriptional Regulation:** Hist1h1b encodes a linker histone H1 variant, which modulates chromatin compaction and transcriptional regulation. Taf1d is a component of the TFIID complex, essential for RNA polymerase II-mediated transcription initiation. Hivep3 encodes a transcriptional regulator involved in immune and developmental gene expression. These proteins influence gene expression by modulating chromatin accessibility and transcriptional machinery.

**Cellular Differentiation and Proliferation:** Btg1 is a regulator of cell cycle progression and differentiation, particularly in hematopoietic and neural cells. Adamts6 encodes a metalloproteinase involved in extracellular matrix remodeling, which can influence cell migration and tissue development. Nfat5 is a transcription factor that regulates osmotic stress responses and immune cell differentiation. These proteins contribute to cellular growth, differentiation, and adaptation to environmental changes.

**Glycan Binding and Cell-Cell Interactions:** Lgals1 encodes galectin-1, a lectin that binds ?-galactoside sugars and modulates cell-cell and cell-matrix interactions. It plays roles in immune tolerance, apoptosis, and tumor progression. This protein highlights the system's involvement in extracellular signaling and immune regulation.

**T-Cell Development and Function:** Themis is a critical regulator of T-cell development and signaling, ensuring proper selection and activation of T-cells in the thymus. This function is essential for maintaining immune homeostasis and preventing autoimmunity.

In summary, this system of interacting proteins is primarily involved in chromosome dynamics, immune regulation, and associated processes such as transcriptional control, calcium signaling, and apoptosis. The interplay between these functions highlights the system's role in maintaining cellular and immune homeostasis.

#### Step 4: Self-verification of the analytical narratives

**Original_claim:** Mki67, Cenpp, Cenpe, Knl1, and Rad51b are involved in chromosome dynamics, including chromatin organization, centromere and kinetochore function, spindle checkpoint signaling, and homologous recombination repair, which are essential for chromosome segregation and genomic stability during cell division.

**Verified_claim:** The claim that Mki67, Cenpp, Cenpe, Knl1, and Rad51b are involved in chromosome dynamics, including chromatin organization, centromere and kinetochore function, spindle checkpoint signaling, and homologous recombination repair, which are essential for chromosome segregation and genomic stability during cell division, is strongly supported by the following evidence:

1. **Enrichment Analysis:**
   - The gene set is significantly associated with terms such as "condensed chromosome" (GO:0000793), "chromosome segregation" (GO:0007059), and "Mitotic Spindle Checkpoint" (REAC:R-HSA-69618). These terms directly relate to chromosome dynamics, spindle checkpoint signaling, and chromosome segregation.
2. **Pathway Analysis:**
   - The genes are involved in pathways such as "Unattached Kinetochores Signal Amplification Via A MAD2 Inhibitory Signal" (R-HSA-141444), "Resolution Of Sister Chromatid Cohesion" (R-HSA-2500257), and "G2-M Checkpoint." These pathways are critical for centromere and kinetochore function, spindle checkpoint signaling, and genomic stability during cell division.
3. **Specific Gene Contributions:**
   - Genes like CENPE, KNL1, and CENPP are repeatedly highlighted in pathways and enrichment terms related to kinetochore function and spindle checkpoint signaling.

- MKI67 is associated with the G2-M checkpoint, which is essential for genomic stability during cell division.

The evidence strongly supports the claim, as the genes are directly linked to the described biological processes and functions.

**Original_claim:** Nfkb1, Il12rb2, Ctla4, and Mir155hg regulate immune responses by controlling cytokine expression, T-helper 1 (Th1) cell differentiation, T-cell activation, and inflammatory processes.

**Verified_claim:** The claim that Nfkb1, Il12rb2, Ctla4, and Mir155hg regulate immune responses by controlling cytokine expression, T-helper 1 (Th1) cell differentiation, T-cell activation, and inflammatory processes is **strongly supported** by the evidence.

1. **Nfkb1**:
   - The gene encodes a protein that is a DNA-binding subunit of the NF-kappa-B (NFKB) protein complex, which is a transcription regulator activated by various stimuli, including cytokines. It stimulates the expression of genes involved in inflammatory responses and immune regulation. Inappropriate activation of NFKB is associated with inflammatory diseases, and its inhibition can lead to delayed immune cell development. This supports its role in regulating cytokine expression and inflammatory processes.
2. **Il12rb2**:
   - This gene encodes a subunit of the interleukin-12 receptor complex. It is upregulated by interferon-gamma in Th1 cells and plays a role in Th1 cell differentiation. Its upregulation is associated with inflammatory responses in diseases like Crohn's disease and leprosy, supporting its involvement in Th1 differentiation and inflammatory processes.
3. **Ctla4**:
   - This gene encodes a protein that transmits inhibitory signals to T cells, playing a critical role in T-cell activation and immune response regulation. Mutations in this gene are associated with autoimmune diseases, further supporting its role in immune regulation.
4. **Mir155hg**:
   - This gene represents a microRNA host gene expressed at high levels in lymphoma and may function as an oncogene. While its direct role in immune regulation is less explicitly stated, its association with immune-related processes is implied.
5. **Enrichment Analysis**:
   - The enrichment analysis for the gene set (Nfkb1, Il12rb2, Ctla4, Mir155hg) shows significant associations with pathways such as "T cell activation SARS CoV 2," "Integrative Analysis of Treg Glial Interactions," "T cell receptor and co-stimulatory signaling," and "Inflammatory bowel disease signaling."

These pathways are directly related to immune responses, T-cell activation, and inflammatory processes.

The evidence collectively supports the claim that these genes regulate immune responses through the mentioned mechanisms.

**Original_claim:** Itpr1 and Gzmb link calcium signaling to immune cell activation and apoptosis, playing critical roles in immune responses and programmed cell death.

**Verified_claim:** The claim that "Itpr1 and Gzmb link calcium signaling to immune cell activation and apoptosis, playing critical roles in immune responses and programmed cell death" is strongly supported by the following evidence:

1. **Gene Summary for ITPR1**:
   - ITPR1 encodes an intracellular receptor for inositol 1,4,5-trisphosphate, which mediates calcium release from the endoplasmic reticulum upon stimulation. This directly links ITPR1 to calcium signaling.
2. **Gene Summary for GZMB**:
   - GZMB encodes granzyme B, a serine protease secreted by natural killer (NK) cells and cytotoxic T lymphocytes (CTLs). It induces apoptosis in target cells, processes cytokines, and degrades extracellular matrix proteins, implicating it in immune cell activation and programmed cell death.
3. **Enrichment Analysis**:
   - The gene set containing ITPR1 and GZMB is significantly enriched in the biological process of "Apoptosis" (KEGG pathway), confirming their roles in programmed cell death.
   - ITPR1 is associated with "inositol 1,4,5-trisphosphate receptor activity involved in regulation of postsynaptic cytosolic calcium levels," further supporting its role in calcium signaling.

These findings collectively validate the claim, as both genes are functionally linked to calcium signaling, immune cell activation, and apoptosis.

**Original_claim:** Hist1h1b, Taf1d, and Hivep3 modulate chromatin structure and transcriptional regulation, influencing gene expression and immune-related transcriptional processes.

**Verified_claim:** The claim that Hist1h1b, Taf1d, and Hivep3 modulate chromatin structure and transcriptional regulation, influencing gene expression and immune-related transcriptional processes, is **partially supported** based on the following evidence:

1. **Hist1h1b (H1-5)**:
   - Summary: Hist1h1b encodes a replication-dependent histone that is part of the histone H1 family. Histones are basic nuclear proteins responsible for the nucleosome structure of chromosomal fibers in eukaryotes. The linker histone H1 interacts with linker DNA between nucleosomes and functions in the compaction

of chromatin into higher-order structures. This supports its role in modulating chromatin structure and transcriptional regulation.

2. **Taf1d**:

    - Summary: Taf1d is a member of the SL1 complex, which includes TBP and other TAF1 proteins, and plays a role in RNA polymerase I transcription. This indicates its involvement in transcriptional regulation but does not directly link it to chromatin structure or immune-related transcriptional processes.

3. **Hivep3**:

    - Summary: Hivep3 encodes a transcription factor that regulates nuclear factor kappaB-mediated transcription by binding the kappaB motif in target genes. It also binds the recombination signal sequence flanking the V, D, and J regions of immunoglobulin and T-cell receptors. This supports its role in transcriptional regulation and immune-related transcriptional processes.

4. **Gene Set Enrichment**:

    - No specific enrichment data was retrieved for the gene set "Hist1h1b, Taf1d, Hivep3," which limits the ability to confirm broader biological functions or pathways.

In conclusion, while there is strong evidence for the involvement of Hist1h1b and Hivep3 in chromatin structure, transcriptional regulation, and immune-related processes, the evidence for Taf1d's role in these specific functions is less direct. The claim is therefore partially supported.

**Original_claim:** Btg1, Adamts6, and Nfat5 contribute to cellular differentiation, proliferation, and adaptation to environmental stress, particularly in hematopoietic and immune cells.

**Verified_claim:** The claim that Btg1, Adamts6, and Nfat5 contribute to cellular differentiation, proliferation, and adaptation to environmental stress, particularly in hematopoietic and immune cells, is partially supported by evidence.

1. **Btg1**:

    - Summary: Btg1 is a member of an anti-proliferative gene family that regulates cell growth and differentiation. It is expressed highly in the G0/G1 phases of the cell cycle and downregulated as cells progress through G1. It interacts with nuclear receptors and functions as a coactivator of cell differentiation. This gene is involved in chromosomal translocation in B-cell chronic lymphocytic leukemia.
    - Evidence: Btg1 contributes to cellular differentiation and proliferation, but there is no direct mention of adaptation to environmental stress or specific involvement in hematopoietic and immune cells.

2. **Adamts6**:

- Summary: Adamts6 encodes a member of the ADAMTS protein family, which includes distinct protein modules such as a metalloproteinase domain and thrombospondin type 1 motif. Expression of Adamts6 may be regulated by the cytokine TNF-alpha.
- Evidence: While Adamts6 is involved in processes regulated by TNF-alpha, which is relevant to immune responses, there is no direct evidence linking it to cellular differentiation, proliferation, or adaptation to environmental stress in hematopoietic and immune cells.

3. **Nfat5**:
   - Summary: Nfat5 is a transcription factor that plays a central role in inducible gene transcription during the immune response. It regulates gene expression induced by osmotic stress in mammalian cells and exists as a homodimer forming stable dimers with DNA elements.
   - Evidence: Nfat5 is directly involved in adaptation to environmental stress (osmotic stress) and plays a role in immune responses, supporting its relevance to hematopoietic and immune cells.

Decision: The claim is **partially supported**. While Nfat5 has clear evidence for its role in adaptation to environmental stress and immune cell function, the roles of Btg1 and Adamts6 in hematopoietic and immune cells or environmental stress adaptation are not explicitly confirmed.

**Original_claim:** Lgals1 mediates glycan binding and cell-cell interactions, playing roles in immune tolerance, apoptosis, and extracellular signaling.

**Verified_claim:** The claim that "Lgals1 mediates glycan binding and cell-cell interactions, playing roles in immune tolerance, apoptosis, and extracellular signaling" is partially supported by the evidence.

1. **Support for glycan binding and cell-cell interactions**:
   - According to the gene summary for LGALS1 (galectin 1), it is a member of the galectin family, which are beta-galactoside-binding proteins. These proteins are implicated in modulating cell-cell and cell-matrix interactions. This directly supports the claim that Lgals1 mediates glycan binding and cell-cell interactions.
2. **Support for roles in immune tolerance, apoptosis, and extracellular signaling**:
   - The gene summary does not explicitly mention immune tolerance, apoptosis, or extracellular signaling. However, the role of galectins in modulating cell-cell interactions and acting as autocrine factors suggests potential involvement in these processes. This association is indirect and not explicitly confirmed in the provided evidence.
3. **Disease association**:
   - No specific diseases were found to be associated with Lgals1 in the disease database query, which does not directly impact the claim but indicates a lack of disease-specific evidence.

Decision: The claim is **partially supported**. While there is strong evidence for Lgals1's role in glycan binding and cell-cell interactions, its involvement in immune tolerance, apoptosis, and extracellular signaling is not explicitly confirmed in the provided evidence and remains ambiguous.

**Original_claim:** Themis is essential for T-cell development and signaling, ensuring proper selection and activation of T-cells in the thymus to maintain immune homeostasis.

**Verified_claim:** The claim that Themis is essential for T-cell development and signaling, ensuring proper selection and activation of T-cells in the thymus to maintain immune homeostasis, is **strongly supported**.

Evidence:

1. According to the gene summary for THEMIS, it encodes a protein that plays a regulatory role in both positive and negative T-cell selection during late thymocyte development. This directly supports the claim that Themis is essential for T-cell development and signaling.
2. The summary also states that the protein functions through T-cell antigen receptor signaling and is necessary for proper lineage commitment and maturation of T-cells. This aligns with the claim that Themis ensures proper selection and activation of T-cells in the thymus.
3. The role of Themis in T-cell antigen receptor signaling and thymocyte development is critical for maintaining immune homeostasis, as proper T-cell selection and activation are fundamental to immune system function.

No contradictory evidence was found.

**Original_claim:**

**Verified_claim:** Failed.

#### Step 5: Summarization

##### Process: Chromosome Dynamics and Immune Regulation

The system of interacting proteins encoded by the provided gene set performs roles in chromosome dynamics during cell division and immune regulation. Below is a revised analysis based on the verification report:

**Chromosome Segregation and Mitotic Regulation:** Mki67, Cenpp, Cenpe, Knl1, and Rad51b are involved in chromosome dynamics. Mki67 is associated with chromatin organization during mitosis and the G2-M checkpoint, ensuring genomic stability. Cenpp, Cenpe, and Knl1 are critical for centromere and kinetochore function, spindle checkpoint signaling, and chromosome segregation. Rad51b participates in homologous recombination repair, maintaining genomic integrity during cell division. These genes are enriched in pathways such as "Resolution Of Sister Chromatid Cohesion" and "Unattached Kinetochores Signal Amplification Via A MAD2 Inhibitory Signal," confirming their roles in chromosome dynamics.

**Immune Response Modulation:** Nfkb1, Il12rb2, Ctla4, and Mir155hg regulate immune responses. Nfkb1 encodes a subunit of the NF-?B transcription factor, controlling cytokine expression and inflammatory processes. Il12rb2 promotes T-helper 1 (Th1) cell differentiation and inflammatory responses. Ctla4 transmits inhibitory signals to T cells, regulating immune activation. Mir155hg encodes a precursor for miR-155, which influences immune cell differentiation. These genes are enriched in pathways such as "Interleukin-12-mediated signaling events" and "T cell signal transduction," supporting their roles in immune regulation.

**Calcium Signaling and Apoptosis:** Itpr1 and Gzmb link calcium signaling to immune cell activation and apoptosis. Itpr1 mediates calcium release from the endoplasmic reticulum, essential for immune cell signaling. Gzmb encodes granzyme B, which induces apoptosis in target cells and processes cytokines. These genes are enriched in pathways related to apoptosis and calcium signaling, confirming their roles in immune responses and programmed cell death.

**Chromatin Structure and Transcriptional Regulation:** Hist1h1b and Hivep3 modulate chromatin structure and transcriptional regulation. Hist1h1b encodes a linker histone H1 variant, which compacts chromatin into higher-order structures. Hivep3 regulates NF-?B-mediated transcription and binds recombination signal sequences in immunoglobulin and T-cell receptor genes, influencing immune-related transcriptional processes. Taf1d is involved in RNA polymerase I transcription but lacks direct evidence for chromatin modulation or immune-related functions.

**Cellular Differentiation and Proliferation:** Nfat5 contributes to cellular adaptation to environmental stress, particularly osmotic stress, and plays a role in immune responses. Btg1 regulates cell cycle progression and differentiation but lacks direct evidence for involvement in hematopoietic or immune cells. Adamts6 is associated with TNF-alpha regulation but lacks direct evidence for cellular differentiation or environmental stress adaptation.

**Glycan Binding and Cell-Cell Interactions:** Lgals1 mediates glycan binding and cell-cell interactions, modulating immune tolerance and apoptosis. It binds ?-galactoside sugars and influences extracellular signaling, though its roles in immune tolerance and apoptosis are less explicitly confirmed.

**T-Cell Development and Function:** Themis is essential for T-cell development and signaling. It regulates positive and negative T-cell selection during thymocyte development and ensures proper lineage commitment and activation of T-cells, maintaining immune homeostasis.

In summary, this system of interacting proteins is primarily involved in chromosome dynamics, immune regulation, and associated processes such as transcriptional control, calcium signaling, and apoptosis. The interplay between these functions highlights the system's role in maintaining cellular and immune homeostasis.

### 7 Comparison: Wilcoxon with groups Naive, Activated, Exhausted (D3) and Exhausted (D4)

#### Input

#### Output

Analysis of your gene set is completed. You can copy the results or copy the retrieval URL and keep it to retrieve the results later. Analysis of gene set: Mki67, Cmss1, Gzmb, Skap1, Rad51b, Pitpnc1, Cenpp, Il2ra, Taf1d, Arhgap15 ...

**Table 13** Gene Aggregated Distance and Rank

| Index | Gene | Aggregated Distance | Aggregated Rank |
|---|---|---|---|
| 0 | Mki67 | 37 | 1 |
| 1 | Cmss1 | 50 | 2 |
| 2 | Gzmb | 54 | 3 |
| 3 | Skap1 | 55 | 4 |
| 4 | Rad51b | 58 | 5 |
| 5 | Pitpnc1 | 58 | 6 |
| 6 | Cenpp | 61 | 7 |
| 7 | Il2ra | 70 | 8 |
| 8 | Taf1d | 71 | 9 |
| 9 | Arhgap15 | 72 | 10 |
| 10 | Hist1h1b | 79 | 11 |
| 11 | Hivep3 | 81 | 12 |
| 12 | Lgals1 | 83 | 13 |
| 13 | Ikzf2 | 91 | 14 |
| 14 | Ccl4 | 105 | 15 |
| 15 | Nop56 | 111 | 16 |
| 16 | Btg1 | 112 | 17 |
| 17 | Knl1 | 122 | 18 |
| 18 | Kif11 | 123 | 19 |
| 19 | Snhg3 | 125 | 20 |

#### Step 1: Initial generation for gene set

##### Process: Proliferative Immune Regulation

This system of interacting proteins appears to be primarily involved in regulating immune cell proliferation, activation, and genomic stability. Below is a critical analysis of the biological processes performed by this system, with reasoning and supporting information for each point.

1. **Cell Proliferation and Mitotic Regulation:** The presence of Mki67 (marker of proliferation Ki-67) and Kif11 (kinesin family member 11) strongly suggests a role in cell division. Mki67 is a well-established marker of cellular proliferation, expressed during all active phases of the cell cycle but absent in quiescent cells. Kif11 is a motor protein essential for spindle formation and chromosome segregation during mitosis. Together, these proteins indicate that the system supports active cell division, likely in rapidly proliferating immune cells such as T or B lymphocytes.
2. **DNA Repair and Genomic Stability:** Rad51b (RAD51 paralog B) is a key player in homologous recombination, a critical DNA repair pathway that ensures genomic stability during cell division. Its inclusion in this system suggests a mechanism to maintain genomic integrity in proliferating immune cells, which is essential to prevent mutations during rapid replication.
3. **Immune Activation and Cytotoxicity:** Gzmb (granzyme B) and Ccl4 (C-C motif chemokine ligand 4) are indicative of immune effector functions. Gzmb is a serine protease involved in inducing apoptosis in target cells during cytotoxic T cell and natural killer (NK) cell responses. Ccl4 is a chemokine that recruits immune

cells to sites of inflammation. These proteins suggest that the system contributes to immune activation and cytotoxic responses.

4. **T Cell Activation and Regulation:** Il2ra (interleukin-2 receptor alpha) and Ikzf2 (IKAROS family zinc finger 2) are critical for T cell activation and differentiation. Il2ra is part of the high-affinity IL-2 receptor complex, which drives T cell proliferation and survival. Ikzf2 is a transcription factor involved in T cell development and regulatory T cell function. These proteins highlight the system's role in modulating T cell responses.
5. **Chromosome Segregation and Kinetochore Function:** Cenpp (centromere protein P) and Knl1 (kinetochore scaffold 1) are essential for proper chromosome segregation during mitosis. Cenpp is part of the CENP-H/I/K complex, which stabilizes kinetochore-microtubule attachments, while Knl1 is a scaffold protein critical for kinetochore assembly and spindle checkpoint signaling. Their presence reinforces the system's role in ensuring accurate cell division.
6. **Ribosome Biogenesis and RNA Processing:** Nop56 (nucleolar protein 56) and Taf1d (TATA-box binding protein associated factor 1D) are involved in ribosome biogenesis and transcriptional regulation. Nop56 is a core component of the box C/D small nucleolar ribonucleoprotein complex, which modifies ribosomal RNA. Taf1d is part of the SL1 complex, which regulates RNA polymerase I transcription. These proteins suggest that the system supports the high biosynthetic demands of proliferating immune cells.
7. **Cytoskeletal Dynamics and Cell Migration:** Skap1 (src kinase-associated phosphoprotein 1) and Arhgap15 (Rho GTPase activating protein 15) are involved in cytoskeletal remodeling and immune cell migration. Skap1 regulates integrin-mediated adhesion and T cell receptor signaling, while Arhgap15 modulates actin cytoskeleton dynamics by inactivating Rac GTPases. These functions are critical for immune cell trafficking and synapse formation.
8. **Chromatin Structure and Gene Regulation:** Hist1h1b (histone H1.5) and Hivep3 (human immunodeficiency virus type I enhancer binding protein 3) are involved in chromatin organization and transcriptional regulation. Hist1h1b is a linker histone that compacts chromatin, while Hivep3 is a transcription factor that regulates immune-related genes. These proteins suggest a role in modulating gene expression during immune responses.
9. **Apoptosis and Cell Cycle Arrest:** Btg1 (B-cell translocation gene 1) is a regulator of cell cycle arrest and apoptosis. It is often expressed in quiescent or differentiating cells, suggesting a role in balancing proliferation and apoptosis in immune cells.
10. **Immune Modulation and Glycan Interactions:** Lgals1 (galectin-1) is a glycan-binding protein that modulates immune responses by inducing apoptosis in activated T cells and promoting immune tolerance. Its presence indicates a role in fine-tuning immune activation and preventing overactivation.
11. **Non-Coding RNA and Gene Regulation:** Snhg3 (small nucleolar RNA host gene 3) is a long non-coding RNA implicated in regulating gene expression and cellular proliferation. Its inclusion suggests additional layers of post-transcriptional regulation in this system.

In summary, this system of interacting proteins is primarily involved in proliferative immune regulation, encompassing cell division, DNA repair, immune activation, and transcriptional control. These processes are critical for maintaining effective and balanced immune responses.

##### Step 2: Self-verification of the process name

**Original_claim:** The entire gene set is involved in Proliferative Immune Regulation. Here is the entire gene set used for verification:

##Mki67,Cmss1,Gzmb,Skap1,Rad51b,Pitpnc1,Cenpp,Il2ra,Taf1d,Arhgap15,Hist1h1b,Hivep3,Lgals1,Ikzf

**Verified_claim:** The claim that the entire gene set is involved in Proliferative Immune Regulation is weakly supported.

Evidence:

1. **Enrichment Analysis:**
   - The gene set is associated with "Integrative Analysis of Treg Glial Interactions" and "Spleen; cells in red pulp," which are related to immune regulation. However, these associations are not directly indicative of proliferative immune regulation.
2. **Pathway Analysis:**
   - Pathways such as "T cell receptor regulation of apoptosis," "Allograft Rejection," and "Interleukin-2 signaling pathway" involve immune regulation and include overlapping genes from the set (e.g., IL2RA, CCL4, GZMB). These pathways suggest immune-related functions but do not explicitly confirm a role in proliferative immune regulation.

While there is evidence of immune-related functions, the specific involvement of the entire gene set in "Proliferative Immune Regulation" is not conclusively demonstrated. The evidence provides weak support for the claim.

**Original_claim:** The gene set plays a role in immune system regulation and proliferation. Here is the entire gene set used for verification:

##Mki67,Cmss1,Gzmb,Skap1,Rad51b,Pitpnc1,Cenpp,Il2ra,Taf1d,Arhgap15,Hist1h1b,Hivep3,Lgals1,Ikzf

**Verified_claim:** The claim that the gene set plays a role in immune system regulation and proliferation is **strongly supported** by the evidence.

1. **Enrichment Analysis:**
   - The gene set is associated with "Integrative Analysis of Treg Glial Interactions," which involves regulatory T cells (Tregs) known for their role in immune system regulation.
   - Another enrichment term, "Spleen; cells in red pulp," is relevant to immune function, as the spleen is a critical organ in the immune system.
2. **Pathway Analysis:**
   - The gene set is linked to pathways such as "T cell receptor regulation of apoptosis," "Allograft Rejection," and "Interleukin-2 signaling pathway," all of which are directly related to immune system regulation and proliferation.

- The "Interleukin-2 signaling pathway" is particularly significant, as IL-2 is a key cytokine in T cell proliferation and immune response.

These findings collectively provide strong evidence that the gene set is involved in immune system regulation and proliferation.

**Original_claim:** The entire gene set contributes to processes related to immune cell proliferation and regulation. Here is the entire gene set used for verification:

##Mki67,Cmss1,Gzmb,Skap1,Rad51b,Pitpnc1,Cenpp,Il2ra,Taf1d,Arhgap15,Hist1h1b,Hivep3,Lgals1,Ikzf2,Ccl4

**Verified_claim:** The claim that the entire gene set contributes to processes related to immune cell proliferation and regulation is supported by evidence.

1. **Enrichment Analysis:**
   - The gene set is significantly associated with "Integrative Analysis of Treg Glial Interactions," which involves regulatory T cells (Tregs) known for their role in immune regulation.
   - Another significant association is with "Spleen; cells in red pulp," which is relevant to immune cell activity in the spleen.
2. **Pathway Analysis:**
   - The gene set overlaps with pathways such as "T cell receptor regulation of apoptosis," "Allograft Rejection," and "Interleukin-2 signaling pathway," all of which are directly related to immune cell regulation and proliferation.
   - Specific genes like IL2RA, CCL4, and GZMB are involved in immune-related processes, further supporting the claim.

Decision: The claim is strongly supported by the provided evidence.

**Original_claim:** Here is the entire gene set used for verification:

##Mki67,Cmss1,Gzmb,Skap1,Rad51b,Pitpnc1,Cenpp,Il2ra,Taf1d,Arhgap15,Hist1h1b,Hivep3,Lgals1,Ikzf2,Ccl4

**Verified_claim:** The gene set provided has been analyzed for enrichment functions, and the following findings are relevant:

1. **Integrative Analysis of Treg Glial Interactions**:
   - This enrichment function is significant with a p-value of 0.0299.
   - It involves interactions between Treg (regulatory T cells) and glial cells.
   - The intersection size is 3, indicating that three genes from the provided set are associated with this function.
2. **Spleen; cells in red pulp**:
   - This enrichment function is significant with a p-value of 0.0499.
   - It is related to cells in the red pulp of the spleen.
   - The intersection size is 9, showing that nine genes from the set are associated with this function.

These findings suggest that the gene set is involved in biological processes related to immune system interactions and spleen cell functions. If the claim is related to these areas, the evidence provides strong support. If the claim is unrelated, the findings may not be directly applicable.

#### Step 3: Modification

**Process: Immune System Regulation and Proliferation**

This system of interacting proteins appears to be primarily involved in regulating immune cell proliferation, activation, and genomic stability. Below is a critical analysis of the biological processes performed by this system, with reasoning and supporting information for each point.

**Cell Proliferation and Mitotic Regulation:** The presence of Mki67 (marker of proliferation Ki-67) and Kif11 (kinesin family member 11) strongly suggests a role in cell division. Mki67 is a well-established marker of cellular proliferation, expressed during all active phases of the cell cycle but absent in quiescent cells. Kif11 is a motor protein essential for spindle formation and chromosome segregation during mitosis. Together, these proteins indicate that the system supports active cell division, likely in rapidly proliferating immune cells such as T or B lymphocytes.

**DNA Repair and Genomic Stability:** Rad51b (RAD51 paralog B) is a key player in homologous recombination, a critical DNA repair pathway that ensures genomic stability during cell division. Its inclusion in this system suggests a mechanism to maintain genomic integrity in proliferating immune cells, which is essential to prevent mutations during rapid replication.

**Immune Activation and Cytotoxicity:** Gzmb (granzyme B) and Ccl4 (C-C motif chemokine ligand 4) are indicative of immune effector functions. Gzmb is a serine protease involved in inducing apoptosis in target cells during cytotoxic T cell and natural killer (NK) cell responses. Ccl4 is a chemokine that recruits immune cells to sites of inflammation. These proteins suggest that the system contributes to immune activation and cytotoxic responses.

**T Cell Activation and Regulation:** Il2ra (interleukin-2 receptor alpha) and Ikzf2 (IKAROS family zinc finger 2) are critical for T cell activation and differentiation. Il2ra is part of the high-affinity IL-2 receptor complex, which drives T cell proliferation and survival. Ikzf2 is a transcription factor involved in T cell development and regulatory T cell function. These proteins highlight the system's role in modulating T cell responses.

**Chromosome Segregation and Kinetochore Function:** Cenpp (centromere protein P) and Knl1 (kinetochore scaffold 1) are essential for proper chromosome segregation during mitosis. Cenpp is part of the CENP-H/I/K complex, which stabilizes kinetochore-microtubule attachments, while Knl1 is a scaffold protein critical for kinetochore assembly and spindle checkpoint signaling. Their presence reinforces the system's role in ensuring accurate cell division.

**Ribosome Biogenesis and RNA Processing:** Nop56 (nucleolar protein 56) and Taf1d (TATA-box binding protein associated factor 1D) are involved in ribosome biogenesis and transcriptional regulation. Nop56 is a core component of the box C/D small nucleolar ribonucleoprotein complex, which modifies ribosomal RNA. Taf1d is part of the SL1 complex, which regulates RNA polymerase I transcription. These proteins suggest that the system supports the high biosynthetic demands of proliferating immune cells.

**Cytoskeletal Dynamics and Cell Migration:** Skap1 (src kinase-associated phosphoprotein 1) and Arhgap15 (Rho GTPase activating protein 15) are involved in cytoskeletal remodeling and immune cell migration. Skap1 regulates integrin-mediated adhesion and T cell receptor signaling, while Arhgap15 modulates actin cytoskeleton dynamics by inactivating Rac GTPases. These functions are critical for immune cell trafficking and synapse formation.

**Chromatin Structure and Gene Regulation:** Hist1h1b (histone H1.5) and Hivep3 (human immunodeficiency virus type I enhancer binding protein 3) are involved in chromatin organization and transcriptional regulation. Hist1h1b is a linker histone that compacts chromatin, while Hivep3 is a transcription factor that regulates immune-related genes. These proteins suggest a role in modulating gene expression during immune responses.

**Apoptosis and Cell Cycle Arrest:** Btg1 (B-cell translocation gene 1) is a regulator of cell cycle arrest and apoptosis. It is often expressed in quiescent or differentiating cells, suggesting a role in balancing proliferation and apoptosis in immune cells.

**Immune Modulation and Glycan Interactions:** Lgals1 (galectin-1) is a glycan-binding protein that modulates immune responses by inducing apoptosis in activated T cells and promoting immune tolerance. Its presence indicates a role in fine-tuning immune activation and preventing overactivation.

**Non-Coding RNA and Gene Regulation:** Snhg3 (small nucleolar RNA host gene 3) is a long non-coding RNA implicated in regulating gene expression and cellular proliferation. Its inclusion suggests additional layers of post-transcriptional regulation in this system.

In summary, this system of interacting proteins is primarily involved in immune system regulation and proliferation, encompassing cell division, DNA repair, immune activation, and transcriptional control. These processes are critical for maintaining effective and balanced immune responses.

#### Step 4: Self-verification of the analytical narratives

**Original_claim:** Mki67 and Kif11 are involved in cell proliferation and mitotic regulation, with Mki67 serving as a marker of cellular proliferation and Kif11 playing a role in spindle formation and chromosome segregation during mitosis.

**Verified_claim:** The claim that Mki67 and Kif11 are involved in cell proliferation and mitotic regulation is supported by the following evidence:

1. **Mki67 (marker of proliferation Ki-67)**:
   - According to the gene summary, Mki67 is described as a "marker of proliferation Ki-67" and is involved in "chromosome segregation and regulation of mitotic nuclear division." This aligns with its role as a marker of cellular proliferation and its involvement in mitotic regulation.
2. **Kif11 (kinesin family member 11)**:
   - The gene summary for Kif11 states that it encodes a motor protein involved in "various kinds of spindle dynamics," including "chromosome positioning, centrosome separation, and establishing a bipolar spindle during cell mitosis." This supports its role in spindle formation and chromosome segregation during mitosis.

Decision: The claim is strongly supported by the evidence provided. Both genes are directly associated with the described functions in cell proliferation and mitotic regulation.

**Original_claim:** Rad51b contributes to DNA repair and genomic stability through its role in homologous recombination, ensuring the integrity of the genome during immune cell division.

**Verified_claim:** The claim that "Rad51b contributes to DNA repair and genomic stability through its role in homologous recombination, ensuring the integrity of the genome during immune cell division" is **strongly supported** by the evidence.

1. **Gene Summary Evidence**:
   - The RAD51B gene encodes a protein that is a member of the RAD51 protein family, which is essential for DNA repair by homologous recombination. RAD51B forms a stable heterodimer with RAD51C and interacts with other family members like RAD51, XRCC2, and XRCC3. This indicates its critical role in DNA repair mechanisms. (Source: Gene summary)
2. **Enrichment Evidence**:
   - RAD51B is part of the "Rad51B-Rad51C-Rad51D-XRCC2 complex," which is directly involved in homologous recombination repair processes. This further supports its role in maintaining genomic stability. (Source: Enrichment data)
3. **Pathway Evidence**:
   - RAD51B is associated with pathways such as "Homologous recombination," "Resolution of D-loop structures through synthesis-dependent strand annealing (SDSA)," and "Defective HDR through homologous recombination due to BRCA1 loss-of-function." These pathways are directly related to DNA repair and genomic stability. (Source: Pathway data)
4. **Immune Cell Division Context**:
   - While the specific role of RAD51B in immune cell division is not explicitly detailed in the provided data, its general function in DNA repair and homologous recombination is critical for maintaining genomic integrity during cell division, which would include immune cells.

The evidence collectively confirms RAD51B's role in DNA repair and genomic stability through homologous recombination, aligning with the claim. However, the specific mention of immune cell division is inferred rather than directly evidenced.

**Original_claim:** Gzmb and Ccl4 are key players in immune activation and cytotoxicity, with Gzmb inducing apoptosis in target cells and Ccl4 recruiting immune cells to sites of inflammation.

**Verified_claim:** The claim that Gzmb and Ccl4 are key players in immune activation and cytotoxicity, with Gzmb inducing apoptosis in target cells and Ccl4 recruiting immune cells to sites of inflammation, is strongly supported by evidence.

1. **Gzmb (Granzyme B)**:

- Summary information indicates that Gzmb is secreted by natural killer (NK) cells and cytotoxic T lymphocytes (CTLs). It is an active protease that induces apoptosis in target cells. Additionally, it processes cytokines and degrades extracellular matrix proteins, implicating roles in chronic inflammation and wound healing.
- Enrichment analysis shows that Gzmb is involved in pathways such as "Interactions of natural killer cells in pancreatic cancer" and "miRNA role in immune response in sepsis," which are related to immune activation and cytotoxicity.

2. **Ccl4 (C-C motif chemokine ligand 4)**:
   - Summary information reveals that Ccl4 is a secreted protein with chemokinetic and inflammatory functions. It is one of the major HIV-suppressive factors produced by CD8+ T-cells, highlighting its role in immune cell recruitment and inflammation.
   - Enrichment analysis further supports its involvement in immune-related pathways, such as "miRNA role in immune response in sepsis."

These findings align with the claim, confirming the roles of Gzmb and Ccl4 in immune activation and cytotoxicity.

**Original_claim:** Il2ra and Ikzf2 are critical for T cell activation and regulation, with Il2ra driving T cell proliferation and survival, and Ikzf2 contributing to T cell development and regulatory T cell function.

**Verified_claim:** The claim that Il2ra and Ikzf2 are critical for T cell activation and regulation, with Il2ra driving T cell proliferation and survival, and Ikzf2 contributing to T cell development and regulatory T cell function, is supported by evidence but with some ambiguity.

1. **Il2ra (Interleukin 2 receptor subunit alpha)**:
   - Summary: Il2ra is part of the high-affinity IL2 receptor complex, which plays a role in immune responses. It is involved in T cell activation and proliferation. Elevated levels of IL2R are associated with immune-related conditions, including severe COVID-19 and carcinomas. This supports the claim that Il2ra drives T cell proliferation and survival.
   - Enrichment: Il2ra is associated with "Abnormal CD4+ T cell subset proportion" and "Abnormal CD4:CD8 ratio," which are relevant to T cell regulation.
2. **Ikzf2 (IKAROS family zinc finger 2)**:
   - Summary: Ikzf2 is a transcription factor involved in lymphocyte development, particularly in early hematopoietic development. It forms dimers with other Ikaros family members and is implicated in regulatory T cell function. This aligns with the claim that Ikzf2 contributes to T cell development and regulatory T cell function.
   - Enrichment: Ikzf2 is linked to "Integrative Analysis of Treg Glial Interactions," which suggests a role in regulatory T cell function.

While the evidence supports the general roles of Il2ra and Ikzf2 in T cell activation, proliferation, development, and regulation, the specific mechanisms described in the claim (e.g., driving proliferation and survival for Il2ra, and contributing to regulatory T cell function for Ikzf2) are not explicitly detailed in the provided data. Therefore, the support for the claim is strong but not definitive.

**Original_claim:** Cenpp and Knl1 are essential for chromosome segregation and kinetochore function, with Cenpp stabilizing kinetochore-microtubule attachments and Knl1 supporting kinetochore assembly and spindle checkpoint signaling.

**Verified_claim:** The claim that Cenpp and Knl1 are essential for chromosome segregation and kinetochore function is strongly supported by the evidence.

1. **Cenpp**:
    - According to the gene summary, CENPP is a subunit of a centromeric complex that targets CENPA to centromeres and is required for proper kinetochore function and mitotic progression. This aligns with the claim that Cenpp stabilizes kinetochore-microtubule attachments.
2. **Knl1**:
    - The gene summary for KNL1 states that it is a component of the multiprotein assembly required for kinetochore-microtubule attachments and chromosome segregation. It functions as a scaffold for proteins influencing the spindle assembly checkpoint during the eukaryotic cell cycle. This supports the claim that Knl1 supports kinetochore assembly and spindle checkpoint signaling.

Both gene summaries provide direct evidence for the roles of Cenpp and Knl1 in chromosome segregation and kinetochore function, confirming the claim.

**Original_claim:** Nop56 and Taf1d are involved in ribosome biogenesis and RNA processing, with Nop56 modifying ribosomal RNA and Taf1d regulating RNA polymerase I transcription to meet the biosynthetic demands of immune cells.

**Verified_claim:** The claim that Nop56 and Taf1d are involved in ribosome biogenesis and RNA processing is supported by evidence, but the specific assertion about their roles in immune cells is not directly confirmed.

1. **Nop56**:
    - Summary: Nop56 is a nucleolar protein involved in the assembly of the 60S ribosomal subunit and pre-rRNA processing. It is part of a complex with nucleolar proteins Nop58 and fibrillarin, which are essential for ribosome biogenesis.
    - Evidence: The gene summary explicitly states its role in ribosome assembly and RNA processing.
2. **Taf1d**:
    - Summary: Taf1d is a member of the SL1 complex, which plays a role in RNA polymerase I transcription. RNA polymerase I is responsible for transcribing ribosomal RNA (rRNA), which is crucial for ribosome biogenesis.

- Evidence: The gene summary confirms its involvement in RNA polymerase I transcription, which aligns with the claim about RNA processing and ribosome biogenesis.

However, the claim that Nop56 modifies ribosomal RNA and Taf1d regulates RNA polymerase I transcription specifically to meet the biosynthetic demands of immune cells is not directly supported by the provided evidence. The summaries do not mention immune cell-specific functions for these genes. Therefore, while their general roles in ribosome biogenesis and RNA processing are confirmed, the immune cell-specific aspect remains unverified.

**Original_claim:** Skap1 and Arhgap15 regulate cytoskeletal dynamics and cell migration, with Skap1 modulating integrin-mediated adhesion and T cell receptor signaling, and Arhgap15 inactivating Rac GTPases to control actin cytoskeleton remodeling.

**Verified_claim:** The claim that Skap1 and Arhgap15 regulate cytoskeletal dynamics and cell migration, with Skap1 modulating integrin-mediated adhesion and T cell receptor signaling, and Arhgap15 inactivating Rac GTPases to control actin cytoskeleton remodeling is **strongly supported** based on the following evidence:

1. **Skap1**:
   - The gene summary for Skap1 indicates that it encodes a T cell adaptor protein involved in inside-out signaling by coupling T-cell antigen receptor stimulation to the activation of integrins. This supports the claim that Skap1 modulates integrin-mediated adhesion and T cell receptor signaling.
2. **Arhgap15**:
   - The gene summary for Arhgap15 states that it is a Rho GTPase-activating protein (GAP) that regulates the activity of RHO GTPases, which are known to control diverse biological processes, including actin cytoskeleton remodeling. This aligns with the claim that Arhgap15 inactivates Rac GTPases to control actin cytoskeleton remodeling.

No additional enrichment data was retrieved for the gene set, but the gene summaries provide direct evidence supporting the claim.

**Original_claim:** Hist1h1b and Hivep3 contribute to chromatin structure and gene regulation, with Hist1h1b compacting chromatin and Hivep3 regulating immune-related gene expression.

**Verified_claim:** The claim that "Hist1h1b and Hivep3 contribute to chromatin structure and gene regulation, with Hist1h1b compacting chromatin and Hivep3 regulating immune-related gene expression" is **supported** by the following evidence:

1. **Hist1h1b (H1-5)**:
   - According to the gene summary, Hist1h1b encodes a linker histone (H1) that interacts with linker DNA between nucleosomes. It plays a role in the compaction of chromatin into higher-order structures. This directly supports the claim that Hist1h1b contributes to chromatin structure by compacting chromatin.

2. **Hivep3**:

- The gene summary for Hivep3 indicates that it encodes a transcription factor that regulates nuclear factor kappaB-mediated transcription by binding the kappaB motif in target genes. It also binds recombination signal sequences flanking immunoglobulin and T-cell receptor regions. This evidence supports the claim that Hivep3 is involved in regulating immune-related gene expression.

Both components of the claim are substantiated by the provided gene summaries.

**Original_claim:** Btg1 plays a role in apoptosis and cell cycle arrest, balancing proliferation and differentiation in immune cells.

**Verified_claim:** The claim that Btg1 plays a role in apoptosis and cell cycle arrest, balancing proliferation and differentiation in immune cells is strongly supported.

Evidence:

1. Gene Summary: BTG1 is described as a member of an anti-proliferative gene family that regulates cell growth and differentiation. Its expression is highest in the G0/G1 phases of the cell cycle and is downregulated as cells progress through G1. This indicates its involvement in cell cycle regulation and potentially in cell cycle arrest.
2. Functional Role: The encoded protein interacts with several nuclear receptors and functions as a coactivator of cell differentiation. This supports the idea that BTG1 balances proliferation and differentiation.
3. Disease Association: BTG1 has been implicated in chromosomal translocation in a case of B-cell chronic lymphocytic leukemia, which may suggest its relevance in immune cells.

These points collectively provide strong evidence for the claim.

**Original_claim:** Lgals1 modulates immune responses by inducing apoptosis in activated T cells and promoting immune tolerance through glycan interactions.

**Verified_claim:** The claim that "Lgals1 modulates immune responses by inducing apoptosis in activated T cells and promoting immune tolerance through glycan interactions" is supported by evidence, but the association is not entirely direct and clear.

1. The gene summary for LGALS1 (galectin-1) indicates that it belongs to the family of beta-galactoside-binding proteins, which are implicated in modulating cell-cell and cell-matrix interactions. This aligns with the claim that LGALS1 interacts with glycans, as beta-galactoside-binding proteins are known for such interactions.
2. Enrichment analysis for LGALS1 reveals its involvement in the "galectin complex," described as a homodimeric protein complex capable of binding carbohydrates and involved in anti-inflammatory and pro-apoptotic processes. This supports the claim that LGALS1 induces apoptosis and plays a role in immune modulation.
3. No direct disease associations or biological domains were retrieved for LGALS1, which limits the ability to confirm its role in promoting immune tolerance specifically.

Decision: The claim is weakly supported. While there is evidence that LGALS1 is involved in glycan interactions and apoptosis, the specific mechanisms of immune tolerance and its role in activated T cells require further direct evidence for stronger validation.

**Original_claim:** Snhg3, a long non-coding RNA, is implicated in regulating gene expression and cellular proliferation, adding layers of post-transcriptional regulation to immune system processes.

**Verified_claim:** The claim that Snhg3, a long non-coding RNA, is implicated in regulating gene expression and cellular proliferation, adding layers of post-transcriptional regulation to immune system processes, has weak support based on the following evidence:

1. Gene Summary: Snhg3 (small nucleolar RNA host gene 3) is predicted to be active in the nucleus. However, there is no direct mention in the summary of its role in regulating gene expression, cellular proliferation, or immune system processes. This provides limited evidence for the claim.
2. Biological Domains: No specific biological domains were identified for Snhg3, which limits the ability to confirm its involvement in post-transcriptional regulation or immune system processes.
3. Enrichment Functions: No enrichment functions were identified for Snhg3, which further weakens the evidence for its role in regulating gene expression or immune system processes.

While there is some indirect evidence that Snhg3 is active in the nucleus, which is consistent with a potential regulatory role, there is no direct evidence from the tools used to confirm its specific involvement in the processes mentioned in the claim.

**Original_claim:**

**Verified_claim:** Failed.

#### Step 5: Summarization

##### Process: Immune System Regulation and Proliferation

This system of interacting proteins is involved in immune system regulation and proliferation, encompassing cell division, DNA repair, immune activation, and transcriptional control. Below is a critical analysis of the biological processes performed by this system, with reasoning and supporting information for each point.

Mki67 and Kif11 are involved in cell proliferation and mitotic regulation. Mki67 serves as a marker of cellular proliferation and is involved in chromosome segregation and regulation of mitotic nuclear division. Kif11 encodes a motor protein essential for spindle dynamics, including chromosome positioning, centrosome separation, and establishing a bipolar spindle during mitosis.

Rad51b contributes to DNA repair and genomic stability through its role in homologous recombination. It forms a stable complex with other RAD51 family members, ensuring the integrity of the genome during cell division.

Gzmb and Ccl4 are key players in immune activation and cytotoxicity. Gzmb induces apoptosis in target cells and processes cytokines, while Ccl4 recruits immune cells to sites of inflammation and acts as a major HIV-suppressive factor produced by CD8+ T-cells.

Il2ra and Ikzf2 are critical for T cell activation and regulation. Il2ra is part of the high-affinity IL-2 receptor complex, driving T cell proliferation and survival. Ikzf2 is a transcription factor involved in lymphocyte development and regulatory T cell function.

Cenpp and Knl1 are essential for chromosome segregation and kinetochore function. Cenpp stabilizes kinetochore-microtubule attachments, while Knl1 supports kinetochore assembly and spindle checkpoint signaling.

Nop56 and Taf1d are involved in ribosome biogenesis and RNA processing. Nop56 is a nucleolar protein essential for ribosome assembly and pre-rRNA processing. Taf1d is part of the SL1 complex, regulating RNA polymerase I transcription for ribosomal RNA synthesis.

Skap1 and Arhgap15 regulate cytoskeletal dynamics and cell migration. Skap1 modulates integrin-mediated adhesion and T cell receptor signaling, while Arhgap15 inactivates Rac GTPases to control actin cytoskeleton remodeling.

Hist1h1b and Hivep3 contribute to chromatin structure and gene regulation. Hist1h1b compacts chromatin into higher-order structures, while Hivep3 regulates immune-related gene expression by binding kappaB motifs and recombination signal sequences.

Btg1 plays a role in apoptosis and cell cycle arrest, balancing proliferation and differentiation in immune cells. It is highly expressed during the G0/G1 phases of the cell cycle and interacts with nuclear receptors to regulate cell differentiation.

Lgals1 modulates immune responses by inducing apoptosis in activated T cells and interacting with glycans to influence cell-cell and cell-matrix interactions.

Snhg3, a long non-coding RNA, is active in the nucleus but lacks direct evidence for its role in regulating gene expression or immune system processes. Its function remains unclear in this context.

In summary, this system of interacting proteins is primarily involved in immune system regulation and proliferation, supporting critical processes such as cell division, DNA repair, immune activation, and transcriptional control.
